# Facing the facts: Adaptive trade-offs along body size ranges determine mammalian craniofacial scaling

**DOI:** 10.1101/2023.09.28.560051

**Authors:** D. Rex Mitchell, Emma Sherratt, Vera Weisbecker

## Abstract

The mammalian cranium (skull without lower jaw) is representative of mammalian diversity and is thus of particular interest to mammalian biologists across disciplines. One widely retrieved pattern accompanying mammalian cranial diversification is referred to as “craniofacial evolutionary allometry” (CREA). This posits that “adults of larger species, in a group of closely related mammals, tend to have relatively longer faces and smaller braincases”. However, no process has been officially suggested to explain this pattern, there are many exceptions, and its predictions potentially conflict with well-established biomechanical principles. Understanding the mechanisms behind CREA and causes for deviations from the pattern therefore has tremendous potential to explain allometry and diversification of the mammalian cranium. Here, we propose an amended framework to characterise the CREA pattern more clearly, in that “longer faces” can arise through several kinds of evolutionary change, including elongation of the rostrum, retraction of the jaw muscles, or a more narrow or shallow skull, which all result in a generalised gracilisation of the facial skeleton with increased size. We define a standardised workflow to test for the presence of the pattern, using allometric shape predictions derived from geometric morphometrics analysis, and apply this to 22 mammalian families including marsupials, rabbits, rodents, bats, carnivores, antelope, and whales. Our results show that increasing facial gracility with size is common, but not necessarily as ubiquitous as previously suggested. To address the mechanistic basis for this variation, we then review cranial adaptations for harder biting. These dictate that a more gracile cranium in larger species must represent a sacrifice in the ability to produce or withstand harder bites, relative to size. This leads us to propose that facial gracilisation in larger species is often a product of bite force allometry and phylogenetic niche conservatism, where more closely related species tend to exhibit more similar feeding ecology and biting behaviours and, therefore, absolute (size-independent) bite force requirements. Since larger species can produce the same absolute bite forces as smaller species with less effort, we propose that relaxed bite force demands can permit facial gracility in response to bone optimisation and alternative selection pressures. Thus, mammalian facial scaling represents an adaptive by-product of the shifting importance of selective pressures occurring with increased size. A reverse pattern of facial “shortening” can accordingly also be found, and is retrieved in several cases here, where larger species incorporate novel feeding behaviours involving greater bite forces. We discuss multiple exceptions to a bite force-mediated influence on facial length across mammals which lead us to argue that ecomorphological specialisation of the cranium is likely to be the primary driver of facial scaling patterns, with developmental and/or phylogenetic constraints a secondary factor. A potential for larger species to have a wider range of cranial functions when less constrained by biomechanical demands might also explain why selection for larger sizes seems to be prevalent in some mammalian clades. The interplay between adaptation and constraint across size ranges thus presents an interesting consideration for a mechanistically grounded investigation of mammalian cranial allometry.

## I INTRODUCTION

### (1) Patterns versus processes in mammalian cranial evolution

Mammals display a staggering degree of diversity in form. The class spans eight orders of magnitude in body mass (Smith & Lyons, 2011) and representatives live in most environments on Earth, through adaptations to various abiotic and biotic factors. The patterns behind the diversity of mammalian morphology have been a core interest of Western science for centuries (Aristotle, ∼450; Linnaeus, 1758). Of particular interest for most of this venerable history has been the evolution of the mammalian cranium (the skull without the lower jaw), which reflects the diversity of mammals like no other part of the skeleton (Novacek, 1993). Crania incorporate the brain and the majority of an animals’ sensory apparatuses, such as the eyes, nose, and ears. Perhaps more importantly, their role in the ingestion of food makes the functional adaptation of the cranium a matter of day-to-day survival, so that cranial morphology offers insights into a given mammal’s feeding ecology and behaviour. The patterns of mammalian cranial shape evolution have thus become a shorthand for understanding how mammalian diversity has evolved. Recent advances in technology, such as digital-based data collection and their applications in quantitative analysis, have revolutionised our capacity to characterise mammalian cranial diversity patterns. As our understanding of these patterns expands in a contemporary setting, we also require new theoretical frameworks to identify the underlying evolutionary processes.

A particularly promising opportunity to link a widespread pattern of morphological variation with evolutionary process comes from the study of cranial allometry, or how mammalian crania vary in shape relative to size. In particular, over the last decade, a trend of increased face length with increasing body size has been repeatedly found. This pattern, termed “craniofacial evolutionary allometry” or CREA (Cardini et al., 2015), is potentially the most widely-retrieved and investigated evolutionary pattern of the mammalian cranium. However, the mechanistic basis for this frequent phenomenon is unclear and requires reconciliation in biomechanical terms. For example, CREA predicts that facial skeletons of larger mammal species are more elongated than those of smaller relatives (Cardini & Polly, 2013; Cardini, 2019). However, elongate faces also confer a weaker bite, so that mammals capable of harder biting are predicted to have stouter cranial proportions (Greaves, 1985; Goswami et al., 2010). What therefore happens when a larger species must also bite harder to exploit the full dietary ranges of their niches? And how do constraints imparted by feeding biomechanics influence scaling patterns across cranial size ranges in general? The fact that cranial shape is so heavily aligned with its biomechanical function also means that the appearance of a CREA pattern might relate to different processes depending on the feeding ecology of the taxonomic group investigated. It is therefore unlikely that the CREA pattern is due to a single blanket mechanism, and that substantial nuance is required to explain its occurrence.

### (2) The “rule” of Craniofacial Evolutionary Allometry: Definition and applicability

CREA describes a tendency for larger species to have longer faces than closely related, smaller species (Cardini & Polly, 2013; Cardini, 2019). Over the last decade, CREA has been proposed as a ubiquitous feature of interspecific mammalian (and possibly even vertebrate) cranial evolutionary allometry (Cardini & Polly, 2013, Cardini et al., 2015; Bright et al., 2016; Linde-Medina, 2016; Tamagnini et al., 2017; Cardini, 2019; Gómez & Lois-Milevicich, 2021). The pattern was initially noted by several authors (Robb, 1935; Radinsky, 1985) and recorded in a formal context across a diverse range of mammalian groups, including fruit bats, mongooses, squirrels, antelope, cats, kangaroos, and particularly clear evidence for the pattern has been found in rodents and bovids across multiple studies (Cardini & Polly, 2013; Magnus et al., 2018; Marcy et al., 2020; Bibi & Tyler, 2022; Rhoda et al., 2022). However, CREA does not appear to be universal. Several studies have since noted a range of cases that can seemingly contradict the predictions of CREA (Hautier et al., 2014; De Muizon et al., 2015; Flores et al., 2018; Law et al., 2018; Mitchell et al., 2018). Furthermore, there are many examples of small mammals with relatively elongate faces, from marsupial honey possums (*Tarsipes rostratus*) and long-nosed potoroos (*Potorous tridactylus*) to placental elephant shrews (Macroscelidea) and nectar-feeding phyllostomid bats (Rosenberg & Richardson, 1995; Panchetti et al., 2008; Mitchell et al., 2018; Rossoni et al., 2019), which appear to rule out a short face as a requisite trait of small body size. Likewise, larger mammals with relatively shorter faces, such as sea otters among mustelids, some primate species including humans, and extinct short-faced kangaroos (Radinsky, 1981; Prideaux, 2004; Fleagle et al., 2010; Mitchell, 2019) appear to reflect a pattern reverse to CREA, thereby questioning its universal applicability for all mammalian groups.

Together, the above examples demonstrate that the CREA pattern does not arise from any fundamental constraints on the potential for elongation of the face. Rather, the aforementioned conflicting accounts suggest that there might be more than one pattern of mammalian facial scaling. This impression is supported when we review the interpretations of the presence of CREA, showing that many researchers, including ourselves, have addressed the existence of facial elongation in their datasets under widely varied definitions. Originally, it was defined quite simply by its original author (Cardini & Polly, 2013; Cardini, 2019): “Adults of larger species, in a group of closely related mammals, tend to have relatively longer faces and smaller braincases”. However, of 50 publications that cite the original publication on CREA by Cardini & Polly (2013) in reference to observed patterns of craniofacial evolutionary allometry at the time of writing, we identify a range of different secondary interpretations (Apprendix A: Table S1), with varying alignment to the initial definition. We note that these sentences are only a very small part of considerably more complex research papers; however, these diverse interpretations reflect mounting evidence that facial elongation on a macroevolutionary scale is likely not achieved by a universal pattern, and that a longer face can be interpreted in a multitude of ways under the current definition. A clear understanding of the processes at play is further obscured because a precise mechanism for why CREA occurs remains elusive (Cardini, 2019). Here, we argue that a mechanistic understanding of mammalian facial allometry is required to clarify the definition of the pattern and find its place in the collective understanding of mammalian cranial diversity. This journey starts with the question of how allometry is understood more broadly, which we cover in the following section.

### (3) The three levels of allometry: constraint *versus* adaptation

Allometry is defined as the size-related changes of morphological traits (Huxley, 1932; Huxley & Tessier, 1936; Gould, 1966; Klingenberg, 1996; 2016). While the physical traits of a larger species will exhibit increased *absolute* size compared to homologous features in smaller species, allometry is related to *relative* size changes. If all features were to become larger at the same rate, this would be isometric enlargement. However, if a given trait is larger than expected by isometry, this is positive allometry (hyperallometric); and if it is smaller than expected, it is negatively allometric (hypoallometric) (Huxley, 1932; Rensch, 1948). In the context of skeletal shape, allometry is typically researched and discussed at three main levels (Fig. 1): *ontogenetic allometry* concerns changes to the size proportions of physical traits that occur throughout the growth and development of an individual; *static allometry* refers to covariation in size or trait proportions for individuals of the same species at a similar age or developmental stage; and *evolutionary allometry* involves interspecific trends in size-correlated changes to trait proportions (Cock, 1966; Cheverud, 1982; Hallgrímsson, et al., 2019).

**Figure 1:**
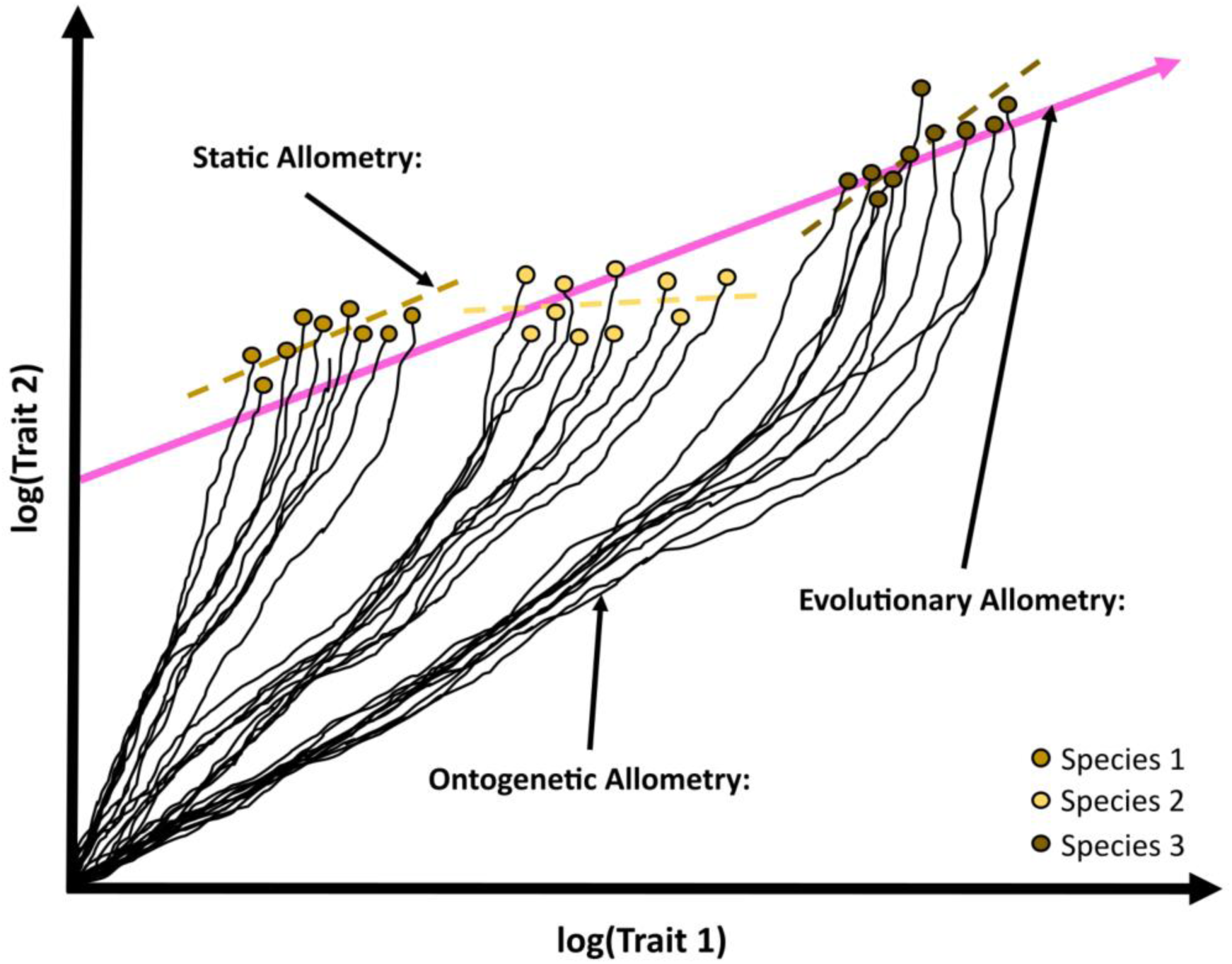
The three levels of allometry, modified from Neiro (2022). Ontogenetic allometry indicating size-related differences in trait proportions for each individual (black lines); static allometry (dashed lines) across individuals of each species at a given age or developmental stage; and evolutionary allometry (solid pink line) showing shape relationships across all individuals of all species at a specific age or developmental stage. Note that ontogenetic allometry is not always log-linear, as demonstrated by species 3 (Pélabon et al., 2013).

Ontogenetic, static, and evolutionary allometry are each important aspects of assessing evolutionary patterns of allometry. However, while the three types are often correlated (Pélabon, et al., 2013), they are not necessarily equivalent, nor always simple to distinguish from one another. For example, even though individuals of a species share similar overall ontogenetic allometry, many aspects of their ontogeny can be influenced by variation in developmental genetics and differences in their environment (e.g., food types or nutrition levels). This variation can cause the species’ ontogenetic allometry to vary among individuals and static allometry to deviate from the ontogenetic pattern (Lande, 1979; Pélabon, et al., 2013; Voje et al., 2013; Hallgrímsson et al., 2019). Similarly, differences in the functional relationships between traits and body size across species can change the evolution of ontogenetic allometries, and thus result in interspecific differences in both ontogenetic and static allometries (Rensch, 1948). This can cause evolutionary allometry to deviate from static allometry (Voje, et al., 2013). For evolutionary allometry to emerge from static allometry, the developmental processes leading to static allometry need to be fixed across species, or a consistent allometric relationship between shape and size must be selected for through evolutionary time (Gould 1975; Lande 1979; Voje et al., 2013). Given these relationships, it is not always easy to distinguish where one kind of allometry ends and another begins. The difficulty of disentangling how the levels of allometry relate to each other is particularly obvious regarding the generation of hypotheses to explain evolutionary allometry in the mammalian cranium.

Hypotheses regarding evolutionary allometry tend to be either defined as a product of developmental constraints on trait evolution, or functional adaptation. With respect to CREA, patterns of relatively longer faces in larger-sized skulls have been explained by the hypoallometric scaling of the brain (and therefore the brain case) with body size (Radinsky, 1984a), which would support a constraint-based hypothesis. Similarly, both interspecific differences in face length and hyperallometry of face length in mammals has been suggested to relate to heterochrony, or size-related constraints on heterochrony, respectively (Cardini & Polly, 2013; Usui & Tokita, 2018, and see Bhullar et al., 2012 for a similar argument in birds). However, this perspective leaves little room for an adaptive component, and requires a strict form of the allometric-constraint hypothesis, whereby evolutionary changes must lack evolvability and follow trajectories imposed solely by ontogenetic or static allometries (Voje et al., 2013). Because these hypotheses do not consider the form-function relationship in the living animal, which is ultimately what is selected for, a constraints-focused explanatory approach has the potential to overlook size-related ecomorphological specialisation (Pyron & Burbrink, 2009). Approaching from a functional perspective, convergence of size-related specialisations can result in similar evolutionary allometries in disparate groups of mammals (Zelditch & Swiderski, 2022). If so, the repeated cases of facial elongation across mammalian clades would be a product of size-correlated adaptations to maintain a base-level of equivalent function across size ranges (von Bertalanffy & Pirozynski, 1952; Emerson & Bramble, 1993; Zelditch & Swiderski, 2022). Furthermore, dietary pressures and biomechanical adaptations have been suggested as potential explanations for the pattern (Cardini, 2019). Defining mechanisms behind allometric patterns in cranial evolution, and teasing apart the processes involved, therefore requires insights into the degree to which constraint and adaptation contribute to the observed pattern of facial elongation with increased body size.

### (4) Aims and scope

This review aims to provide a framework to help differentiate the extent to which functional adaptation and developmental constraint produce the frequently observed pattern of facial elongation in larger mammalian species first raised by CREA. Using the ambiguity regarding what constitutes the CREA pattern as a starting point, we define a repeatable, testable hypothesis for facial allometry across any taxonomic group of animal crania. We then use our standardised testing to compare cranial evolutionary allometry across 22 mammalian families, encompassing more than 550 species of marsupials, rabbits, rodents, bats, carnivores, antelope, and whales. Based on these results, we then produce the first mechanistic framework that can broadly explain the presence of allometric facial elongation across disparate clades. Finally, we discuss some limitations and exceptions to the framework, and suggest how these patterns all fit into our collective understanding of the role of evolutionary allometry in ecological specialisation and morphological diversification.

## II HOW DO WE TEST FOR CRANIOFACIAL ALLOMETRY?

Cranial allometric variation in mammals is often related to changes in the facial dimensions relative to the braincase (Radinsky, 1984a, 1985; Emerson & Bramble, 1993; Zelditch & Swiderski, 2022). CREA captures this by noting that, “Adults of larger species, in a group of closely related mammals, tend to have relatively longer faces and smaller braincases” (Cardini, 2019). However, for this definition to be translated into testable scenarios, several aspects of it require clarification: (1) what is “closely related”, and how is relatedness considered in testing? (2) what is a “relatively longer face”? and (3) are we seeking evidence of just one allometric pattern or a combination of several partially or entirely independent patterns? The following discusses these concepts, which we then use to put forward a simple methodological framework that accommodates them, in order to test for allometric facial elongation across species.

### (1) What is “closely related” and how is relatedness considered in testing?

The definition of “closely related” postulated as part of CREA requires careful attention, because identifying the evolutionary levels and times of divergence across which we expect allometric regularities is an important part of analysing them. Cardini (2019) discussed the difficulties of defining a taxonomic scale for which to test CREA, and the initial studies of CREA are a case in point because taxa that have been tested in these studies range from Order (Cingulata) to Genus (*Equus*), with divergence times spanning an estimated ∼46 million years ago (mya) (family: Leporidae) to less than 3mya (genus: *Equus*) (Fig. 2). However, most tested taxa tend to be at the level of family to tribe, with divergence times of 30mya-6mya. The influence of taxonomic scale on patterns of craniofacial allometry has not been explicitly tested; however, some subsets within clades have been assessed in isolation after initial analysis, all of which have yielded results congruent with the broader taxon under study (Cardini et al., 2015; Tamagnini et al., 2017).

**Figure 2:**
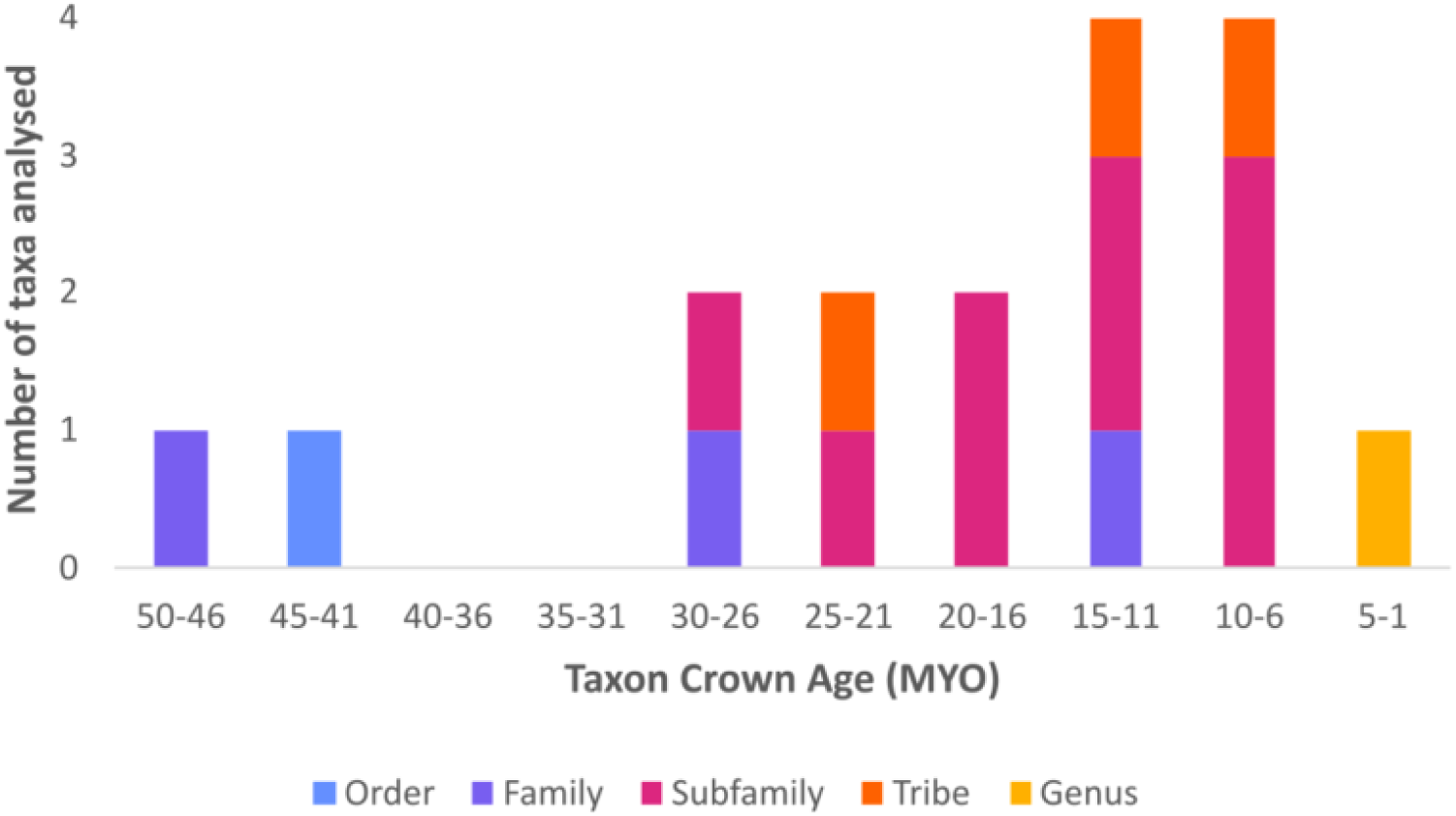
The divergence times and taxonomic scale of clades tested in key works on CREA (Cardini & Polly, 2013; Cardini et al., 2015; Tamagnini et al., 2017; Cardini, 2019). Clade age given as millions of years ago (mya), see Appendix A for sources.

Another aspect to consider is how relatedness is factored into allometry testing, and how this might influence results. If the degree of relatedness is defined by phylogenetic proximity, there may be a problem with applying phylogenetic comparative methods when size is correlated with phylogeny. For example, Cope’s rule states that there can be a tendency of larger body sizes to evolve over evolutionary time (Cope, 1896; Benton, 2002); and this is particularly prevalent in mammals (Alroy, 1998). How potential correlations between relatedness and body size might impact analyses and interpretations of cranial evolutionary allometry therefore needs addressing as well, because accounting for phylogenetic relatedness in statistical analysis has the potential to obscure size-correlated shape changes.

### (2) What is a “relatively longer face”?

Relative face length is a subjective concept, because the term “longer” is a question of proportions. For example, a longer face might be a product of a longer rostrum (projected maxillae/premaxillae/nasal bones); more posterior positioning of the zygomatic arches and jaw muscles; or just an overall contrast between stout (wider/deeper/more sturdy) and gracile (narrower/flatter/more slender) shape. In other words, a relatively shallower or narrower face might also be interpreted as having a longer face when rostrum (or muzzle/snout) length, in fact, remains consistent. This is similar to Figueirido et al. (2010a)’s finding that the extinct giant short-faced bear (*Arctodus simus*) has a similar rostrum length to other bears, but the appearance of a short face in this species is an optical illusion due to having a much deeper skull. Some research has noted a relatively thinner or shallower face within the context of CREA (Tamagnini et al., 2017; Magnus et al., 2018; Weisbecker et al., 2019; Hennekam et al., 2020; van der Geer, 2020), and the CREA pattern itself has occasionally been defined as including relative skull flatness (Zelditch & Swiderski, 2022), braincase width (Linde-Medina, 2016), or width of the zygomatic arches (Weisbecker et al., 2019) alongside shifts in face length (Appendix A; Table S1). None of these interpretations are necessarily inaccurate, but they highlight the need for a more consistent definition to allow an appropriate interpretation of craniofacial allometric patterns. This requires consideration of whether a “longer” face must involve elongation of the rostrum, as defined in a large proportion of studies that reference CREA (Appendix A: Table S1) or can be accepted as present simply due to increased gracility of the cranium.

The definition of what constitutes the “face” also represents a substantial challenge to understanding craniofacial allometry, as reflected in the diverse interpretations of the limits of the “face” relative to the remainder of the skull. For example, ∼20% of studies define CREA specifically as involving a longer rostrum (Appendix A: Table S1), with some studies mentioning both rostrum and face interchangeably. It would seem that many researchers therefore define face length as rostrum length. The rostrum is typically defined as the part of the cranium that is anterior to the orbits (Devillers et al., 1984; Van Valkenburgh et al., 2014), and this region of the cranium is usually delineated by the nasofrontal suture and anterior orbits (e.g., Cardini & Polly, 2013; Mitchell et al., 2018; Marcus et al., 2000). However, it has not been acknowledged that relative orbit size itself can influence the appearance of a longer rostrum in larger species. In mammals with a body mass above ∼1kg, eye size is hypoallometric (Hughes, 1977; Howland et al., 2004), with an eye of 35-50 mm diameter meeting the needs of most mammalian species over ∼500kg in body mass (Hughes, 1977). Shifts in eye size are evidenced in the size of the bony orbits, such that relatively smaller eyes in larger species are accompanied by relatively smaller orbits (e.g., Radinsky, 1984b; 1985; Emerson & Bramble, 1993; Kitchener et al., 2010; Debey et al., 2013; Singleton, 2013; Krone et al., 2019). Figure 3 demonstrates how, in smaller mammals, the orbit is positioned more anterior relative to the cheek tooth row. Relatively smaller orbits in larger mammals can therefore potentially be interpreted as rostrum elongation with increased body size, despite being due to relative orbit size. This trade-off between orbit size and rostrum length is similar, in principle, to the theoretical morphospace of diapsid skulls, which predicts that a proportional increase in orbit size must result in a corresponding proportional decrease in rostrum length (Marugán-Lobón & Buscalioni, 2003). Orbit size could be influencing interpretations of facial elongation; however, as previously discussed, “face length” is also a question of skull proportions, which orbit size has little influence over. Furthermore, the face has been measured from the ventral aspect to include regions as far posterior as the rear palate (Wayne, 1986; Cardini & Polly, 2013; Cardini, 2019) or rear tooth row (Radinsky, 1985; Hallgrímsson et al., 2006). These regions fall behind the orbits in many (but not all) species, suggesting that the “face” should include more than just the rostrum anterior to the orbits. Facial elongation might therefore be better defined in the absence of this confounding issue of orbit size, and instead via investigation of overall craniofacial proportions.

**Figure 3:**
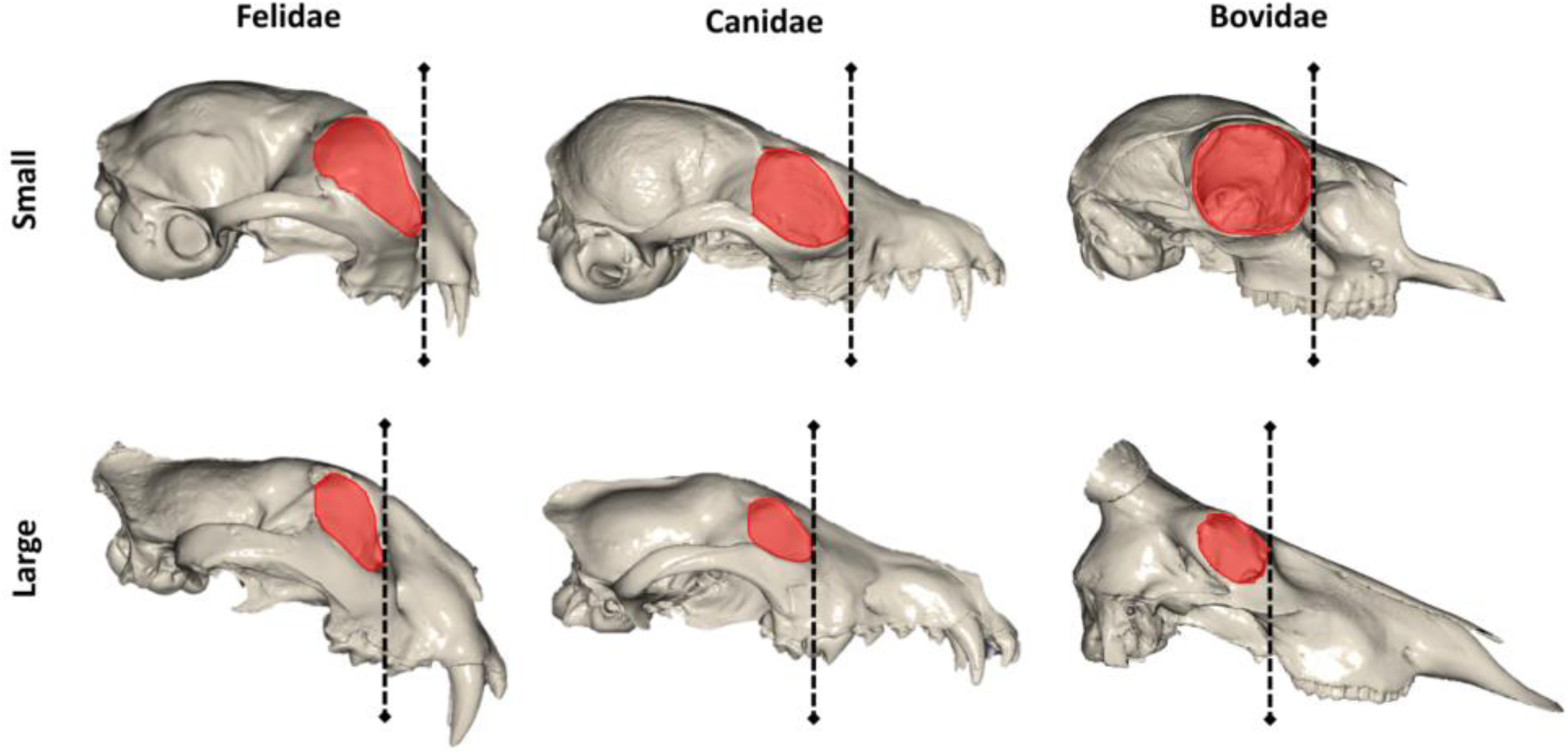
Larger orbits in smaller species extend further forward on the cranium, possibly enhancing interpretations of the degree of facial elongation in larger species of some taxa. Felidae: small sand cat (*Felis margarita*) vs large lion (*Panthera tigris*), Canidae: small Fennec fox (*Vulpes zerda*) vs large wolf (*Canis lupus*), Bovidae: small Salt’s dik-dik (*Madoqua saltiana*) vs large Hartebeest (*Alcelaphus buselaphus*). Images not to scale.

### (3) Are we seeking evidence of one pattern or several?

While facial elongation might arise from certain underlying allometric effects, a potential issue with the testing for the presence of suspected craniofacial allometric patterns is having them conflated with other known allometric patterns acting on cranial shape. An obvious candidate is Haller’s rule (Rensch, 1948), which states that larger species of a given clade will tend to have relatively smaller brains. Conflation then is likely very common, because if brain volume scales with hypoallometry to body mass (i.e., with a slope of <1; Martin, 1981; Weisbecker & Goswami, 2010; Smaers et al., 2021) and cranial size is often used as a proxy for body mass because of the consistently high correlations between cranial size and body size (Hood, 2000; Ercoli and Prevosti, 2011; Meloro and O’Higgins, 2011; Cassini et al., 2012), it follows that the size of the braincase can also be expected to scale to cranial size with a slope of <1. Accordingly, relatively smaller braincases among species with larger cranial sizes are often observed in studies of skull shape (Frost et al., 2003; Klingenberg & Marugán-Lobón, 2013; Meloro & Slater, 2012; Mitchell et al., 2018; Marcy et al., 2020). Therefore, some morphometric approaches measuring the viscerocranium and neurocranium might not identify elongation of the face as intended, but instead a relatively smaller braincase. Since the neurocranium is almost always relatively smaller in larger species, linear regressions of any single dimensions of neurocranium size with viscerocranium size will be confounded by Haller’s rule and will generally result in a slope of >1 for “face length” over brain size (hyperallometry), despite potentially being only loosely related to allometric patterns that affect the face.

## III REVISITING FACIAL SCALING WITH STANDARDISED METHODS

In the following section, we apply a standardised framework to test for allometric patterns that addresses the aforementioned concerns. Firstly, we limit taxonomic level of investigation to a single level, the family, which has been used in previous studies (Fig. 2). The family level tends to contain sufficient species diversity for a reasonable sample across diverse mammalian taxa from published research. Focusing on the family level is arbitrary and might not be biologically meaningful in terms of morphological and ecological ranges. However, choosing an arbitrary taxonomic level removes the potential for biased sampling due to prior assessment of what kind of lineage might be suitable for a study. Secondly, we provide a standardised definition for what constitutes the face. This accommodates the concerns we outline above: the CREA pattern reported by others does not always involve elongation of the rostrum, but instead a general increase of gracility across the entire facial skeleton, or viscerocranium, with increased body mass across species, henceforth referred to here as “hyperallometric gracilisation”. While elongation of the rostrum might be common, it is not a necessary feature to identify relative gracility, which can also be observed through a narrower or shallower viscerocranium and perhaps its individual features as well (e.g., zygomatic arches, maxillae, etc). This represents a more literal definition of gracility as meaning slender or thin, rather than the absence of broader facial features, as it is often used in anthropology (e.g., “robust” vs “gracile” australopithecines). Under this new condition, the term “face”, and considerations of the influence of relative orbit size, are no longer required. Thirdly, we more explicitly observe hyperallometric gracilisation in association with hypoallometry of the braincase through allometric predictions of shape differences. These predictions can be obtained through analysis of landmarks representing gross cranial morphology. All specimens in a given study need to have homologous landmarks. But because our hypothesis relates to overall cranial proportions, the homology is not necessary between data from different studies as long as the same features are being described (e.g., landmarks from the premaxillae representing the anterior rostrum), and as long as landmarks encompass major dimensions of the entire cranium. This rationale permits the comparison of outputs from diverse landmarking protocols (see Appendix B). We use examples of both 3D (length/width/depth) and 2D (length/width) landmark data to show that hypotheses relating to hyperallometric gracilisation can be evidenced in both formats. By using landmarks representing the entire braincase and facial skeleton, shape differences of the viscerocranium proportions along an allometric gradient can be visually identified alongside changes in relative brain size expected from Haller’s rule, yet not confounded by Haller’s rule.

We apply this approach to 22 taxonomic families of mammals, and test for patterns of hyperallometric gracilisation across the ranges of cranial sizes. The data used are geometric morphometric landmark coordinates representing whole crania from undomesticated mammals, sourced from published studies. These data are analysed with a standard suite of multivariate statistics frequently used in shape analysis, involving a combination of ordinary least squares regression (OLS), principal component analysis (PCA), and phylogenetic generalised least squares regression (PGLS). Given that elongation is often represented in PCA in the first few dominant principal components (PCs) for most geometric morphometrics-based shape analyses (reviewed e.g., in Weisbecker et al., 2021), we explore relationships between size and the first three PCs individually. We then generate shape predictions along the size range of each family. Importantly, predictions for the sample are generated through estimates of shape scores along the regression line of shape∼log(cranial size), and a regression line for the sample will exist whether the regression is significant or not. Therefore, evidence of CREA requires any instances of more gracile cranial proportions predicted in larger species to be accompanied by statistical support for allometry from the model of shape∼log(cranial size), which we perform both with and without phylogenetic considerations.

We also test for correlations of phylogeny and size within each family by estimating phylogenetic signal of cranial size, as this could influence the results of phylogenetic comparative methods (see Methods in Appendix B. R scripts with data for each family are available at https://github.com/DRexMitchell/Mitchell-etal-facial-scaling). As allometric tests would not be meaningful across a group with very similar sizes, we also generate an approximation for size diversity within each family with a “centroid size index” (CSI), by dividing the largest cranial size in each sample with the smallest cranial size. We then test how strongly the CSI correlates with the coefficient of determination (R^2^) values of allometry tests to see whether size disparity influences the amount of variation attributable to allometry. With the above approaches, we show that the allometric pattern of facial gracilisation is common, but not necessarily ubiquitous, among mammals. We then demonstrate how the occurrence of size-related functional traits can be obscured when using phylogenetic comparative methods, and that abrupt shifts in ecology across size gradients can subvert existing patterns. For the remainder of the review, we present a theoretical framework detailing why allometric facial gracility is most often a by-product of shifting selective pressures incurred with increasing body size on a macroevolutionary scale, and to suggest the place of facial scaling patterns in our understanding of skeletal allometry as a whole.

## IV GRACILISATION PATTERNS IN MAMMALIAN FAMILIES

The results of our analyses across 22 mammalian families are summarised in Table 1, and a more detailed interpretation for each family is provided in Appendix C. We accept a P-value of 0.052 for the Herpestidae as marginal evidence of allometry, given its support in prior research (Cardini & Polly, 2013). Evidence of evolutionary allometry was found for 20 of the 22 families, as given by statistically significant OLS regressions (without including phylogenetic relatedness). The two exceptions were families of bats, the Phyllostomidae and the Molossidae. The CSI was not correlated with the R^2^ values of allometric tests across the families (R^2^=0.028, p=0.501), indicating that the strength of allometry is not influenced by the scale of size variation. Therefore, the two exceptions were not due to a smaller range of cranial sizes in these families, because there were several families with smaller skull size ranges (CSI in Table 1) that did exhibit allometry.

**Table 1:**
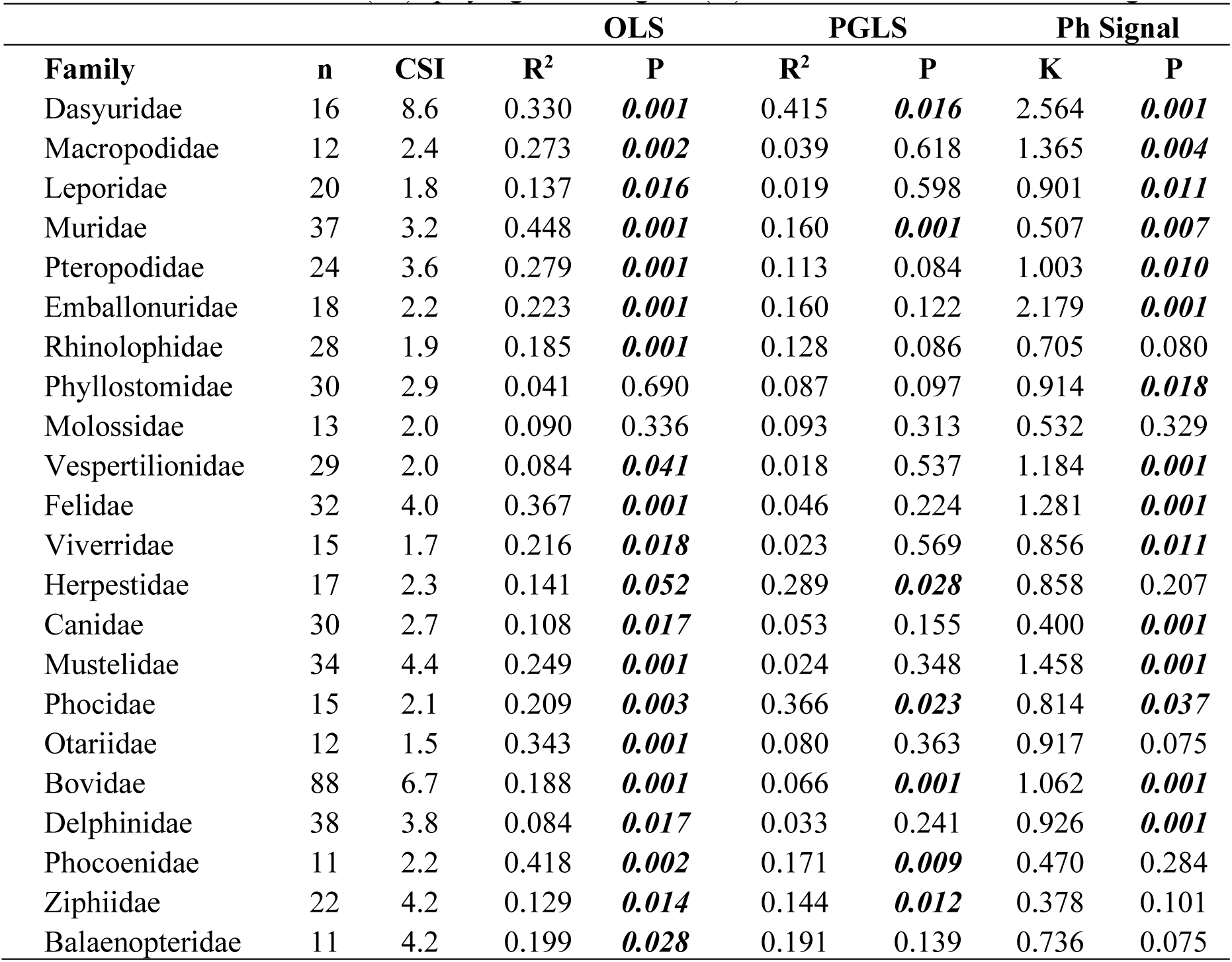
Results for tests of allometry (OLS and PGLS) and phylogenetic signal of cranial size across all 22 mammalian families. All except the Dasyuridae are from published data (see Appendix B). Number of species (n), cranial size index (CSI = largest size/smallest size), coefficient of determination (R^2^), phylogenetic signal (K) and associated P-values are given.

| Family | n | CSI | OLS |  | PGLS |  | Ph Signal |  |
| --- | --- | --- | --- | --- | --- | --- | --- | --- |
| | | | $R^2$ | P | $R^2$ | P | K | P |
| Dasyuridae | 16 | 8.6 | 0.330 | **0.001** | 0.415 | **0.016** | 2.564 | **0.001** |
| Macropodidae | 12 | 2.4 | 0.273 | **0.002** | 0.039 | 0.618 | 1.365 | **0.004** |
| Leporidae | 20 | 1.8 | 0.137 | **0.016** | 0.019 | 0.598 | 0.901 | **0.011** |
| Muridae | 37 | 3.2 | 0.448 | **0.001** | 0.160 | **0.001** | 0.507 | **0.007** |
| Pteropodidae | 24 | 3.6 | 0.279 | **0.001** | 0.113 | 0.084 | 1.003 | **0.010** |
| Emballonuridae | 18 | 2.2 | 0.223 | **0.001** | 0.160 | 0.122 | 2.179 | **0.001** |
| Rhinolophidae | 28 | 1.9 | 0.185 | **0.001** | 0.128 | 0.086 | 0.705 | 0.080 |
| Phyllostomidae | 30 | 2.9 | 0.041 | 0.690 | 0.087 | 0.097 | 0.914 | **0.018** |
| Molossidae | 13 | 2.0 | 0.090 | 0.336 | 0.093 | 0.313 | 0.532 | 0.329 |
| Vespertilionidae | 29 | 2.0 | 0.084 | **0.041** | 0.018 | 0.537 | 1.184 | **0.001** |
| Felidae | 32 | 4.0 | 0.367 | **0.001** | 0.046 | 0.224 | 1.281 | **0.001** |
| Viverridae | 15 | 1.7 | 0.216 | **0.018** | 0.023 | 0.569 | 0.856 | **0.011** |
| Herpestidae | 17 | 2.3 | 0.141 | **0.052** | 0.289 | **0.028** | 0.858 | 0.207 |
| Canidae | 30 | 2.7 | 0.108 | **0.017** | 0.053 | 0.155 | 0.400 | **0.001** |
| Mustelidae | 34 | 4.4 | 0.249 | **0.001** | 0.024 | 0.348 | 1.458 | **0.001** |
| Phocidae | 15 | 2.1 | 0.209 | **0.003** | 0.366 | **0.023** | 0.814 | **0.037** |
| Otariidae | 12 | 1.5 | 0.343 | **0.001** | 0.080 | 0.363 | 0.917 | 0.075 |
| Bovidae | 88 | 6.7 | 0.188 | **0.001** | 0.066 | **0.001** | 1.062 | **0.001** |
| Delphinidae | 38 | 3.8 | 0.084 | **0.017** | 0.033 | 0.241 | 0.926 | **0.001** |
| Phocoenidae | 11 | 2.2 | 0.418 | **0.002** | 0.171 | **0.009** | 0.470 | 0.284 |
| Ziphiidae | 22 | 4.2 | 0.129 | **0.014** | 0.144 | **0.012** | 0.378 | 0.101 |
| Balaenopteridae | 11 | 4.2 | 0.199 | **0.028** | 0.191 | 0.139 | 0.736 | 0.075 |

Seven families (∼32%) had significant evolutionary allometry when phylogeny was included in the model (PGLS; Table 1). Interestingly, evidence for evolutionary allometry in the PGLS and a phylogenetic signal of cranial size might often be mutually exclusive: 14 of the 22 families tested (∼64%) had either a significant phylogenetic signal of centroid size (11/14) or significant evolutionary allometry (3/14), but not both. This suggests that size-related shape variation can be obscured by phylogenetic corrections in statistical analysis when size has a significant phylogenetic signal, which adds an additional complication to assessments of cranial allometry that deserves further research. The finding is likely attributable to the prevalence of size-correlations in phylogenies, such as Cope’s rule and others, across mammals. Covariation of size and phylogeny make it impossible to statistically distinguish between shape variation attributable to either evolutionary divergence unrelated to size or shape variation causally related to size, such that factoring the phylogeny into the model would result in a lack of significance of potentially existing evolutionary allometry. In cases where a phylogenetic signal is present in the predictor trait (cranial size), inclusion of phylogenetic relatedness in tests should not be necessary (Rohlf, 2001; 2006; Uyeda et al., 2018).

A summary of the presence of hyperallometric gracilisation across the size ranges of each family, assessed through visual inspections of predicted landmark vector displacements, is presented in Figure 4. All families exhibiting evolutionary allometry display relatively smaller braincases in larger species, in agreement with Haller’s rule. However, predictions for hyperallometric gracilisation, requiring both significant evolutionary allometry and a more gracile viscerocranium predicted for larger sizes, were more variable (see all significant predictions in Appendix C). Eleven families showed evidence of allometric facial gracility, while nine showed a reverse trend of more stout features (any or all of shortening/widening/deepening of the viscerocranium) predicted in larger species. Among analyses of the seven families with significant evolutionary allometry after phylogenetically informed tests, five exhibited increased gracility in species with larger crania. Therefore, a finding of significant allometry in a PGLS test alongside a significant phylogenetic signal of cranial size might indicate that size variation is influenced by a developmental constraint on shape that is consistent regardless of relatedness. This would agree with previous suggestions that allometry explains the majority of cranial shape diversity in some of these taxa, such as the Bovidae (Bibi & Tyler, 2022; Rhoda et al., 2022) and rodents (Marcy et al., 2020). However, an alternative that could be investigated in the future is that phylogenies comprising speciose taxa with multiple monophyletic groups each spanning much of the total size ranges for the clade, such as bovids and rodents, can limit the impact of including the phylogeny in the model and result in similar statistics to the OLS. In contrast, phylogenies with small-sized monophyletic clades and large-sized monophyletic clades might have more of an impact on congruence of the two statistical approaches.

**Figure 4:**
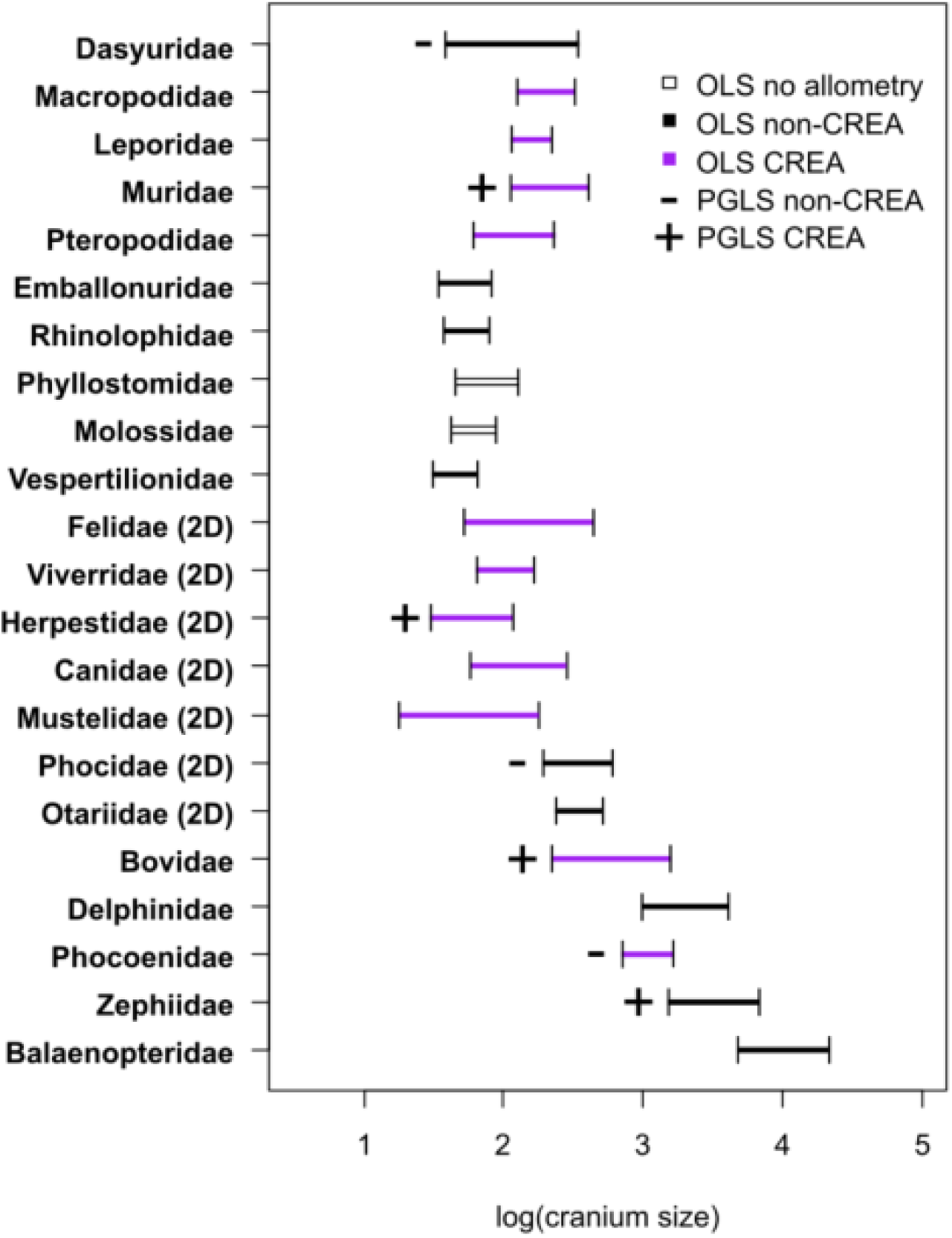
Hyperallometric gracilisation presence or absence across the 22 families of mammals. Support for the pattern requires both a significant result (α ≤ 0.052) for evolutionary allometry and a more gracile viscerocranium predicted for larger sizes (purple for OLS and + for PGLS, as given in legend). For the 2D basicranial landmarks of the Carnivora, cranium sizes were adjusted to be comparable to sizes with a third dimension (see Appendix B).

Importantly, Figure 4 demonstrates that both smaller-bodied and larger-bodied families exhibit examples consistent and inconsistent with hyperallometric gracilisation of the cranium, such that large species can have more stout skulls than smaller relatives within a family, just as small species can have longer, more slender skulls than larger species. As a result, facial gracility cannot be reliably predicted from cranial size. This result speaks against a strong influence of a universal developmental constraint as a source of allometric facial scaling in mammals, which would be expected to result in predictable and repeatable allometric changes across certain size thresholds.

While our allometric shape predictions are useful to examine the broad trends of allometric variation, our analyses include an important caution about the assessments of patterns across taxa with high levels of shape variation, which can obscure the diversity inherent in the taxon in question. To illustrate this point, we will here present a more detailed analysis of the 2D-landmark-based data from three Carnivoran clades (Figure 5). We focus on the cats (Felidae; Fig. 5A), weasels and relatives (Mustelidae; Fig 5B), and wolves and relatives (Canidae; Fig. 5C). Figure 5 uses Principal Component Analysis plots to demonstrate that producing a single allometric prediction along the size gradient of an entire clade can miss many important nuances of allometry. For example, Fig. 5B shows an allometric facial elongation pattern on PC1, but also a second allometric trajectory among the Mustelidae that is uncorrelated with facial elongation and represents facial stoutness. This is exemplified by the sea otter (*Enhydra lutris*) at the maximum of PC2, a species with an extremely stout cranium that can reach a body mass greater than 30kg (Laidre et al., 2006) (Fig. 5B). Otters therefore present an example of a taxonomic subset of the Mustelidae that would not follow a pattern of hyperallometric gracilisation, showing that not all taxonomic subsets can be expected to conform to a broader pattern of allometric facial elongation across the family (e.g., as found in the subsets in Cardini et al., 2015 and Tagmanini et al., 2017).

**Figure 5:**
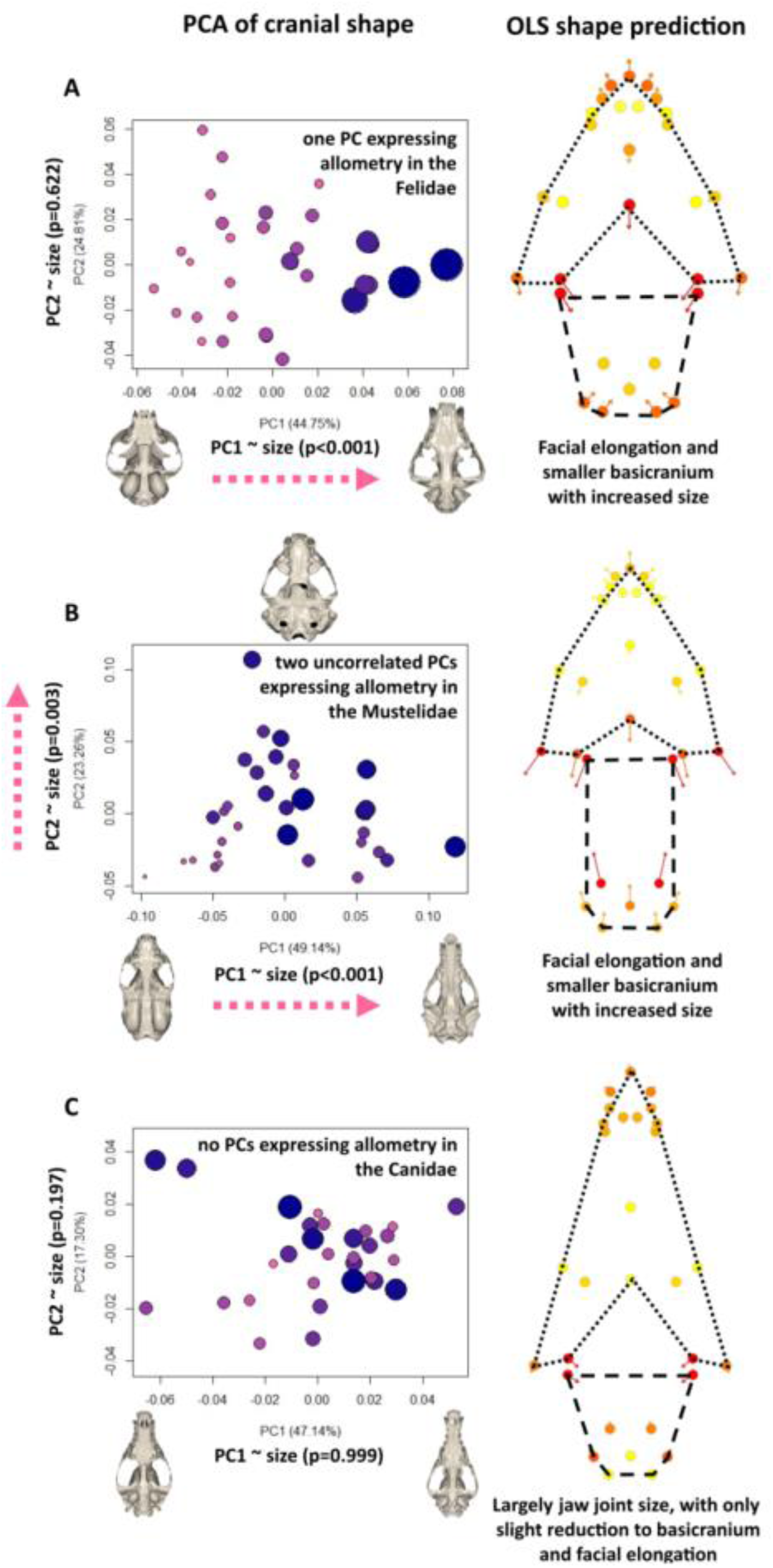
Results for tests for evolutionary allometry using 2D ventral landmarks of three families from the Carnivora. (A) Felidae, (B) Mustelidae, (C) Canidae. Allometry is depicted in the PC scatterplots (left), where size is denoted by both colouration and point size. 3D meshes represent species crania occupying the PC extremes. Pink arrows indicate significant allometric relationships. Allometric shape predictions are depicted in the accompanying shape change graphs (right), where landmark vectors are coloured as heatmaps proportional to magnitude of shape variation, with the points representing the shape of the smallest size and end of arrows representing largest size. Viscerocranium = dotted lines, braincase = dashed lines.

Despite the above exception, we found evidence for hyperallometric gracilisation in more than half of the families tested, suggesting that the pattern is quite common. However, as with the otter sample discussed above, there are easily conceivable evolutionary scenarios where the pattern might represent a substantial component of cranial evolution, without strong (or indeed, any) evidence from visualisations of allometric shape variation. We demonstrate this on the example of the Canidae, by showing how hyperallometric gracilisation can be observed at finer taxonomic scales (see Appendix C for more detailed analyses). The phylogeny of Canidae (Fig. 6A) can be broadly separated into three monophyletic groups: One dominated by foxes (*Vulpes* spp.), one dominated by South American foxes (*Lycalopex* spp.), and one containing wolves, dogs, and coyotes (*Canis* spp.). Hyperallometric gracilisation is present among the *Vulpes* group crania alone (R^2^=0.220, p=0.044) (Fig. 6B). However, this effect disappears outside of the *Vulpes* clade because the bush dog (*Speothos venaticus*), the smallest of the *Canis* clade, is larger than most species of the *Vulpes* clade, but has the most robust cranium in the entire Canidae family. Therefore, while the smaller *Vulpes* clade might appear at a glance to be at odds with hyperallometric gracilisation, in being “longer faced” and smaller than the more “short faced” species, the pattern can be found in the *Vulpes* branch when observed in isolation. Notably, the same can be found in the *Canis* branch, but only when accounting for their feeding strategies.

**Figure 6.**
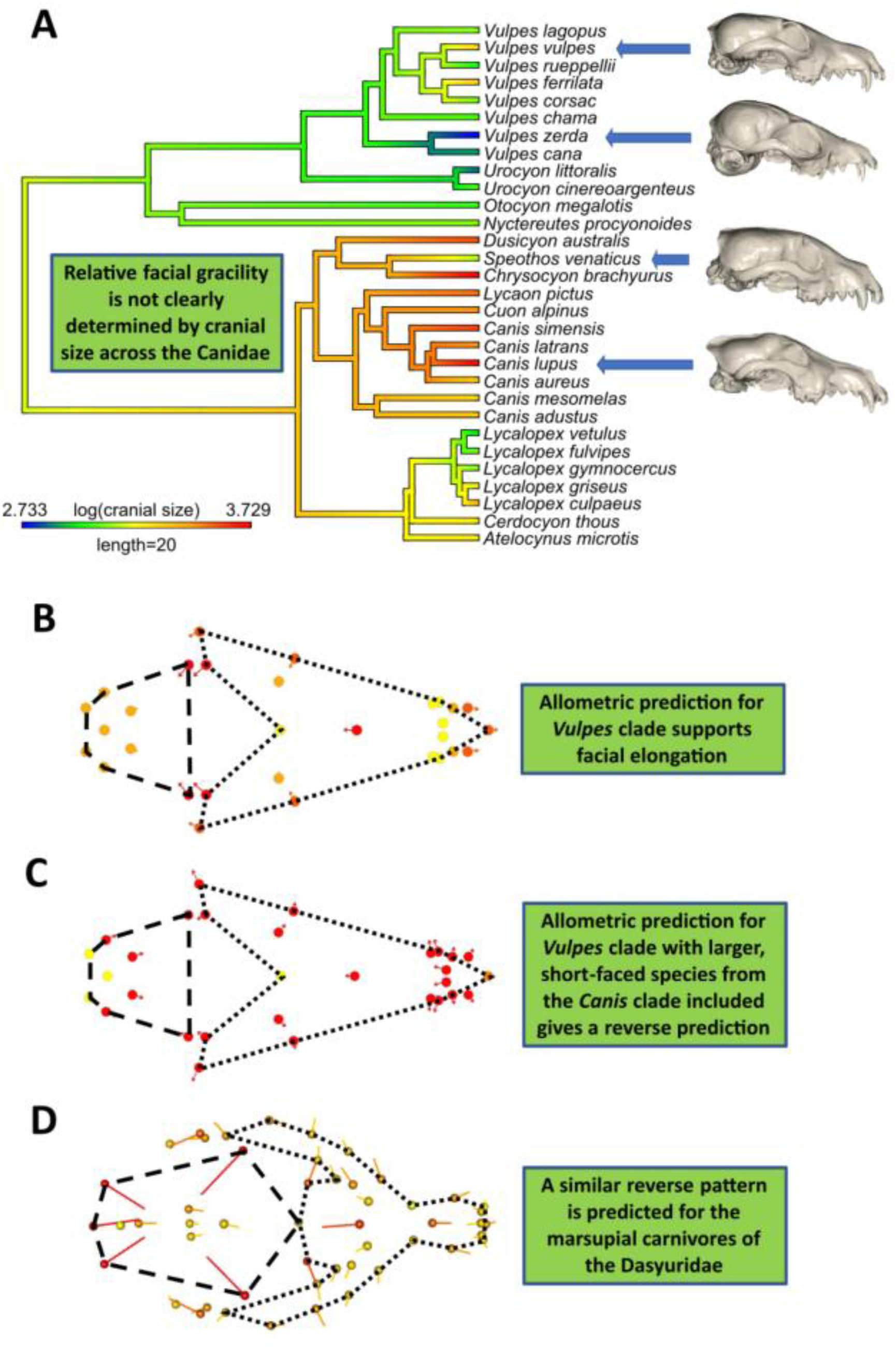
Testing for hyperallometric gracilisation at a finer scale within the crania of the Canidae.

The *Canis* clade is unique within the Canidae, in comprising both small-prey specialists and large-prey specialists, where the remaining two clades only specialise on small prey (see Van Valkenburgh & Koepfli, 1993; Christiansen & Wroe, 2007 for classifications). The largest of the *Canis* clade, the wolf (*Canis lupus*), has a more gracile cranium than the remaining three large-prey specialists (*S. venaticus, Lycaon pictus*, and *Cuon alpinus*) that represent the stout shape extremes of the Canidae in Fig. 5C. But *C. lupus* also has a more stout cranium than the small-prey specialists within the *Canis* clade (see Appendix C). This results in no evidence of allometry within the *Canis* clade alone because the more stout large prey specialists envelope the small prey specialists along their size range. Yet, among the large-prey specialists alone, larger crania are noticeably more gracile. Together, these trends among the *Vulpes* and *Canis* clades and the results and interpretations of the other families tested (Appendix C) suggest that the scale of relatedness for testing craniofacial allometry is largely arbitrary and the presence of hyperallometric gracilisation often depends on the consistency of the dietary ecology for all species involved.

With this in mind, the dataset of Canidae can be used to demonstrate how the introduction (or removal, e.g., through extinction) of ecomorphs within a clade can disrupt overall trends of hyperallometric gracilisation. When a hypothetical clade is generated that includes the *Vulpes* clade of small-prey specialists and also the three most stout morphologies of the large-prey specialists from the *Canis* clade, the linear model shows a significant association of shape with size (R^2^=0.175, P=0.044), but instead predicts larger species to have wider crania (Fig. 6C). This is because the three larger species included represent a more stout ecomorph capable of accommodating the mechanical demands of more resistant foods; in this case, large prey. A similar effect can be observed in our dataset of marsupial carnivores, the Dasyuridae. This is due to the broad-faced, bone-cracking morph of the largest species, the Tasmanian devil (*Sarcophilus harrisii*), positioned as a sister taxon to the quolls *(Dasyurus* spp.) on the Dasyuridae phylogeny. The above examples show that predictions for hyperallometric gracilisation appear highly sensitive to the occurrence of ecological shifts that occur towards size extremes. Our analyses exhibit multiple instances of species occupying divergent dietary niches within their respective clades (see Appendix C), generally associated with ecomorphological shifts of their respective cranial shapes that can result in less gracility at larger sizes, or increased gracility at smaller sizes. In summary, if niche-specific adaptations resulting in stouter proportioned crania occur in larger species within a clade; this appears to impact predictions of facial gracility with increased size (Cardini & Polly, 2013; Tamagnini et al., 2017). This concept of common and divergent ecomorphs is, in principle, similar to the common ecomorphological “bauplan” discussed in previous publications on CREA (Cardini et al., 2015; Tamagnini et al., 2017; Cardini, 2019), and is a highly relevant point when considering the inclusion of extinct species, many of which possibly occupied niches no longer utilised by their extant relatives.

The above shows that the common approach of studying only in extant taxa can be an issue if a clade contains extinct ecomorphs that are not represented in the extant sample. It is well-known that inclusion of only extant morphologies has the potential to oversimplify, misidentify, or misrepresent morphological diversity and macroevolutionary patterns (Hautier et al., 2014; Raj Pant et al., 2014; Cuff et al., 2015; Jablonski, 2019). This issue is relevant to the majority of studies specifically examining craniofacial allometry, and CREA in particular, which have only included extant species, with the exceptions of Gomes Rodrigues et al. (2018), Krone et al., (2019), and the cetacean dataset we used here from Coombs et al. (2020). The lack of fossil representation is understandable because many fossil specimens can exhibit damage and deformation that can limit their use in shape analysis (Mitchell et al., 2021), but this nonetheless presents a clear issue with testing an accurate approximation for cranial diversity within clades (see discussion of this issue by Tamagnini et al., 2017). Many mammalian taxa include extinct stout skull morphs that could influence the results for tests of facial elongation. These include bone-cracking borophagine canids (Van Valkenburgh et al., 2003; Tseng & Wang, 2010) and hyaenids (Turner & Antón, 1996; Palmqvist et al., 2011), durophagous mustelids (Valenciano et al., 2016; Tseng et al., 2017), short-faced bears (Figueirido et al., 2009; Figuierido & Soibelzon, 2010), giant short-faced kangaroos (Prideaux, 2004; Mitchell, 2019; Mitchell & Wroe, 2019), glyptodont armadillos (Machado et al., 2022), merycoidodont artiodactyls (Greaves, 1972), and pantodontids (de Muizon et al., 2015). Such examples showcase an apparent abundance of stout cranial morphologies that have not been included in tests thus far and we revisit the abundance of stout cranial morphs in the fossil record in Section VIII. But for now, we focus on the fact that, despite many exceptions, increases of facial gracility with size are a common occurrence across mammals, particularly among those with a common ecomorphology. This leads us to ask: what might be driving the frequent pattern of hyperallometric gracilisation in extant species groups? In order to answer this question, we begin with the question of what conditions can govern the evolution of more stout cranial proportions.

## V WHY THE SHORT FACE? CRANIAL ADAPTATIONS FOR HARDER BITING

To assess the mechanisms behind deviations from allometric cranial gracility, it is imperative to contextualise the mammalian cranium with its crucial role in food prehension and processing. Biting biomechanics present a strong functional constraint on cranial shape, and much of the morphological variation of the mammalian feeding apparatus at a macroevolutionary scale is considered a product of adaptations that accommodate the most strenuous biting activities employed (Van Valkenburgh, 1989; Strait et al., 2009; Santana et al., 2012; Figueirido et al., 2013; Mitchell, 2019), further enhanced by safety factors to withstand occasional extremes (Alexander, 1981; Willie et al., 2020). In the assessment of functional morphology of the cranium, feeding groups are therefore often better predicted by the most challenging foods consumed (Santana et al., 2010; Zelditch et al., 2017; Hedrick & Dumont, 2018; Mitchell & Wroe, 2019). This is because cranial features of biomechanical significance can arise in parallel across different clades for similar functional demands, in a manner that is independent of body size and instead more associated with how hard an animal can bite (Radinsky, 1985; Emerson & Bramble, 1993; Figueirido et al., 2012).

The manner by which a given species interacts with its environment can, in part, be delineated by size-independent (absolute) bite force capability. Bite force can determine the ability for an animal to manipulate, obtain, or process foods and other resources. Vertical force during biting is a product of muscle size (muscle cross sectional area) and leverage (Popowics & Herring, 2006), but a skull must also be able to accommodate the bone stresses associated with the hardest bites. Figure 7 offers first-principles schematic representations of well-established and tested cranial adaptations involved in the production and accommodation of harder bites. In summary, adaptations for harder biting can involve: an increase of bite force via increased muscle mass (Fig. 7A-C; Greaves, 1985; Wroe et al., 2005; Herrel et al., 2008; Attard et al., 2011; Tseng, 2013; Tseng & Flynn, 2018), or increased leverage (i.e., mechanical efficiency) through changes to the relative lengths of the in-lever and out-lever (Fig. 7D-F; Herring & Herring, 1974; Van Valkenberg & Koepfli, 1993; Antón, 1996; Biknevicius & Van Valkenburgh, 1996; Ravosa, 1996; Aguirre et al., 2003; Wroe et al., 2005; Christiansen & Wroe, 2007; Wroe & Milne, 2007; Koyabu & Endo, 2009; Nogueira et al., 2009; Figueirido et al., 2010b; Goswami et al., 2010; Dumont et al., 2012; 2014; 2016; Smith et al., 2015; Mitchell et al., 2018; Ledevin & Koyabu, 2019; Mitchell & Wroe, 2019; Giacomini et al., 2021; Harano & Asahara, 2022). In addition, to accommodate the forces experienced in the cranium during hard biting (Greaves, 1991; Covey & Greaves, 1994; Ross & Hylander, 1996; Ross, 2001; Herring et al., 2001; Rafferty et al., 2003; Herring, 2007; Santana et al., 2012; Mitchell, 2019), deepening and buttressing of the skull at specific regions of high stress adds further bone volume as reinforcement (Fig. 7G-H; Constantino, 2007; Strait et al., 2008; Samuels & Van Valkenburgh, 2009; Tseng, 2009; Kitchener et al., 2010; Tseng & Wang, 2010; Wilson & Sanchez-Villagra, 2010; Figueirido et al., 2011; 2013; Wilson, 2013; Dumont et al., 2016; Ledogar et al., 2017; Mitchell, 2019). This reinforcement often happens because bone stress is equal to force per unit of area; hence, an increase in the amount of bone at a specific region of the cranium will result in a decrease in stress for a given magnitude of force at that region (Mitchell, 2019; Mitchell et al., 2021).

**Figure 7:**
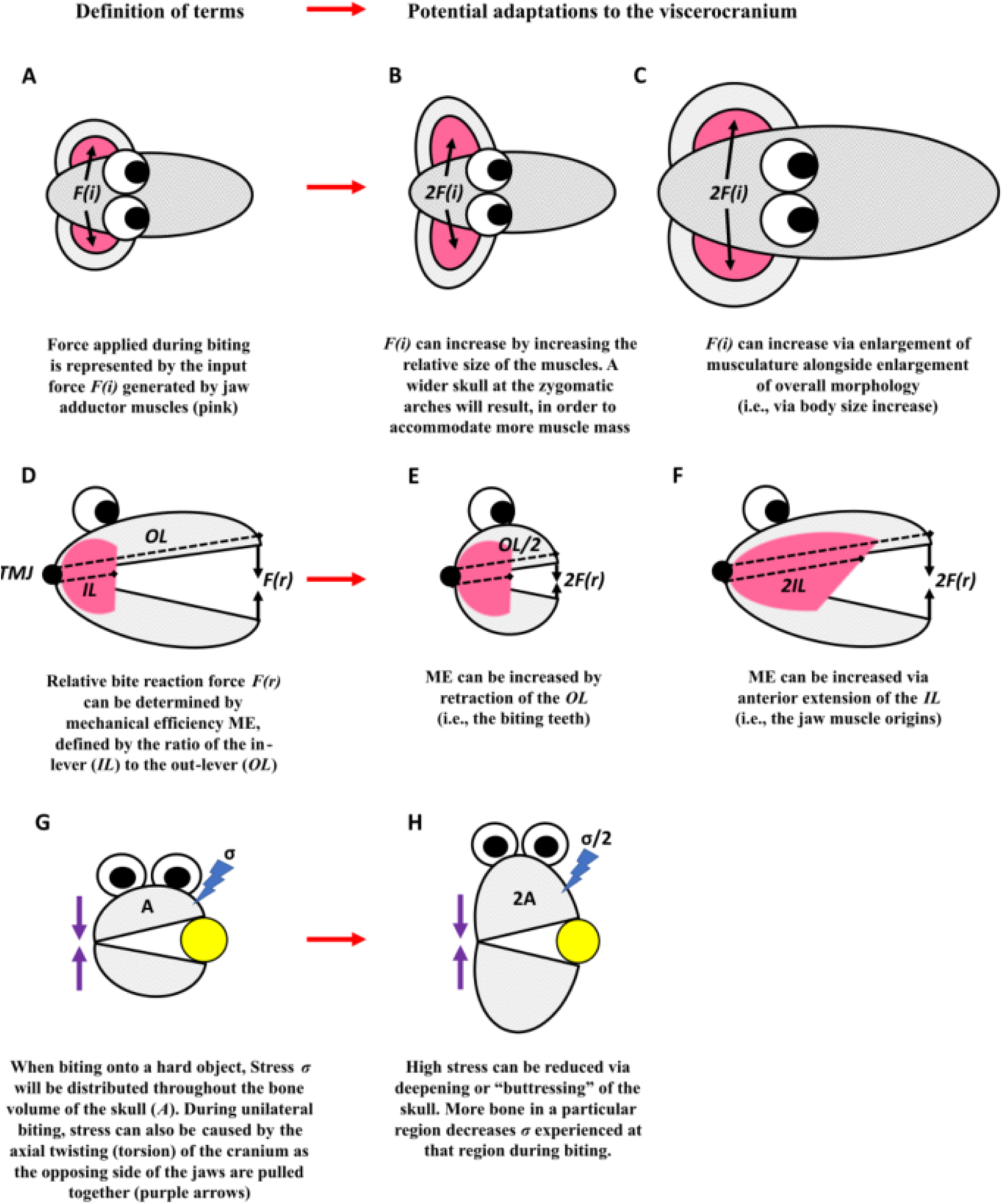
Adaptations for hard biting in the cranial morphology of mammals. Terms are defined (A,D,G), and five scenarios are shown (B,C,E,F,H).

The adaptations given in Figure 7 are not necessarily exclusively associated with hard biting. For example, a deeper maxilla and mandible might also accommodate deeper rooted, continuously growing (hypsodont) molars (Bargo, 2001; Raia et al., 2010). Furthermore, there are also possible adaptations to muscle physiology that might, at least to some extent, mitigate losses to bite force in a more gracile cranium (Wall et al., 2008; Allen et al., 2010; Hartstone-Rose et al., 2012; Martin et al., 2020). However, all the mentioned adaptations nonetheless hold biomechanical significance in improving the skull’s ability to produce and withstand higher bite forces, and crania adapted for harder biting will likely exhibit more than one of them when compared to relatives that do not perform strong bites. In many cases, several of these adaptations will also covary to some extent. With this understanding, facial stoutness would be considered an example of “many-to-one mapping”, whereby multiple evolutionary adaptations result in similar function (Wainwright, 2007; Figueirido et al., 2011; Tseng & Flynn, 2015). Alternatively, the reverse of any of these adaptations (i.e., from more stout to more gracile morphologies) will represent a reduction in either muscle force, mechanical efficiency, or the ability to withstand higher stress during biting. Of course, a species without robust features will not necessarily be incapable of attempting tasks requiring harder bites. For example, a longer-faced animal can add additional bite force for hard foods by increasing the usage of posterior (postcanine) dentition during biting (see Dumont, 1999; Dumont & O’neal, 2004). Yet, such behaviours emphasise the ineffectiveness of the anterior dentition for hard biting in long-faced species, which are the teeth often used by most mammals in the initial prehension of food or other behaviours for environmental manipulation (but see section VIII regarding exceptions). It is also true that an elongate rostrum can increase jaw closure speed (see Section VII), thereby potentially maintaining higher absolute bite forces. However, a longer rostrum is also less efficient at withstanding the stresses of a given bite force than a more compact rostrum (e.g., Mitchell et al., 2018; Mitchell & Wroe, 2019). Therefore, selection for harder biting behaviours over evolutionary time should be accompanied by more stout cranial proportions in many cases.

It is crucial to recognise that all but one of the adaptations to higher bite forces listed in Figure 7 can result in a facial skeleton that could be perceived as more stout, robust, or “shorter”. The important exception is an increase in total body size - the only process also associated with allometry (Fig. 7C). If larger-bodied species within clades frequently exhibit an allometric pattern of cranial gracility, Figure 7 shows that these species must be sacrificing a capacity to generate or withstand higher relative bite forces. To understand the processes behind hyperallometric gracilisation, we must give additional consideration to the distributions of diet regimes, mechanical properties of food, and biting behaviours along allometric gradients, which we discuss in the following section.

## VI ALLOMETRIC DISTRIBUTIONS OF DIET

Size-related shifts in feeding ecology and behaviour can be observed across many mammalian clades (Fleagle, 1985; Bodmer, 1990; Carbone et al., 1999; Chemisquy et al., 2021; Bubadué, et al., 2022), and are thus expected to also play a role in facial allometry. For example, small herbivores are more likely to incorporate nutrient-rich foods including fruits, nuts, and seeds, while larger herbivores tend to include larger proportions of more abundant graze (grasses) and/or browse (leaves, twigs, and stems of trees and shrubs) (Bodmer, 1990; Arman & Prideaux, 2015). Different resources can present variable demands for their access and processing because of their varying mechanical properties, including hardness, toughness, or elasticity (Berthaume, 2016). A shift in resource usage across size ranges within a clade might therefore sometimes result in selection for either increased or decreased absolute bite force. This will, in turn, be evidenced in their cranial morphology and influence predictions of shape along a size gradient. We can summarise this by identifying three potential trajectories of bite force shifts that can occur within a clade.

One scenario is a dietary shift with increasing size that reduces the amount of absolute bite force required. An example can be seen in the polar bear (*Ursus maritimus*). This species recently diverged from omnivorous brown bears during the mid-Pleistocene (Kurtén, 1964). Despite being the largest of extant bears (Loch et al., 2022), polar bears are considered to predominantly focus on prey much smaller than themselves, such as juvenile seals. There also tends to be a strong preference for the consumption of blubber (Perry, 1966). While increasing the size differential between predator and prey allows this species to overpower prey with body size rather than maximising bite force (Figueirido et al., 2011), blubber is also likely to be less biomechanically demanding. The smaller carnassial blades (Sacco & Van Valkenburgh, 2004), reduced jaw muscle leverage (Sacco & Van Valkenburgh, 2004; Figueirido et al., 2009), and biomechanically weaker skull (Slater et al., 2010; Oldfield et al., 2012; Goswami et al., 2013) of *U. maritimus* compared to close relatives has been suggested to have resulted from this shift in feeding ecology. Larger-bodied species that have switched through the course of their evolutionary history to specialise on foods that, uncharacteristically for their close phylogenetic associations, involve little to no biting, such as insect-feeding (entomophagy), would also be aligned with this scenario. If the maximum absolute bite force requirement decreases with increased size, facial morphology can become more gracile or structurally weaker over evolutionary time.

A second scenario is an ecological shift with increasing size that raises the amount of absolute bite force used. This is expected when larger species incorporate more mechanically demanding resources into their diets than smaller closely related species or participate in novel behaviours requiring harder bites (e.g., incisal digging; Samuels & Van Valkenburgh, 2009; Gomes Rodrigues & Damette, 2023). Bone cracking, shell crushing, consumption of seeds and unripe fruit (sclerocarpy), and heavier browsing are examples of such dietary shifts, and might also include “fallback foods”, consisting of generally less desirable resources that tend to be only exploited during less productive times (Constantino & Wright, 2009; Strait et al., 2010). If the bite force requirement relative to cranial size increases along an allometric gradient, the facial skeleton should exhibit adaptations capable of delivering and withstanding harder biting, as described in Figure 7. This would lead to interpretation of a shortening of the face, contrary to trends of gracility observed elsewhere. Note that a trend appearing as reverse to hyperallometric gracilisation would also be observed if smaller species within a clade have secondarily specialised on less mechanically challenging diets, such as nectar feeding (see Appendix C for examples).

A third scenario occurs when the absolute bite force requirements remain mostly consistent across a size gradient of species. This occurs when little evolutionary change occurs in either mechanical properties of food ranges or absolute bite forces performed for niche-specific behaviours. An obvious example is the mostly uniform material properties of grasses (Jarman, 1974; Shipley, 1999) that change very little across the entire size gradient of grazers. If the maximum bite force requirements remain mostly consistent across the allometric gradient, the viscerocranium can become more gracile because a consistent absolute (size-independent) bite force requires a lower relative (size-dependent) bite force in larger species, and therefore mechanical efficiency can be sacrificed while maintaining consistent bite force capacity (Tseng, 2013).

### (1) A mechanism: bite force allometry and niche conservatism

We propose a prevalence of the third scenario outlined above - retention of dietary mechanical resistance across body size ranges - as the underlying reason for the common occurrence of hyperallometric gracilisation in the mammalian cranium. It offers a particularly powerful explanation when considering a widespread pattern known as phylogenetic niche conservatism, involving the retention of ecological traits over time among related species (Wiens & Graham, 2005; Wiens et al., 2010; Losos, 2008). In other words, more closely related species tend to be more ecologically similar. In the context of hyperallometric gracilisation, we suggest that phylogenetic niche conservatism can be observed in mammals when closely related species share more similar diets and/or absolute bite force requirements. Phylogenetic niche conservatism relating to diet has been widely quantified across animals (Olalla-Tárraga et al., 2017; Fraser et al., 2018; Roman-Palacios et al., 2019). Thus, mapping diet onto any mammalian phylogenetic tree will show a tendency for more closely related species to preference similar foods, regardless of their body sizes (e.g., Cruz-Neto et al., 2001; Arman & Prideaux, 2015; Mitchell et al., 2018; Kartzinel & Pringle, 2020; Melstrom et al., 2021). Within these groups of species with similar diets, a skull capable of high mechanical efficiency and low muscle force can generate comparable absolute bite force to a skull with low mechanical efficiency and high muscle force (Tseng, 2013). Assuming that bite force requirements remain relatively consistent for larger species with similar niche specifications, the relative reduction of mechanical constraints offered by increased size can therefore permit cranial gracility, without sacrificing mechanical safety factors the bone.

Some taxa might appear at first glance not to follow predictions whereby hyperallometric gracilisation of the cranium is a product of consistent bite forces across body size ranges. There are in fact many cases where larger species of some mammalian groups exhibit a broadening of the diet to include items with greater mechanical resistances (Lucas & Luke, 1984), which yet also have more elongate skulls. For example, the largest species of the Cervidae, the moose (*Alces alces*), feeds on browse from the same trees and shrubs as smaller, sympatric cervid species. However, this species also consumes thicker twigs or branches of these plants (Shipley et al., 1999; Nichols et al., 2015). This naturally involves greater bite forces, however, the facial skeleton of the moose has more gracile proportions than smaller relatives (Schilling et al., 2019; Rhoda et al., 2023), contrary to our above arguments. Similarly, the Felidae clearly exhibit a pattern of hyperallometric gracilisation (Fig. 5A), but the tiger (e.g., *Panthera tigris*) does not have absolute bite forces comparable to the smallest cat species, as they subdue and process larger prey (Christiansen & Wroe, 2007) with diverse tissues, including tough hide, sinew, and bone (Pollock et al., 2021). For both moose and tigers, these dietary additions occur alongside the increase of bite forces gained through the stronger jaw muscles from simply being larger (Fig. 7C). For such instances, hyperallometric gracilisation will be found as a product of the relative gradients of bite force increasing alongside increased body mass, and food resistance increasing with increased body mass. This is described in the following equations:

1. Bite force (BF) increases with body mass (BM) at a rate of:

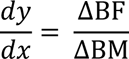
2. And if food resistance (FR) increases with body mass, this occurs at a rate of:

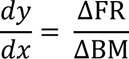
3. Then the relationship between absolute bite force and food resistance across body size ranges is given by:

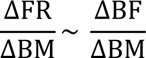

We refer to the slope of this relationship as the allometric bite coefficient (ABC). If ABC = 1, there is isometry in the biomechanical demands of the cranium across size ranges (Fig. 8A). If ABC > 1, the biomechanical demands will be greater and non-size adaptations for harder biting shown in Figure 7 are more likely to be present in larger species. If ABC < 1, a more gracile facial skeleton is more likely. In other words, for a given change in body size, if the rate of increase to food resistance is less than the rate of increase to bite force capacity, this results in reduced biomechanical demand on the facial skeleton, permitting facial gracility in larger species.

**Figure 8:**
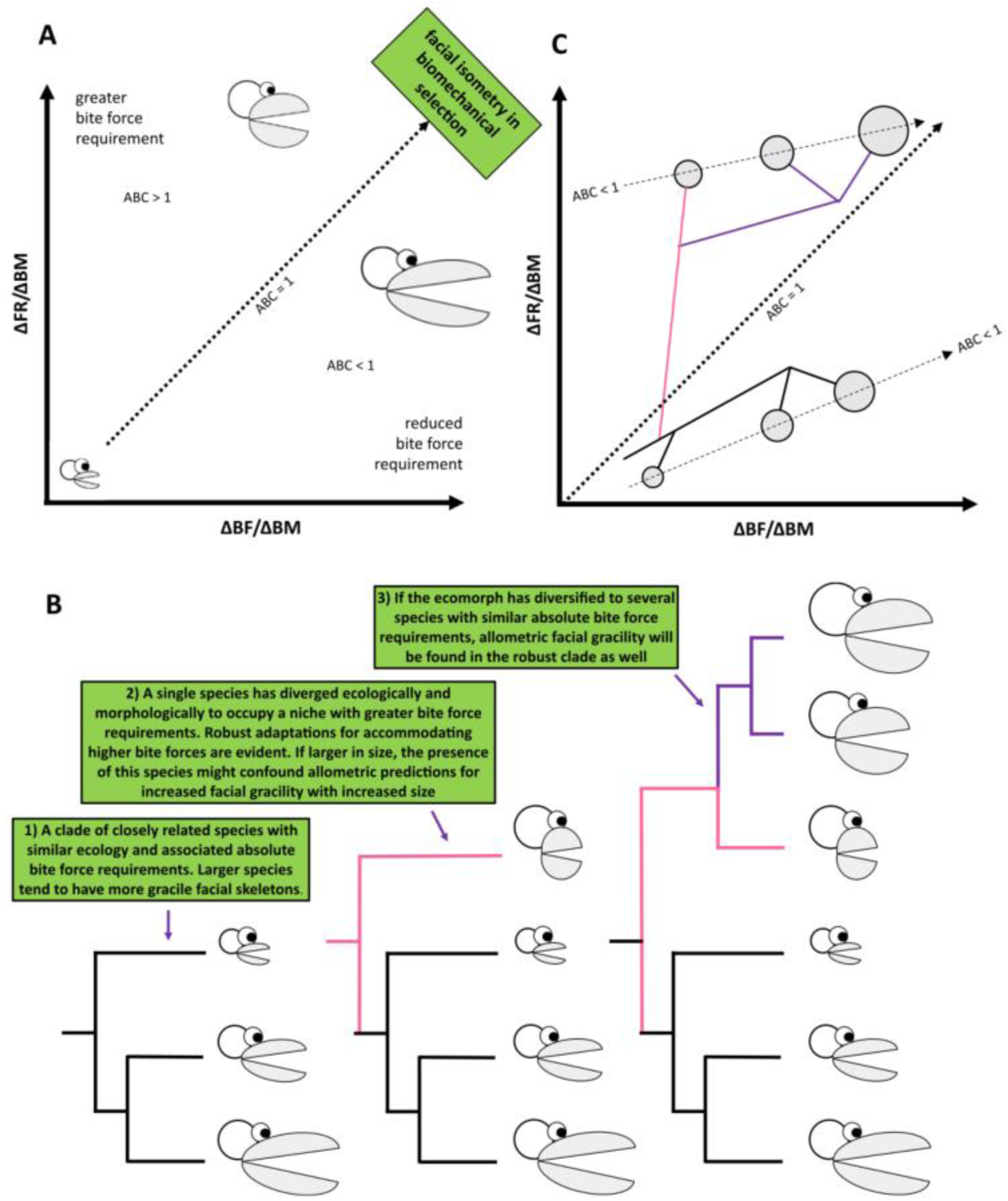
**(A)** The Allometric Bite Coefficient (ABC) is defined by the slope of the correlation between increased bite force with increased body mass (ΔBF/ΔBM) and increased food resistance with increased body mass (ΔFR/ΔBM). For a clade with similar ecology, where larger species incorporate more resistant plant or animal tissues into their diet, allometric facial gracility can be found when the slope (ABC) is less than 1. (B) The presence or absence of allometric facial gracility as a product of phylogenetic niche conservatism and ecological divergence. (C) Mapping the previous phylogeny onto the ABC space shows how facial gracility can be present in diverged ecomorphs.

Importantly, any future considerations of bite force allometry must also quantitatively factor in the hypoallometry of muscle force common across mammals, known as the 2/3rd power rule. In other words, larger species have relatively weaker bites than smaller species because muscle force scales with cross-sectional area, while body mass scales by volume (Alexander, 1985; Wroe & Sansalone, 2023). However, this will not ultimately affect the relationship defined by the ABC, because the 2/3rd power rule is an intrinsic aspect of the rate by which muscle force scales with size, thus standardising its effects across all observations.

Our analyses in Section III demonstrate that hyperallometric gracilisation should be more often observed in the crania of clades where species share similar diets and biting behaviours. However, consistent with the concept of biomechanical adaptations outlined above, the pattern often does not occur in allometric predictions when species deviate from relatives in ecomorphology towards substantially more mechanically demanding biting activities at larger sizes. Figure 8B presents a simplified case of how different allometric predictions might be obtained from novel stout ecomorphs represented in a phylogeny, and then again be supportive of allometric facial gracility if the ecomorph has also diversified to multiple species of various body sizes that can accommodate the new greater threshold of absolute bite force. We then map these examples over the ABC regression (Fig. 8C), showing that when a greater food resistance threshold is crossed by a novel ecomorph, subsequent diversification will likely follow a gracility pattern with an ABC < 1, assuming those species also maintain similar ecology. This distribution is reflected well in the effects of dietary shifts from small-prey specialists to large-prey specialists of the Canidae described in Section IV.

In this section, we have detailed instances where hyperallometric gracilisation is a more likely observation within clades. However, there remains the question of why the facial skeletons of larger mammalian species, within an ecologically similar clade, have shown a tendency to become more gracile in the first place, when relaxed biomechanical demands for bite force should permit morphological shifts along any dimensions. We discuss several potential hypotheses in the following section.

## VII SO WHY THE LONG FACE?

A reduction of biomechanical constraints does not translate to selection for a more gracile face. Instead, it is expected to relax selection for harder bite forces, and permits adaptations related to other selective pressures. We suggest that the prevalence of apparent directional selection for hyperallometric gracilisation across clades of ecologically similar mammals is largely a product of three main factors: (1) moderation of bone resources, (2) a range of other, well-documented selective advantages to an elongate viscerocranium, and (3) a secondary influence of negative braincase and orbit allometry.

### (1) Moderation of bone resources

Bone is both heavy and metabolically expensive to produce (Covey & Greaves, 1994; Dumont, 2010; Mitchell, 2019). This leads to the expectation that cranial development evolves to result in the allocation of minimal amounts of bone given a species’ cranial mechanical performance, including the aforementioned safety factors. Thus, larger skulls that experience lighter mechanical loading are expected to be selected for less bone allocation during development. Referencing Figure 7, gracilisation would be an obvious outcome of this mechanism. For example, slighter masticatory muscles with less cross-sectional area can be accompanied by a narrowing of the zygomatic arches (i.e., the reverse of Fig. 7B), and a reduction in bone reinforcement might also produce a more shallow, or less reinforced cranium (i.e., the reverse of Fig. 7H). Both outcomes, which represent a clear saving in bone cost and weight, would create a more gracile facial skeleton. The previously mentioned polar bear with its biomechanically weaker skull architecture compared to close relatives is one example of this pattern.

Within the context of bone resources, the differentiation between ontogenetic, static, and evolutionary scaling processes discussed in section 3 of the Introduction probably represents an issue with the confidence by which cranial gracilisation can be attributed to natural selection. This is because adaptive mechanisms of bone deposition and resorption are well known to occur within the lifetime of an individual animal (Wolff 1892; Frost 1994; Pearson and Lieberman 2004) as an example of developmental plasticity. Variation in bone volume and mineralisation of the skull widely occur in response to shifts in activity levels, including those brought about by diet composition. These processes play a substantial role in intraspecific cranial shape variation and associated mechanical performance, and are often not allometric (Ravosa et al. 2008; Menegaz et al., 2010; Franks et al., 2017; Weisbecker et al., 2019; Mitchell et al., 2021). This can potentially complicate interpretations of the evolution of facial gracilisation and requires further examination. However, the predictability by which bone is expected to experience specific forces is considered a major factor regarding macroevolutionary changes to bone optimisation (Willie et al., 2019). The reductions to bone volume leading to more gracile morphology that we speak of would therefore occur on a macroevolutionary scale. These would not involve short-term effects of deposition or resorption, but instead selection for less bone on a genetic level, thus affecting interspecific variation in cranial gracilisation in a way that would be independent of developmental plasticity. As a simple contrast, a polar bear with a lifetime of feeding on more resistant foods will likely have thicker bone in regions of the skull that experience high levels of stress during biting, as demonstrated in the skulls of other mammals (e.g., Bouvier & Hylander, 1981; Mitchell et al., 2021), but it will not have a cranium more similar in proportions to the exceedingly robust cranium of the giant panda (*Ailuropoda melanoleuca*) that represents a product of long-term evolutionary change towards a particularly resistant diet (Figueirido et al., 2012; 2014).

### (2) Other selective advantages

The frequent occurrence of allometric facial gracility suggests that there are selective benefits associated with an elongate facial skeleton that may be more influential at larger sizes with relaxed bite force requirements. Yet, these benefits are almost certainly nuanced and differ for each taxon in question. An in-depth discussion on every family tested here is beyond the scope of this review; however, some examples of benefits to an elongate facial skeleton discussed across mammals include: a wider gape (Herring and Herring 1974; Greaves 1982; Dumont and Herrel 2003; Bourke et al. 2008; Slater & Van Valkenburgh, 2009; Williams et al., 2009; Figueirido et al., 2011; Hylander, 2013; Hennekam et al., 2020;), increased jaw closure speed (Preuschoft & Witzel, 2005; Slater et al., 2009; Van Valkenburgh & Slater, 2009), improved olfaction (Van Valkenburgh et al., 2014), accommodation of larger anterior teeth (Meachen-Samuels & Van Valkenburgh, 2009; Hylander, 2013; Tamagnini et al., 2017), improved forage selectivity (Gordon & Illius, 1988; Janis & Ehrhardt, 1988; Jarman and Phillips, 1989; Janis, 1995), enhanced cropping ability (Greaves, 1991; Codron et al., 2019; Dawson et al., 2021), the potential for improved vigilance in some prey species (Spencer, 1995; Mitchell et al., 2018), and housing long tongues (Greaves, 1991; Coombs, 1983; Freeman, 1995; 1998; Endo et al., 2007; Nogueira et al., 2009). Of course, reverse evolutionary scenarios (where species decrease in size over evolutionary time) also exist and we expect them to be subject to the same trade-off between bite force allometry and advantages of facial elongation. In these cases, smaller cranial sizes would involve selection for greater relative bite force over other benefits of elongated crania, allowing the exploitation of a more similar dietary range to larger related species. A smaller species of similar feeding ecology within a clade would therefore be expected to exhibit some evidence of non-size related adaptations for accommodating higher bite forces (Fig. 7B,E,F,H), resulting in a more stout facial morphology (see Kraatz & Sherratt, 2016; van der Geer et al., 2018; Alhajeri, 2021 for potential examples).

### (3) A secondary influence of negative braincase and orbit allometry

In Section II, we argued that the interpretation of facial proportions – particularly length – should be made independently of where the orbits are positioned. This is because assessments of longer or shorter face lengths could then be purely due to orbit position, rather than changes in facial proportions. However, even when facial proportions are not defined with reference to orbits, the orbits can still influence the interpretation of facial length due to their own hypoallometry and the fact that their position in the cranium depends on the similarly hypoallometric braincase, and their proximity to it (Fig. 3). The family of Bovidae shows that this effect indeed exists, and is an excellent example of how a taxon with larger ranges of cranial size exhibits substantial differences in orbit size and position as the braincase and orbits become relatively smaller in larger species (see Fig. 3 and Appendix C). This effect would be compounded if selection favours facial proportions that approach isometry (i.e., an ABC of approximately 1), leaving smaller, more posteriorly situated orbits in larger species as the predominant difference in facial morphology. However, the degree to which orbit and braincase hypoallometry contribute to assessments of facial gracilisation is likely small, because our landmark visualisations showed consistent braincase hypoallometry across all families that exhibited significant overall cranial allometry, regardless of whether the overall allometric pattern indicated gracility or stoutness at larger sizes. This suggests that braincase hypoallometry and associated relative position of the orbits is not a constraint on the evolution of relative length of the viscerocranium, but it should be considered in studies of individual taxa.

## VIII EXCEPTIONS: EVOLUTION DOESN’T ALWAYS PLAY BY THE RULES

Much of our above arguments assume that most mammals employ biting, particularly with the anterior dentition, as a primary jaw action. But this is of course not always the case, so that regularities or ‘rules’ of bite-related allometry cannot be expected to explain the evolution of all cranial shapes. There are many examples where biting to obtain and mechanically break down food is either not necessary, or assisted by alternative anatomy and behaviour. For example, strong lips, tongues, or trunks are frequently used by a range of mammalian herbivore taxa to acquire vegetation (Coombs, 1983; Mitchell et al., 2018; Williams, 2019), often bypassing the need for hard biting with the anterior dentition and limiting associations between bite force and food accessibility. In a similar fashion, much of primate food prehension, and sometimes initial processing, is assisted by forelimb manipulation and, occasionally, tool use (Heldstab et al., 2016). Macroevolutionary shifts in facial morphology for such groups are probably less influenced by selection for bite force optimisation (though not necessarily absent altogether) and are thus potentially able to respond more strongly to selection from alternative factors. Contrary to the predictions of our framework, there are also cases where low bite force demands might instead lead to more stout cranial dimensions. For example, the stout craniofacial anatomy of round-headed pilot whales and relatives (Globicephalinae) are adapted not for hard biting, but instead for suction feeding (Werth, 2006), and the effects of contrasting conditions of aquatic environments towards feeding mechanics in the context of CREA rightly deserve further investigation. Likewise, the extinct giant short-faced kangaroo (*Procoptodon goliah*) was the largest representative of the extinct Sthenurine kangaroos, and yet did not have a more gracile facial skeleton than smaller relatives. In fact, this species exhibited extreme reduction of the incisors and premaxillary region reflecting further shortening of the rostrum (see Prideaux et al., 2009). Prideaux (2004) suggested that the dietary yield per incisor bite would not have been efficient enough for such a large animal, and that vegetation would instead have been fed directly to the molars. Therefore, in some cases, rostrum length might exhibit hypoallometry when incisor biting is not an important part of food acquisition. However, it is important to note that for a group such as the sthenurine kangaroos that are often characterised by extremely robust facial skeletons, a perceptibly gracile morphology might have been unattainable. This leads us to discuss the potential limitations to morphological adaptation imparted by phylogenetic legacy.

Dollo’s Law of irreversibility states that an organism can never return exactly to a former macroevolutionary state, even if placed under identical former evolutionary selection regimes (Gould, 1970). Exactly how much facial gracility or stoutness can evolve before a morphology comparable to the ancestral condition can no longer be attained is difficult to answer and is probably unique to each clade in question. From a biomechanical perspective, there is less selective pressure for a stout-faced clade to become more gracile over evolutionary time than the reverse because, while a more gracile form will not as easily bite foods that require higher bite forces, a stout-faced animal can exploit both less resistant and more resistant resources. Therefore, in the absence of alternative selective pressures that would encourage a more elongate facial skeleton, such as those listed in Section VII, more stout proportions might be retained. In such cases, a stout cranium might not always be indicative of hard biting behaviours, but instead be a result of phylogenetic contingency. While some stout-faced ecomorphs could have arisen due to genetic bottlenecks or fallback foods during dietary stresses in their evolutionary history, subsequent diet shifts to more easily obtained and processed foods might not immediately result in a return to more gracile forms, if at all (see Liem’s Paradox; Liem, 1980; Robinson et al., 1998; Ungar et al., 2008). Thus, some stout ecomorphs might represent macroevolutionary “ratchets”, in which reversals from hyperspecialised cranial dimensions are rare, thereby increasing extinction risk when subjected to changing environmental conditions or competition with species that exhibit more gracile ecomorphs (Van Valkenburgh, 2007; Tseng & Wang, 2011; Tseng, 2013; Balisi, Casey, & Van Valkenburgh, 2018). In carnivorans, stout cranial proportions among larger extinct species are associated with heightened dietary specialisation, and a greater vulnerability of these morphotypes to extinction has been suggested (Holliday & Steppan, 2004; Van Valkenburgh et al., 2004; Van Valkenburgh, 2007). If stout facial proportions are attributable to hyperspecialisation for particular diets, and these morphologies are relatively immutable, metabolically expensive, and impractical in novel environmental conditions, this could possibly contribute to the prevalence of these ecomorphs among extinct groups that are not represented in tests for allometric facial elongation.

## IX WHERE DOES MAMMALIAN FACIAL SCALING FIT INTO OUR UNDERSTANDING OF CRANIAL ALLOMETRY?

For the many mammalian groups that have a strong reliance on biting with the anterior teeth as part of their niche demands, we have presented a functional mechanism governing patterns of facial scaling, based on biting biomechanics. However, as outlined above, there remain many instances where biting mechanics are less influential and facial scaling is instead sometimes a product of other adaptations. This strongly indicates that observed patterns of allometric facial scaling are not a product of a ubiquitous “rule” across all mammals, nor a product of a specific constraint, but are instead the combined result of multiple well-documented macroevolutionary phenomena involving ecology, behaviour, anatomy, and phylogenetic proximity that form the foundations of our proposed framework. The degree of influence and combined effects of these phenomena are undoubtedly unique to each taxon of interest and should be considered for analyses and interpretations going forward, rather than attributing results to any singular pattern. We therefore use this section to discuss how the framework and patterns of facial scaling might fit into a new, simple conceptualisation of allometry of the mammalian skeleton.

It is well established that a particular phenotype is the result of genetic, structural, functional, and environmental factors. Constructional Morphology is a conceptual framework that binds these factors together for an integrated understanding of the diversity of biological form (Seilacher, 1970; Thomas, 1979; Briggs, 2017). The framework involves three main pillars: the phylogenetic factor is a product of genetic heritage, the morphogenetic factor defines the contribution of growth and development processes, and the adaptive factor describes the niche-specific functional improvements obtained through natural selection. Interestingly, these three pillars naturally align with the three levels of allometry (Fig. 9; Thomas, 1979): Ontogenetic allometry is represented by individual growth, following a genetic blueprint determined by phylogenetic history (i.e., the phylogenetic factor); static allometry describes variation in size-related shape at a specific developmental stage, attributable to variation in genetics and environmental factors during growth (Lande, 1979; Voje et al., 2013) (i.e., the morphogenetic factor); and evolutionary allometry is influenced by size-related functional adaptation causing deviations to ontogenetic and static allometries (Voje, et al., 2013) (i.e., the adaptive factor). As detailed in the Introduction, these three pillars of allometry can influence each other in different and often complicated ways (Fig. 9) (Rensch, 1948; von Bertalanffy & Pirozynski, 1952; Lande, 1979; Voje et al. 2013), thereby acting together to generate allometric diversity in skeletal form.

**Figure 9:**
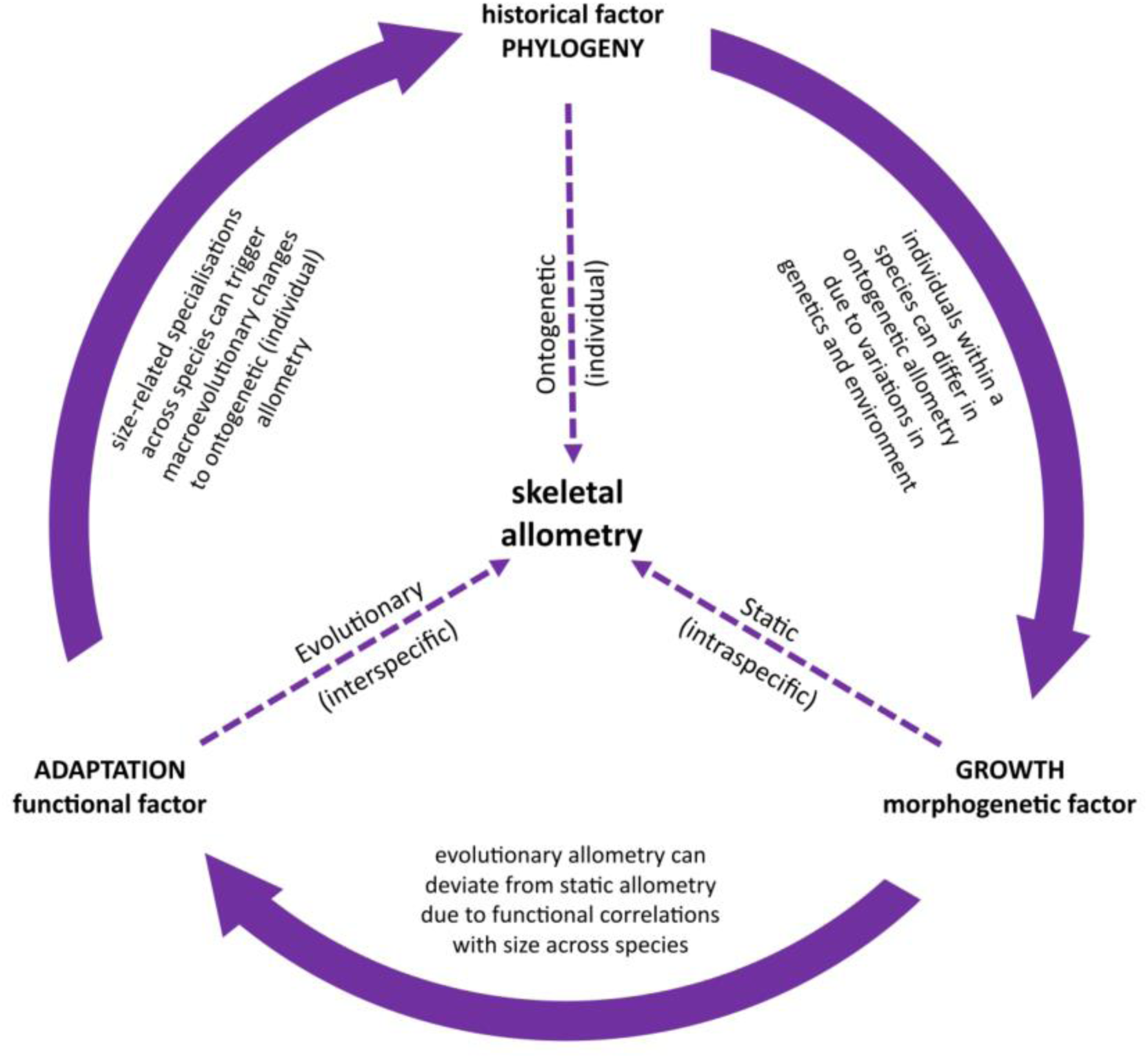
Skeletal allometry superimposed onto the Thomas’ (1979) Constructional Morphology framework.

While no mechanism has been identified to explain the pattern of hyperallometric facial elongation that is currently most discussed in the context of CREA (Cardini, 2019), it has been suggested to be related to heterochrony and to represent a size-related constraint on cranial diversity (Cardini & Polly, 2013). Through our investigation of 22 mammalian families, we found some evidence suggesting the presence of size-mediated constraints that are independent of phylogeny, but this was only found in seven of the 22 families. However, this could also be an artifact of how size is structured within these phylogenies. We found that braincase size was consistently smaller in larger species of all taxa with significant results for allometry, in agreement with Haller’s rule. But this was not coupled with patterns of facial gracilisation. Instead, the biomechanical principles we have outlined in this review, their close association with hyperallometric gracilisation, and the exceptions we have discussed, suggest that allometric variation related to facial elongation is dominated by adaptive responses to functional demands across size ranges, rather than a product of developmental constraint. It is certainly possible for developmental constraints, such as discussed for orbit size, to influence interpretations of facial elongation, but this can only occur within the bounds of functional selection. Therefore, the question of interspecific differences in “face length” is better answered by considering aspects of natural selection, placing patterns of allometric facial scaling among the adaptive factors influencing morphological diversity, with developmental constraints likely a secondary factor.

The interplay between allometry and adaptation is a venerable field which can be substantially improved through better understanding of the processes behind the evolution of allometric patterns. This is important because of the fundamental importance of size variation in the evolution of mammalian diversity. For example, Huxley (1924) noted that advantages to larger body sizes might explain the prevalence of evolution for larger species across most mammalian groups. Size variation indeed provides the basis for a greater phenotypic plasticity among mammals, facilitating the arrival of novel adaptive morphologies (Björkland, 1996; Bubadué et al., 2022). The capacity for adaptive innovation of the cranium to expand beyond the confines of biomechanical constraints might impart selective advantages to increased body sizes, alongside other advantages suggested to promote Cope’s rule (Kingsolver & Pfennig, 2004 and references therein). This could be an important component in the allometric “line of least resistance” that facilitates evolutionary adaptation in the mammalian cranium (Huxley, 1924; Marroig and Cheverud 2005; Shirai & Marroig, 2010). It has been argued previously that the CREA pattern follows an evolutionary line of least resistance (Cardini & Poly, 2013; Cardini, 2019; Rhoda et al., 2023), to which we agree. However, we suggest that this might be largely mediated by bone moderation and enhanced cranial functions at larger sizes (see Section VII), rather than extensions of ontogenetic or static allometries. With a better understanding of the interplay between adaptations and constraints, and the realisation that this relationship can change along body size ranges, future research will have an excellent starting point for an integrated view of how allometry-driven diversity arises across the vast range of mammalian body sizes.

## X CONCLUSION

1. Facial elongation with increasing size has been argued as a prevalent allometric constraint across mammals, and possibly vertebrates in general. However, we showed that the definition of CREA has been ambiguous, with multiple interpretations of what the pattern involves. We have addressed this by refining the conditions of the definition, suggesting that a “long face” is represented, not necessarily solely by elongation of the rostrum, but by a general increase in gracility of the facial skeleton in larger species. This can arise through multiple adaptive pathways. We presented a simple procedure for testing cranial evolutionary allometry that does not conflate the findings with Haller’s rule (i.e. brain hypoallometry) and have shown that the pattern is not ubiquitous, but common.
2. Predictions for the hyperallometric gracilisation pattern in the cranium are influenced by the degree of ecological congruence across the species tested and which species are included. This means that the taxonomic scale of a given sample is arbitrary and most often depends on the range of ecological traits of the taxon being examined. The prevalence of a phylogenetic signal of size among mammalian taxa means that factoring phylogenetic relatedness into models can frequently obscure evolutionary allometry when size correlates with relatedness, and so morphological variation throughout a clade should also be examined without phylogenetic correction.
3. The biomechanical principles behind various cranial adaptations for hard biting dictate that a “shorter face” among mammals often represents an example of many-to-one mapping, in that multiple adaptive pathways can occur that achieve similar functions towards producing and withstanding greater relative bite forces. From this, it also follows that larger species exhibiting facial gracilisation reflect an evolutionary sacrifice of relative bite force or the ability to withstand the stresses associated with harder biting.
4. The reason why hyperallometric gracilisation tends to occur more often among closely related species appears to be phylogenetic niche conservatism, where more closely related species are likely to have more similar absolute bite force requirements. Gracile cranial proportions are the more common outcome among larger animals because they often consume foods of similar mechanical resistance to smaller relatives, but with less relative mechanical effort, permitting optimisation of bone deposition or the accommodation of alternative selective pressures that encourage facial elongation. For more variable dietary ranges within a clade, as long as the rate of increase in bite force requirements remains lower than the rate of increase in bite force capacity that naturally comes with greater body size, facial gracility is a frequent outcome. A reverse pattern can therefore be observed if larger species occupy different niches requiring relatively greater bite forces than their size would normally accommodate. Hypoallometry of the orbits and braincase are likely to be a secondary constraining factor that might also result in interpretations of facial elongation under some conditions.
5. Taken together, our theoretical framework suggests that patterns relating to proportions of the viscerocranium, or “face length”, are less a result of developmental constraints, and instead dominated by functional trade-offs that can shift in importance along size ranges. Increasing body size can therefore potentially facilitate evolutionary adaptation and diversification. The ways by which macroevolutionary changes to body size can mediate relationships between adaptation and constraint present an interesting and important line of consideration for future investigation.

## XI ACKNOWLEDGEMENTS

We thank all those who uploaded surface scans to Morphosource for use in our figures: Sharon Grant, Serjoscha Evers, Carol Spencer, Joseph Tyler, Douglass Rovinsky, Justin Adams, Edwin Dickinson, and Thomas Macrini and Jessica Maisano (DigiMorph.org and NSF IIS-9874781) for their helpful assistance. We would like to personally thank all the researchers who freely uploaded their data with their studies. Without your academic collegiality, this work would not be possible. VW was supported by Australian Research Council (ARC) Future Fellowship FT180100634, VW and DRM were supported by the ARC Centre of Excellence for Australian Biodiversity and Heritage CE170100015, and ES was supported by ARC Future Fellowship FT190100803.

