## Appendix C for "Facing the facts: Adaptive trade-offs along body size ranges determine mammalian craniofacial scaling"

**Overview**

- The centroid size index (CSI) provides a standardised measure of cranial size disparity within the taxon. We obtain this by dividing the largest centroid size by the smallest.
- Each family has results of ordinary least squares regression (OLS) test for evolutionary allometry, principal component analysis (PCA), and phylogenetic generalised least squares regression (PGLS) test for evolutionary allometry, as well as significant correlations of PCs with size, and phylogenetic signal of cranial size.
- Only figures of relevance to either cranial size or facial gracility are presented.
- All analyses presented here were implemented with the geomorph R package in R Statistical Environment (see Appendix B). The script and data are available on Github (https://github.com/DRexMitchell/Mitchell-etal-facial-scaling).

### Dasyuridae

The Dasyuridae data includes 16 species. The largest centroid size divided by the smallest gives the largest centroid size scaling index of all families tested, at 8.640. This dataset was the only original creation of data for this project, generated by the authors using a landmark protocol of fixed 3D landmarks derived from Viacava et al., 2020). There is a significant allometric signal ($R^2$ = 0.330, p=0.001) (Fig. S1A). The predicted shape changes suggest larger species have a smaller braincase, but there is no obvious relation to rostrum length (Fig. S1B) (see also Viacava et al., 2020). The first principal component (PC1) is correlated with size (p<0.001) and includes braincase size (Fig. S1C,D). PC2 is not correlated with size, but is associated with cranial width, mostly due to the large, bone cracking morph of the Tasmanian Devil. Allometry remains significant after phylogenetic correction ($R^2$=0.415, p=0.016), with a significant phylogenetic signal of centroid size (p=0.008). While PC2 is concerned with relative skull width and evidences a clade-wide trend towards a narrower skull with increased size, larger species are predicted to have a smaller braincase and a generally wider cranium (Fig. S1E), contradicting predictions for hyperallometric gracilisation. This is due to the broad-faced, bone-cracking morph of the largest species, the Tasmanian devil (*Sarcophilus harrisii*), positioned as a sister taxon to the larger-bodied quolls (*Dasyurus* spp.) (Fig. S2), and positioned entirely outside of the narrowing trend in PC2.

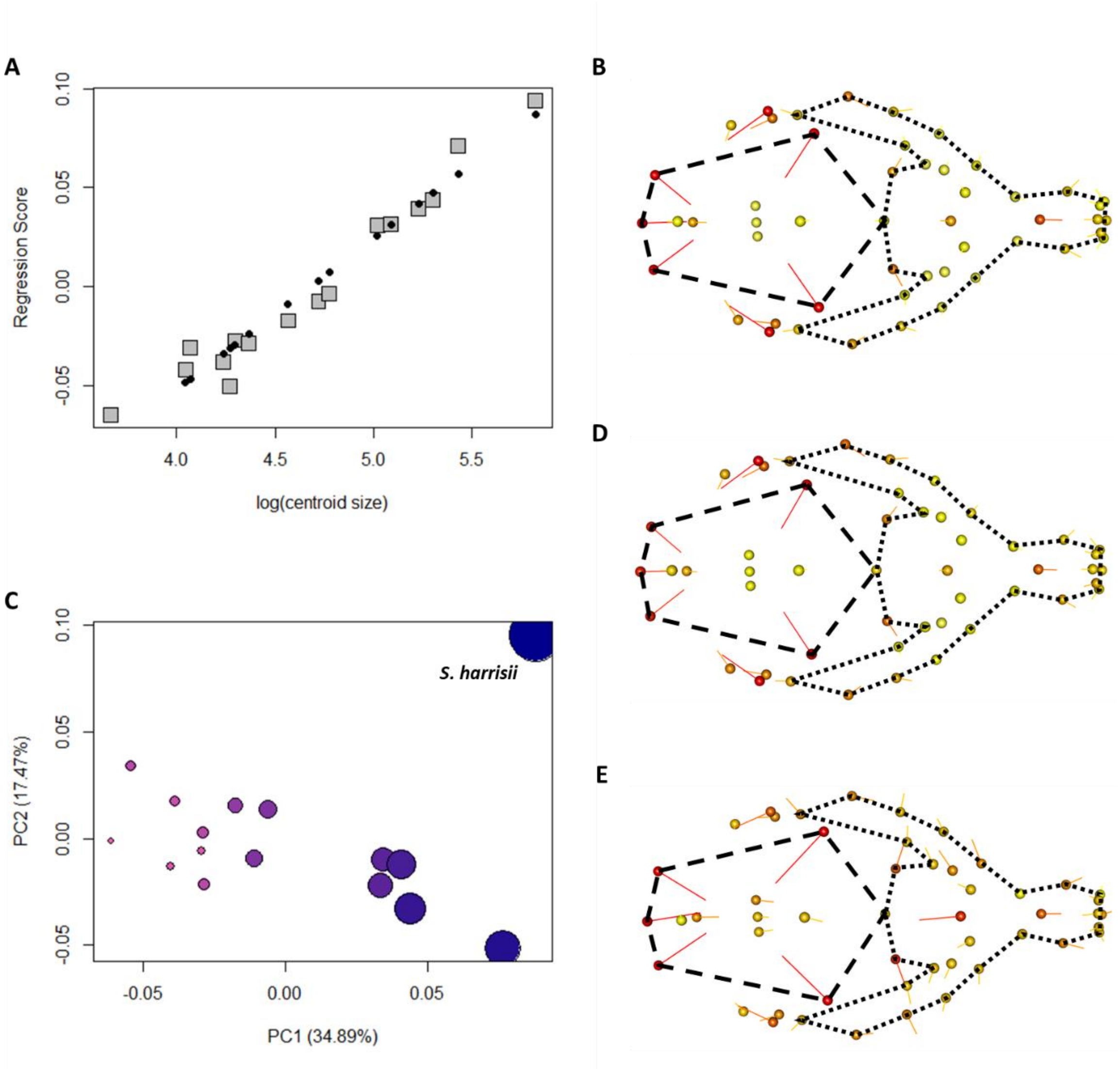

**Figure S1: (A) Shape regression score over log(centroid size) indicates allometric relationship (black dots = PredLine). (B) The predicted shape of the PredLine for the ordinary least squares allometry test, (C) plot of the first two principal components (orb size represents cranial centroid size), (D) shape variation defined by PC1 (orbs represent the minimum extreme, line tips are the maximum) (E) Allometric shape changes predicted after phylogenetic correction (PGLS) (orbs represent smaller sizes). Dashed lines = braincase, dotted lines = face.**

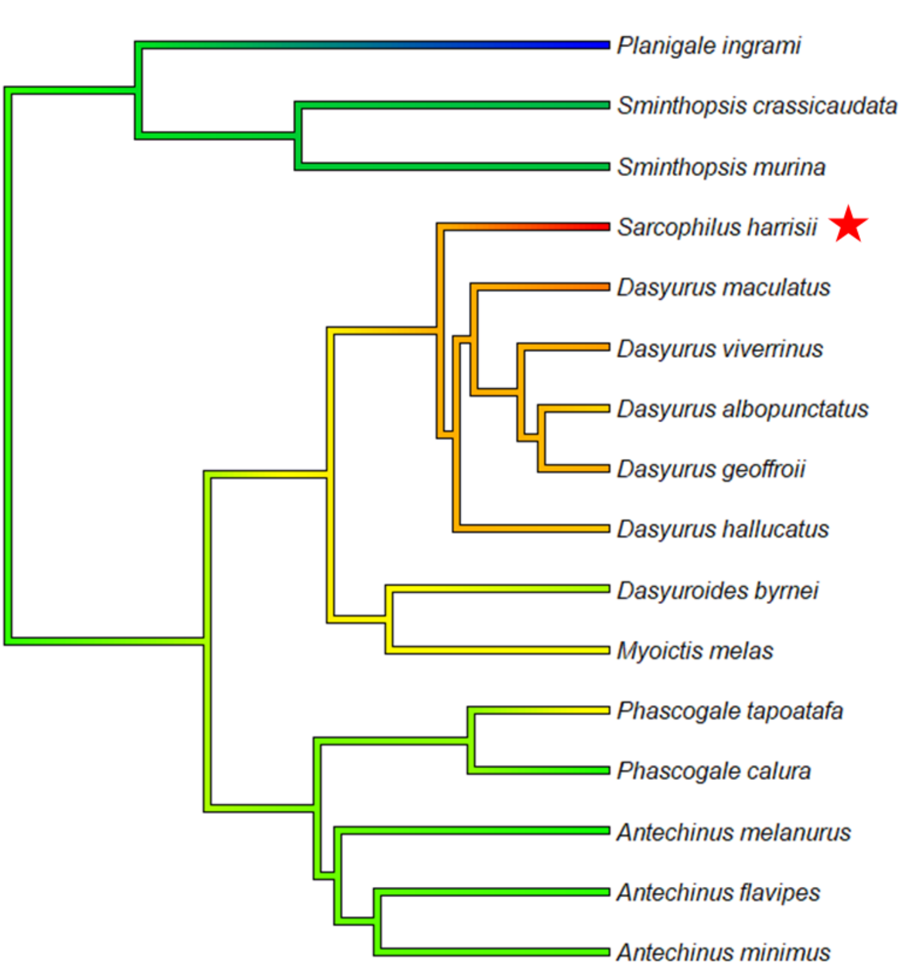

**Figure S2: Phylogeny of the Dasyuridae, coloured by cranial centroid size from smallest (blue) to largest (red). The bone-cracking skull morphology of the large-bodied Tasmanian Devil (*S. harrisii*; red star) is positioned as a sister-taxon to the genus *Dasyurus*, which has likely caused the reverse pattern towards facial stoutness predicted for this clade.**

### Macropodidae

The data comes from Mitchell et al. (2018), comprising 12 species with a centroid size scaling index of 2.44. There is a significant allometric signal ($R^2$ = 0.273, p=0.002) (Fig. S3A). The predicted shape changes suggest larger species have a smaller braincase, and projection of the anterior dentition, supporting increased face length in this family (Fig. S3B). The first principal component (PC1) is correlated with size (p=0.003) and includes braincase size and projection of the anterior dentition in larger species, consistent with the ordinary least squares predictions (Fig. S3C, D). There is a significant phylogenetic signal of centroid size (p=0.004), but allometry is not significant after phylogenetic correction. The phylogeny of the species included has two monophyletic clades, the *Petrogale* group in which the species are relatively similar in size, and the *Macropus* group in which there is a phylogenetic trend of increasing size with more recent divergences (Cope's Rule). However, while the family does tend towards rostrum elongation with increased size (Cardini et al., 2015), the tree-kangaroos (*Dendrolagus* spp.) occupy the minimum of PC1 (Fig. S3C) and are clearly more stout-faced than all other species despite being medium-sized (Fig. S4). This is likely associated with a more mechanically variable diet of browse, while the largest species feed on grasses with more consistent mechanical properties (Mitchell et al., 2018).

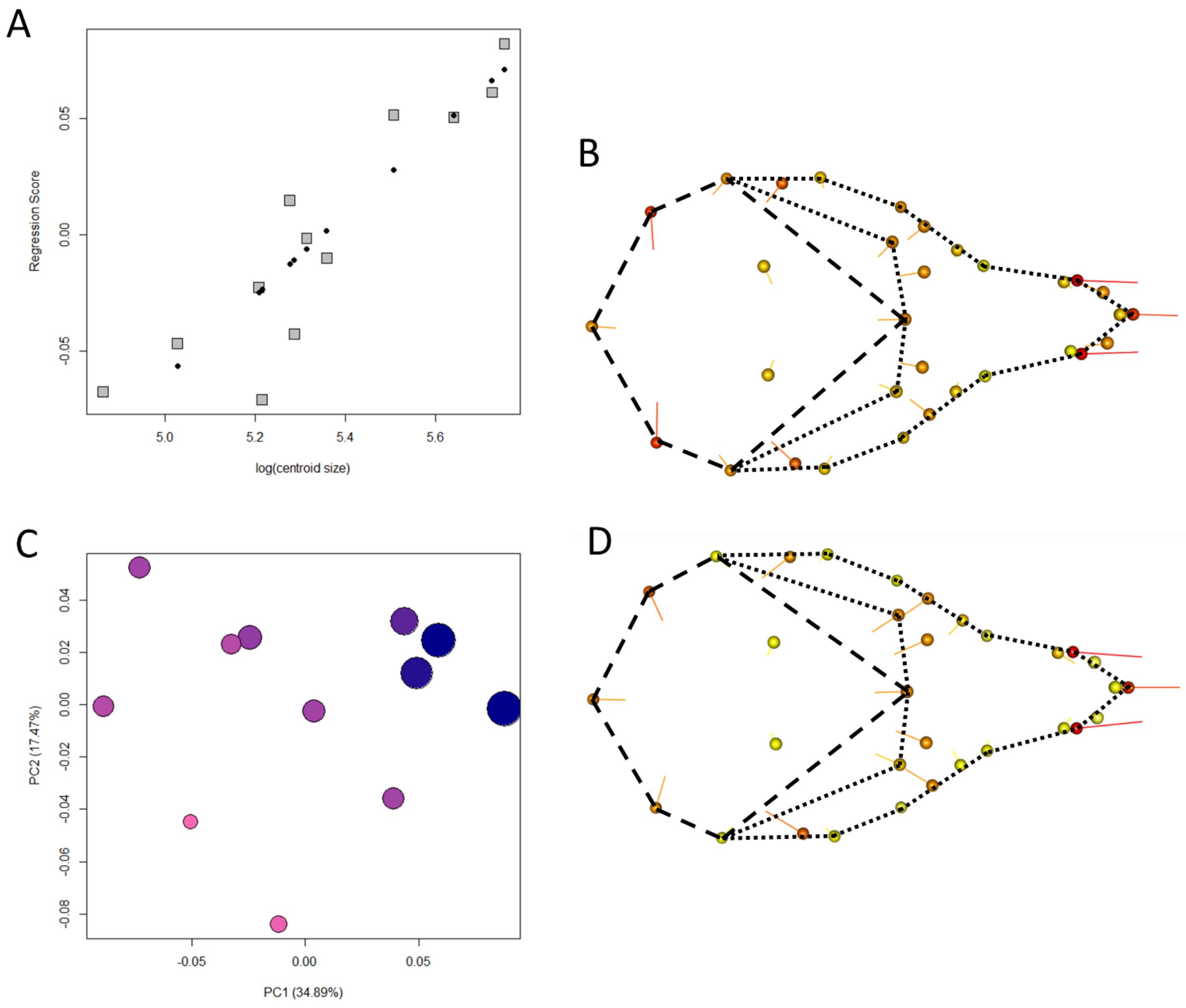

**Figure S3: (A) Shape regression score over log(centroid size) indicates allometric relationship (black dots = PredLine). (B) The predicted shape of the PredLine for the ordinary least squares allometry test, (C) plot of the first two principal components (orb size represents cranial centroid size), (D) shape variation defined by PC1 (orbs represent the minimum extreme, line tips are the maximum). Dashed lines = braincase, dotted lines = face.**

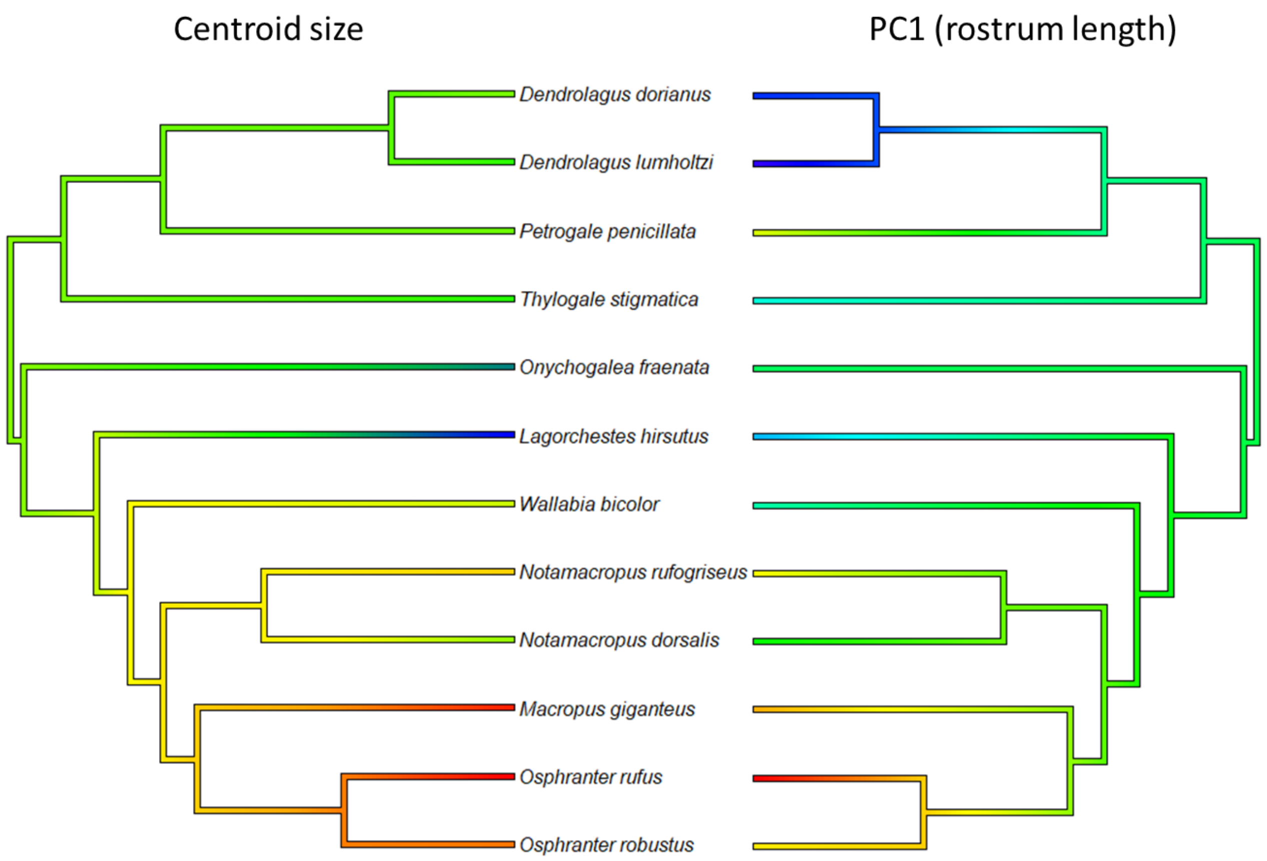

**Figure S4: Phylogeny of the Macropodidae, coloured by cranial centroid size from smallest (blue) to largest (red).**

**Leporidae**

The data comes from Kraatz & Sherratt (2016), comprising 20 species with a centroid size scale of 1.804. There is a significant allometric signal ($R^2$ = 0.137, p=0.016) (Fig. S5A). The predicted shape changes suggest larger species have a smaller braincase, and projection of the rostrum, supporting increased face length (Fig. S5B). PC1 is not correlated with size, but PC2 is (p=0.008) and is defined by braincase size and projection of the rostrum in larger species (Fig. S5C, D). There is a significant phylogenetic signal of centroid size (p=0.011), but allometry is not significant after phylogenetic correction. There is some indication of Cope's Rule in the phylogeny (Fig. S6), with the smaller species tending to be sister taxa to larger-bodied, more recently diversified monophyletic groups. As per Kraatz & Sherratt (2016), allometry of facial proportions appears to be largely driven by the smallest pygmy species, *Brachylagus*, which has a more stout viscerocranium.

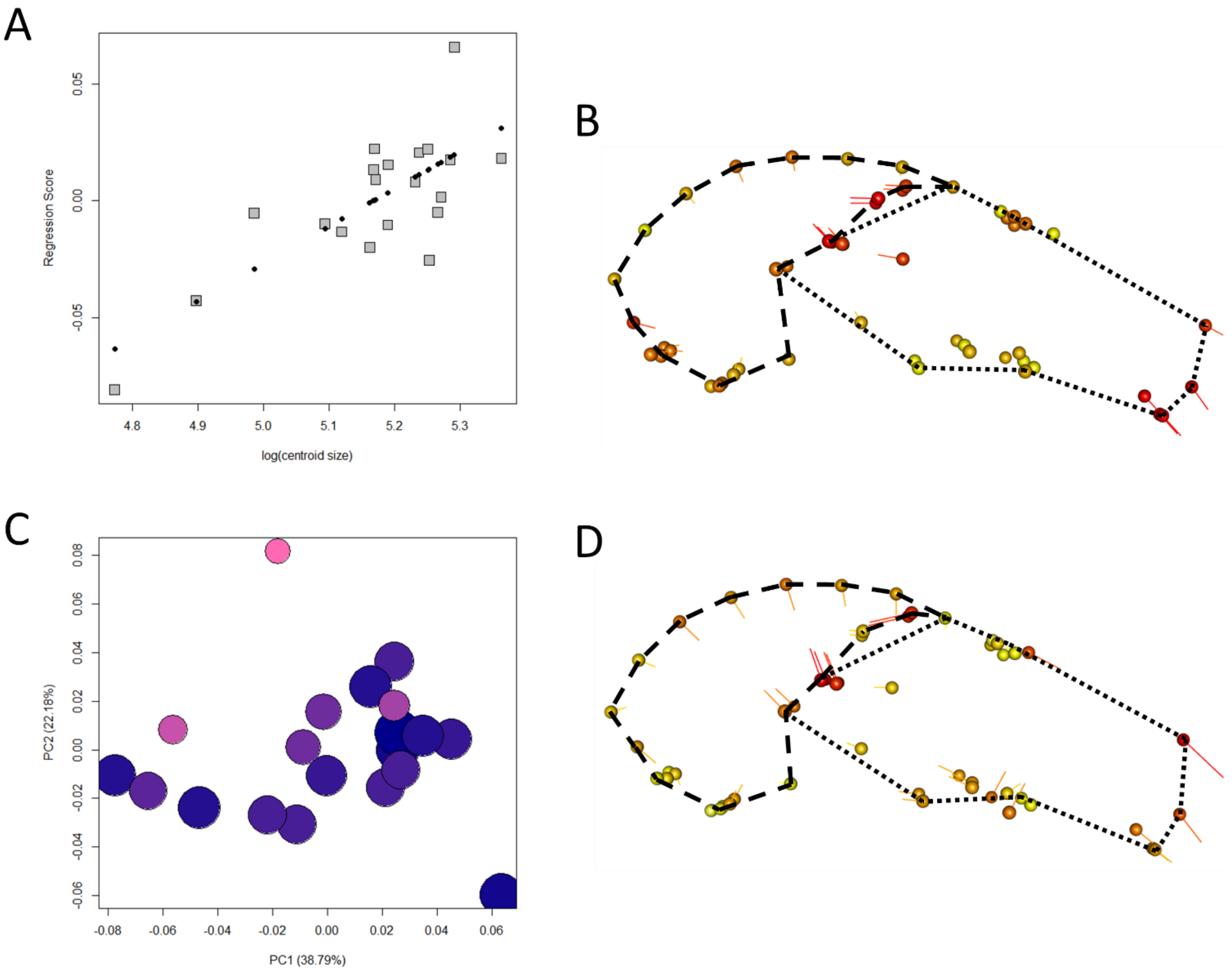

**Figure S5: (A) Shape regression score over log(centroid size) indicates allometric relationship (black dots = PredLine). (B) The predicted shape of the PredLine for the ordinary least squares allometry test, (C) plot of the first two principal components (orb size represents cranial centroid size), (D) shape variation defined by PC2 (orbs represent the minimum extreme, line tips are the maximum) (E) Allometric shape changes predicted after phylogenetic correction (PGLS) (orbs represent smaller sizes). Dashed lines = braincase, dotted lines = face.**

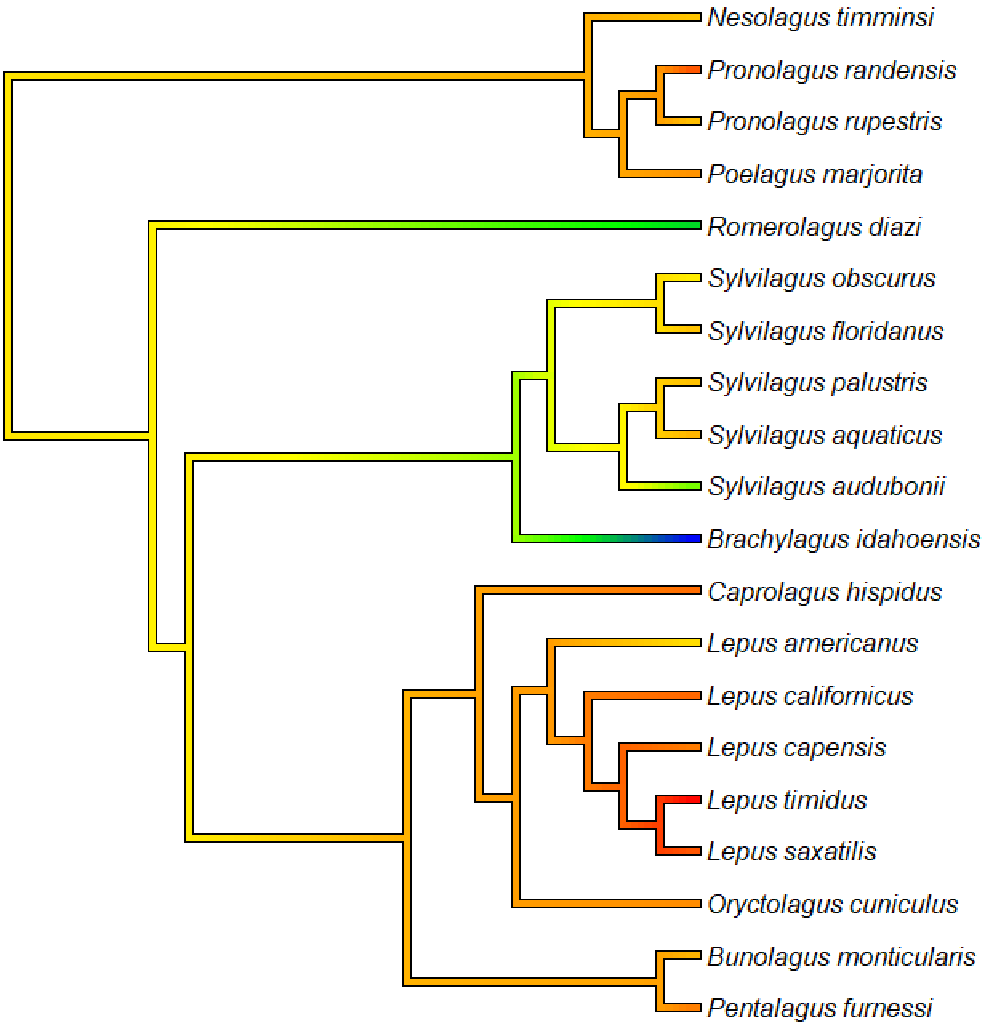

**Figure S6: Phylogeny of the Leporidae, coloured by cranial centroid size from smallest (blue) to largest (red).**

**Muridae**

This data comes from Marcy et al. (2020), comprising 37 species with a centroid size range of 3.217. Allometry was significant ($R^2$ = 0.448, p=0.001) (Fig. S4A), with larger species predicted to have relatively smaller braincases and an elongated rostrum (Fig. S4B). The first principal component (Fig. S4C) is significantly correlated with centroid size (p<0.001) and is defined by braincase size and rostrum depth and projection (Fig. S4D). Allometry remains significant after phylogenetic correction ($R^2$=0.160, p=0.001). Larger species are expected to exhibit smaller braincases and an elongated rostrum (Fig. S4E) supporting the pattern of hyperallometic gracilisation on all fronts, with a significant phylogenetic signal of centroid size (p=0.001). A comparison of cranial centroid size and relative face length across the phylogeny indicates high similarity between the two features, despite some differences in absolute values (Fig. S8).

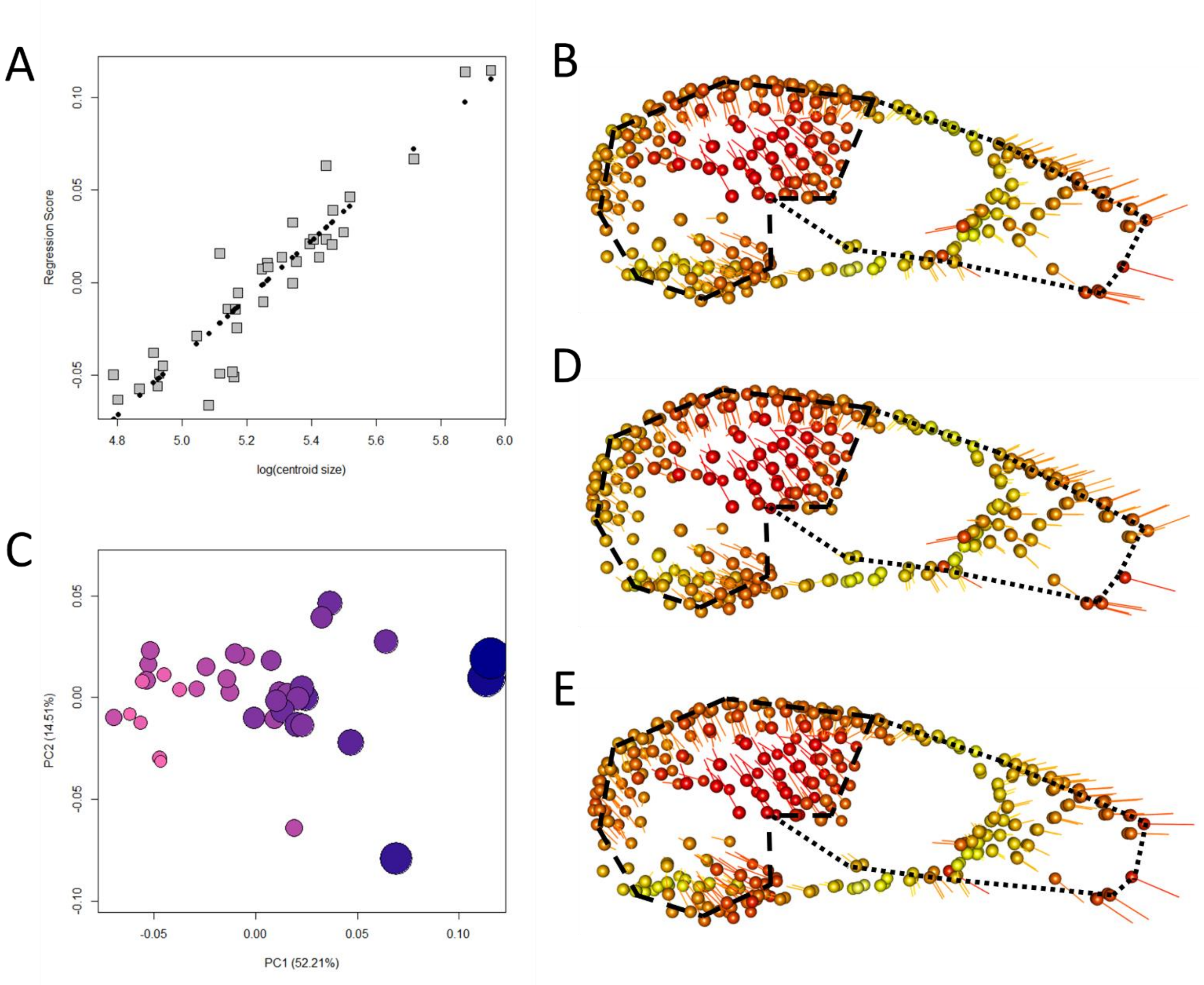

**Figure S7: (A) Shape regression score over log(centroid size) indicates allometric relationship (black dots = PredLine). (B) The predicted shape of the PredLine for the ordinary least squares allometry test, (C) plot of the first two principal components (orb size represents cranial centroid size), (D) shape variation defined by PC1 (orbs represent the minimum extreme, line tips are the maximum) (E) Allometric shape changes predicted after phylogenetic correction (PGLS) (orbs represent smaller sizes). Dashed lines = braincase, dotted lines = face.**

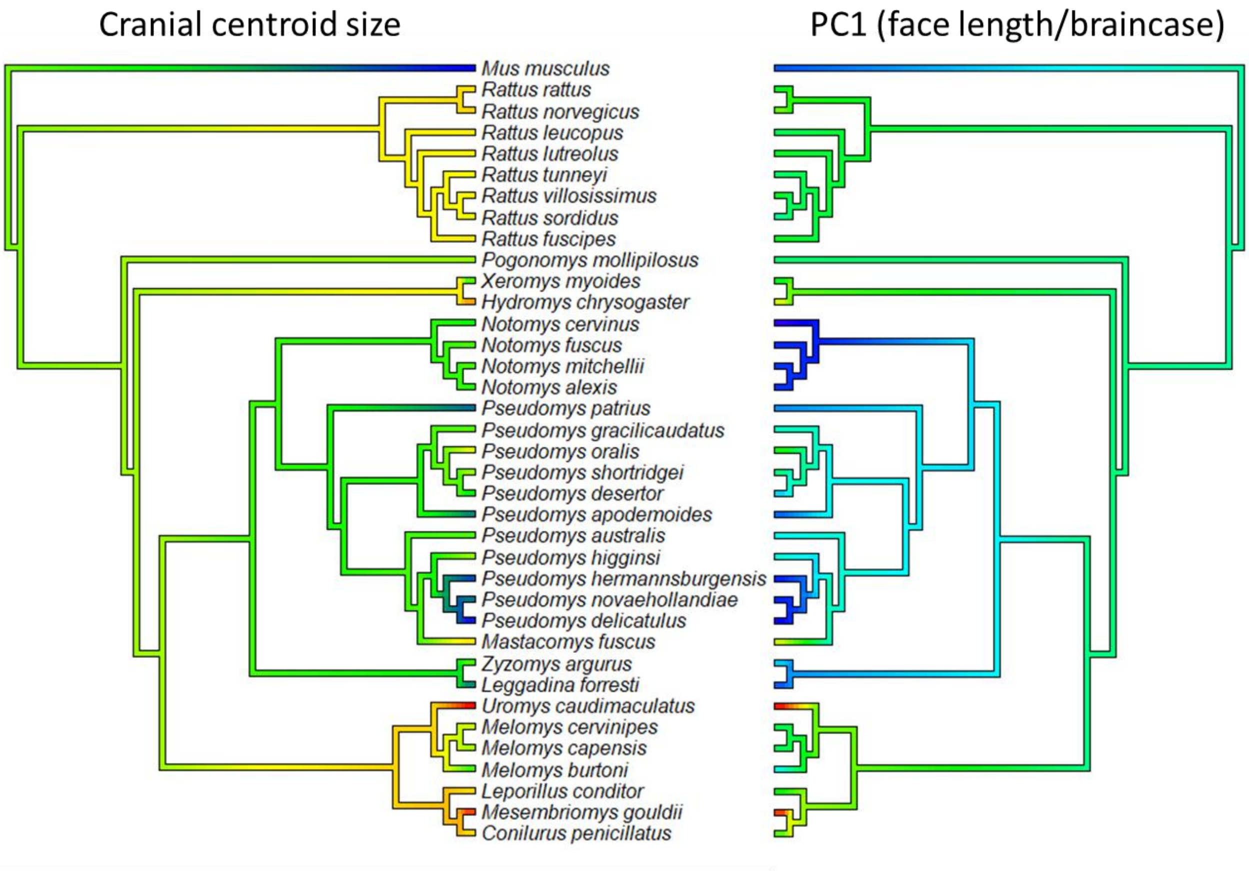

**Figure S8: Phylogeny of the Muridae, coloured on the left by cranial centroid size from smallest (blue) to largest (red), and on the right by PC1 representing relative rostrum length from shortest (blue) to longest (red).**

**Pteropodidae**

The Chiroptera data comes from Arbour et al. (2019). The Pteropididae comprises 24 species with a centroid size scaling index of 3.60. Allometry was significant ($R^2$ = 0.279, p=0.001) (Fig. S9A), with larger species predicted to have relatively smaller braincases and an elongated rostrum (Fig. S9B). The first principal component (Fig. S9C) is significantly correlated with centroid size (p<0.001) and is defined by rostrum depth and projection, with less associations with braincase size (Fig. S9D). However, this is explained by the second PC which is also correlated with size (p=0.001) and represents braincase size and zygomatic arch depth. These two first PCs have been noticeably combined in the allometric prediction. Allometry is no longer significant after phylogenetic correction, but there is a significant phylogenetic signal of centroid size (p=0.010). Comparisons of cranial centroid size and relative face length across the phylogeny indicate high similarity between the features, with the smallest species having the shortest faces and the largest species having the longest (Fig. S10). However, there are some noteworthy exceptions among the smaller species evidenced by discrepancies between the two phylogenies. The shorter-faced common tube-nosed fruit bat (*Nyctimene albiventer*) representing the minimum of PC1 and the longer-faced long-tongued fruit bat (*Macroglossus sobrinus*) located towards the maximum of PC1 (Fig. S9C). While *M. sobrinus* is nectarivorous, *N. albiventer* is known to incorporate hard fruits into its diet without needing to alter biting strategies during their consumption (Dumont & O'Neal, 2004). One very small species, *Syconycteris australis,* has a mid-range relative facial length (Fig. S9C) and feeds mostly on pollen and nectar.

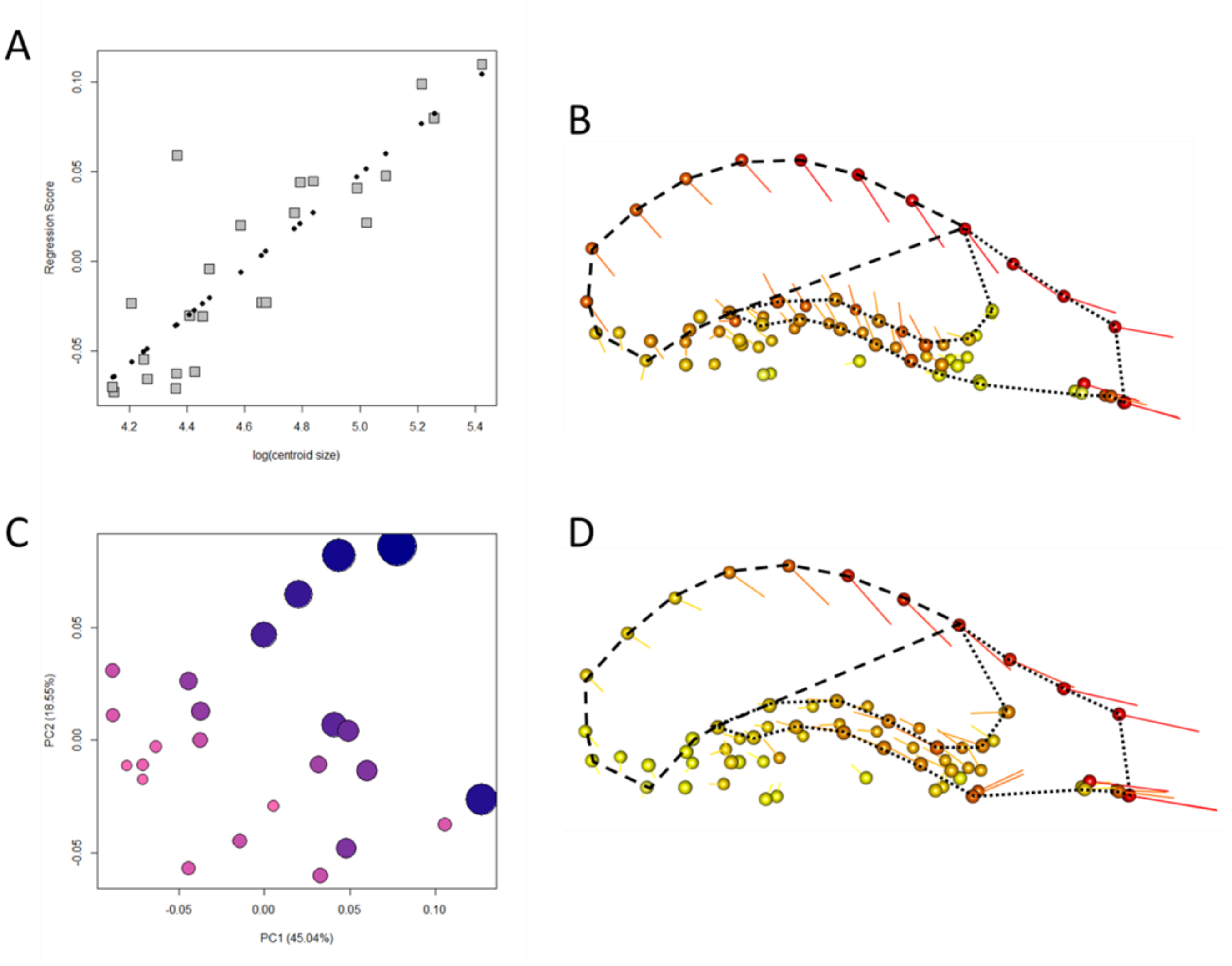

**Figure S9: (A) Shape regression score over log(centroid size) indicates allometric relationship (black dots = PredLine). (B) The predicted shape of the PredLine for the ordinary least squares allometry test, (C) plot of the first two principal components (orb size represents cranial centroid size), (D) shape variation defined by PC1 (orbs represent the minimum extreme, line tips are the maximum). Dashed lines = braincase, dotted lines = face.**

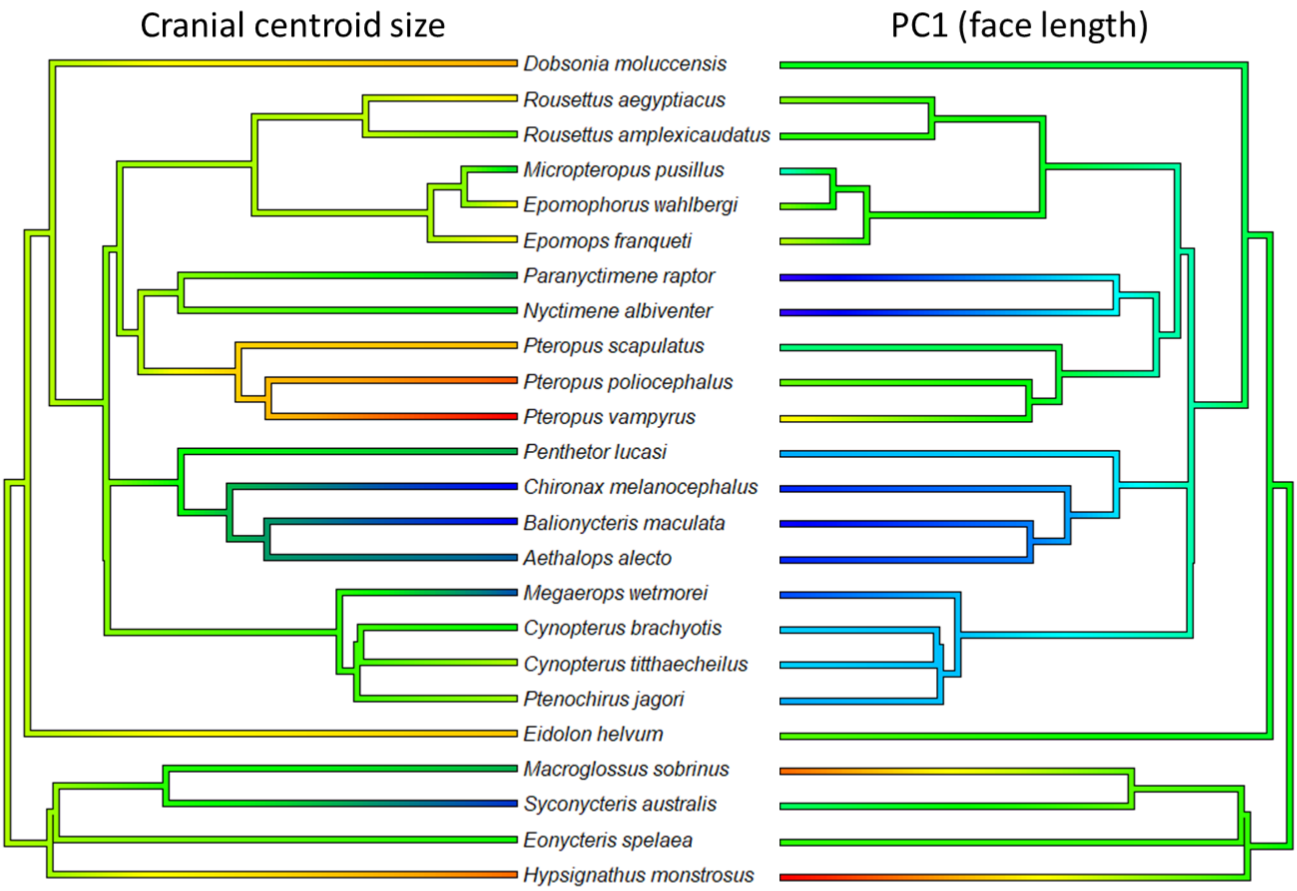

**Figure S10: Phylogeny of the Pteropodidae, coloured on the left by cranial centroid size from smallest (blue) to largest (red), and on the right by PC1 representing relative rostrum length from shortest (blue) to longest (red).**

### Emballonuridae

The Emballonuridae data includes 18 species with a cranial centroid size scaling index of 2.235. There is a significant allometric signal ($R^2$ = 0.223, p=0.001) (Fig. S11A). The predicted shape changes suggest larger species have a smaller braincase and more dorsally positioned premaxillae, but no facial changes in support of hyperallometric gracilisation (Fig. S11B). PC1 is correlated with size (p=0.002) (Fig. S11C) and is largely explained by braincase size and depth of the premaxillae (Fig. S11D). There is no significant allometry after phylogenetic correction, but there is a significant phylogenetic signal of centroid size (p=0.001).

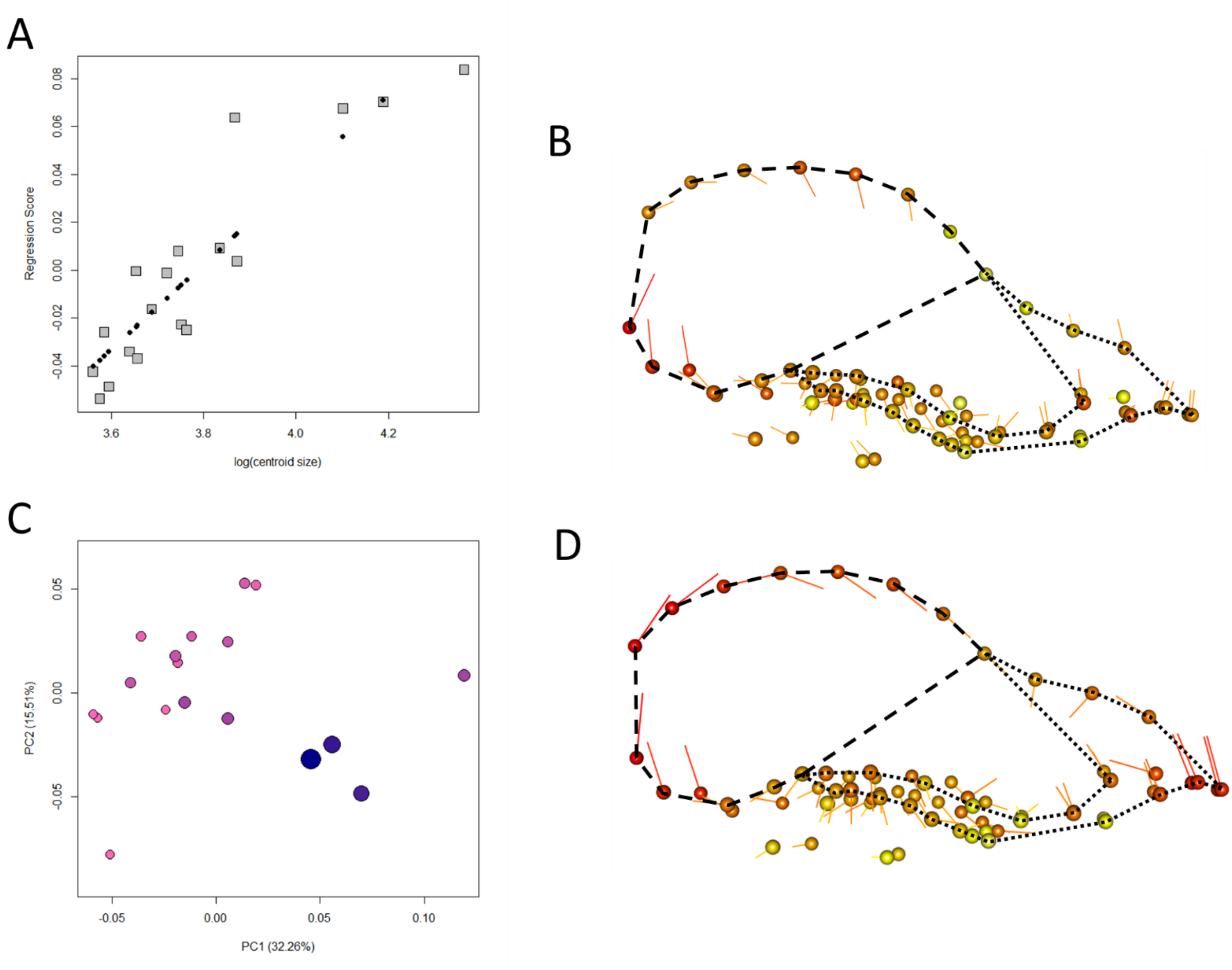

**Figure S11: (A) Shape regression score over log(centroid size) indicates allometric relationship (black dots = PredLine). (B) The predicted shape of the PredLine for the ordinary least squares allometry test, (C) plot of the first two principal components (orb size represents cranial centroid size), (D) shape variation defined by PC1 (orbs represent the minimum extreme, line tips are the maximum). Dashed lines = braincase, dotted lines = face.**

### Rhinolophidae

The Rhinolophidae comprises 28 species with a centroid size scaling range of 1.92. Allometry was significant ($R^2$ = 0.185, p=0.001) (Fig. S12A), with larger species predicted to have relatively smaller braincases and a deeper rostrum, inconsistent with predictions for gracility (Fig. S12B). However, the first principal component (Fig. S12C) is significantly correlated with centroid size (p<0.001) and is defined by rostrum projection with increased size (Fig. S12D, S13). Therefore, the smallest species occupying the mid-range of PC1, such as *R. monoceros*, *R. pusillus*, and *R. hipposideros*, are likely confounding predictions for hyperallometric gracilisation. Allometry is no longer significant after phylogenetic correction, but there is a significant phylogenetic signal of centroid size (p=0.001). As for the Emballonuridae, the Rhinolophidae are insectivorous, however, most studies of diet have only reported proportions of insect orders found in scats. While an influence of echolocation on face length is possible, two species with known differences in echolocation frequency and evidenced resource partitioning, *R. macrotis* and *R. Lepidus* (Shi et al., 2009), both occupy the mid-range of PC1. Thicker insect cuticles are more difficult to break down (Evans et al., 2005) and the proportions of specific species of more resistant insects, such as some species of Coleoptera, might therefore play a role in facial morphology (see Santana et al., 2012 regarding phyllostomid bats and Freeman, 1979 regarding molossid bats).

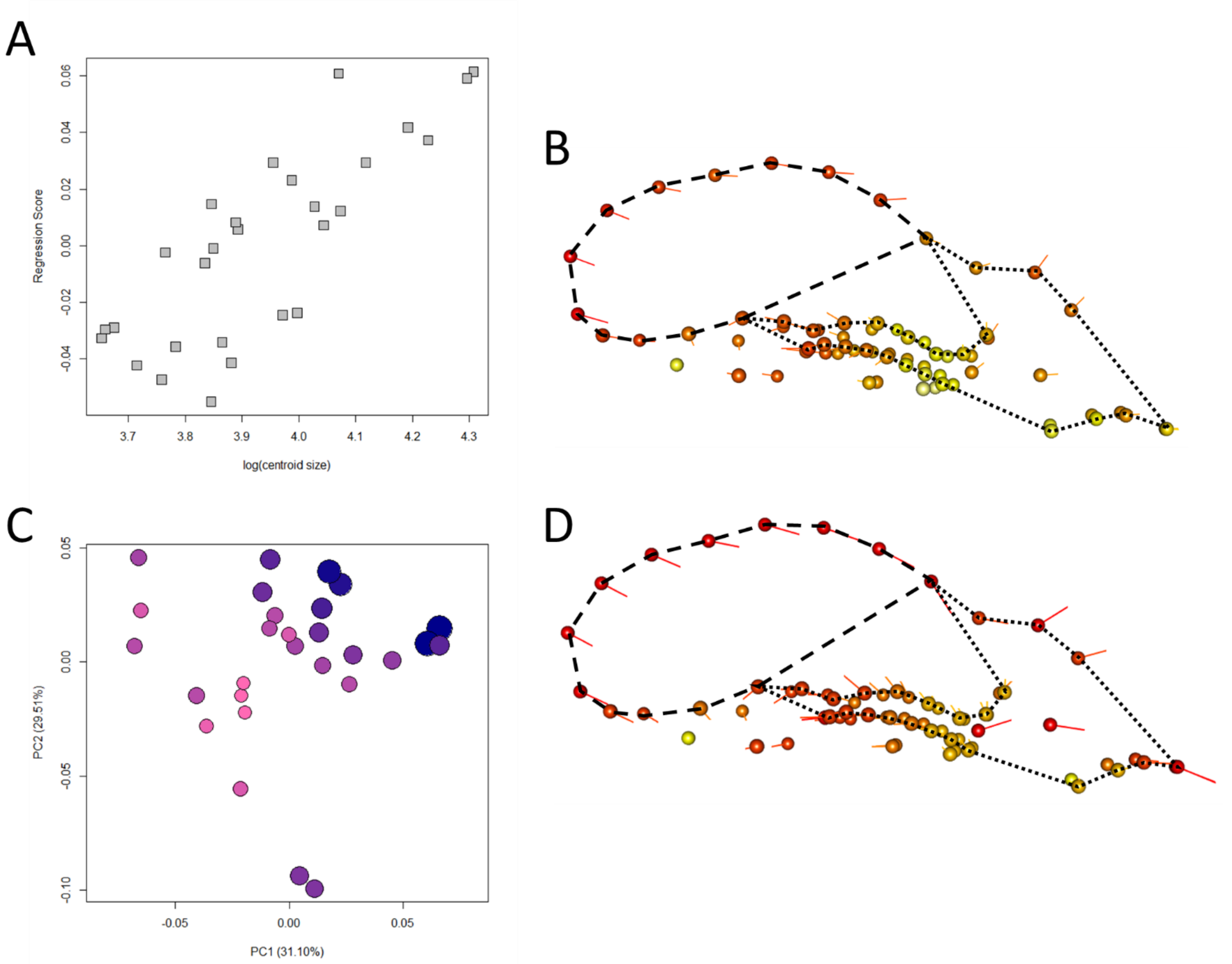

**Figure S12: (A) Shape regression score over log(centroid size) indicates allometric relationship (black dots = PredLine). (B) The predicted shape of the PredLine for the ordinary least squares allometry test, (C) plot of the first two principal components (orb size represents cranial centroid size), (D) shape variation defined by PC1 (orbs represent the minimum extreme, line tips are the maximum). Dashed lines = braincase, dotted lines = face.**

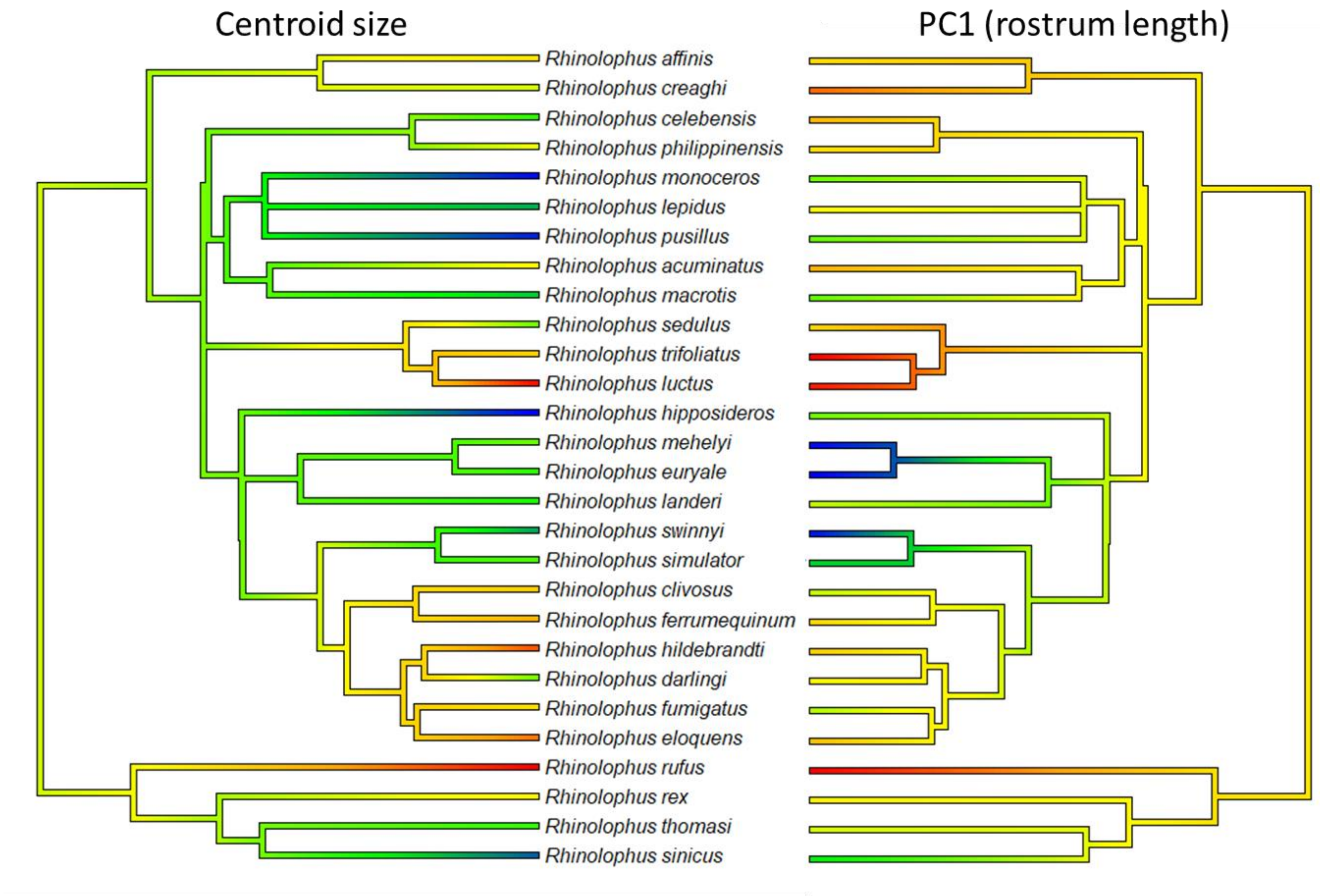

**Figure S13: Phylogeny of the Rhinolophidae, coloured on the left by cranial centroid size from smallest (blue) to largest (red), and on the right by PC1 representing relative rostrum length from shortest (blue) to longest (red).**

### Phyllostomidae

The Phyllostomidae comprises 30 species with a centroid size scaling index of 2.88. There was no significant allometry detected (Fig. S14A). The largest and smallest species are morphologically average within the shape space (Fig. S14B), but the first principal component does represent relative face length (Fig. S14C). There is no significant allometry after phylogenetic correction, but there is a significant phylogenetic signal of centroid size (p=0.018). There is no relationship between cranial size and facial gracility (Fig. S15). All size-related shape in this group appears to be overwhelmed by the extraordinary diversity in diet-related facial adaptations at similar size ranges, from long-faced nectarivorous species at the maximum of PC1 to the extremely short-faced seed predators at the minimum (Fig. S15). This is a highly studied family, given their diversity in cranial form (e.g., Nogueira et al., 2009; Santana et al., 2012; Dumont et al., 2014; Rossoni et al., 2017; Hedrick & Dumont, 2018). Dumont et al. (2014) suggest that selection for mechanical advantage in association with diet has determined much of the cranial diversity in this family.

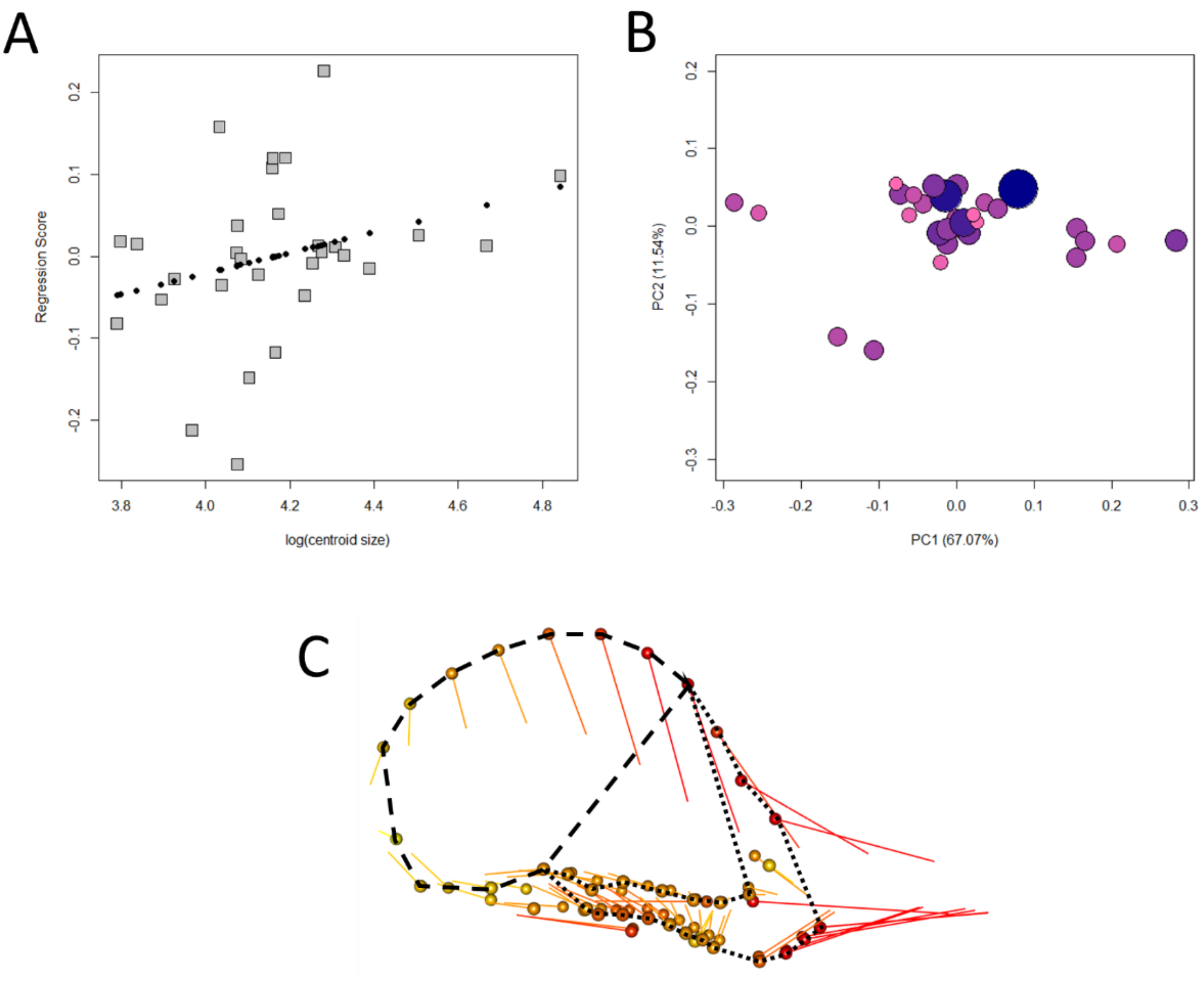

**Figure S14: (A) Shape regression score over log(centroid size) (black dots = PredLine). (B) plot of the first two principal components (orb size represents cranial centroid size), (C) shape variation defined by PC1 (orbs represent the minimum extreme, line tips are the maximum). Dashed lines = braincase, dotted lines = face.**

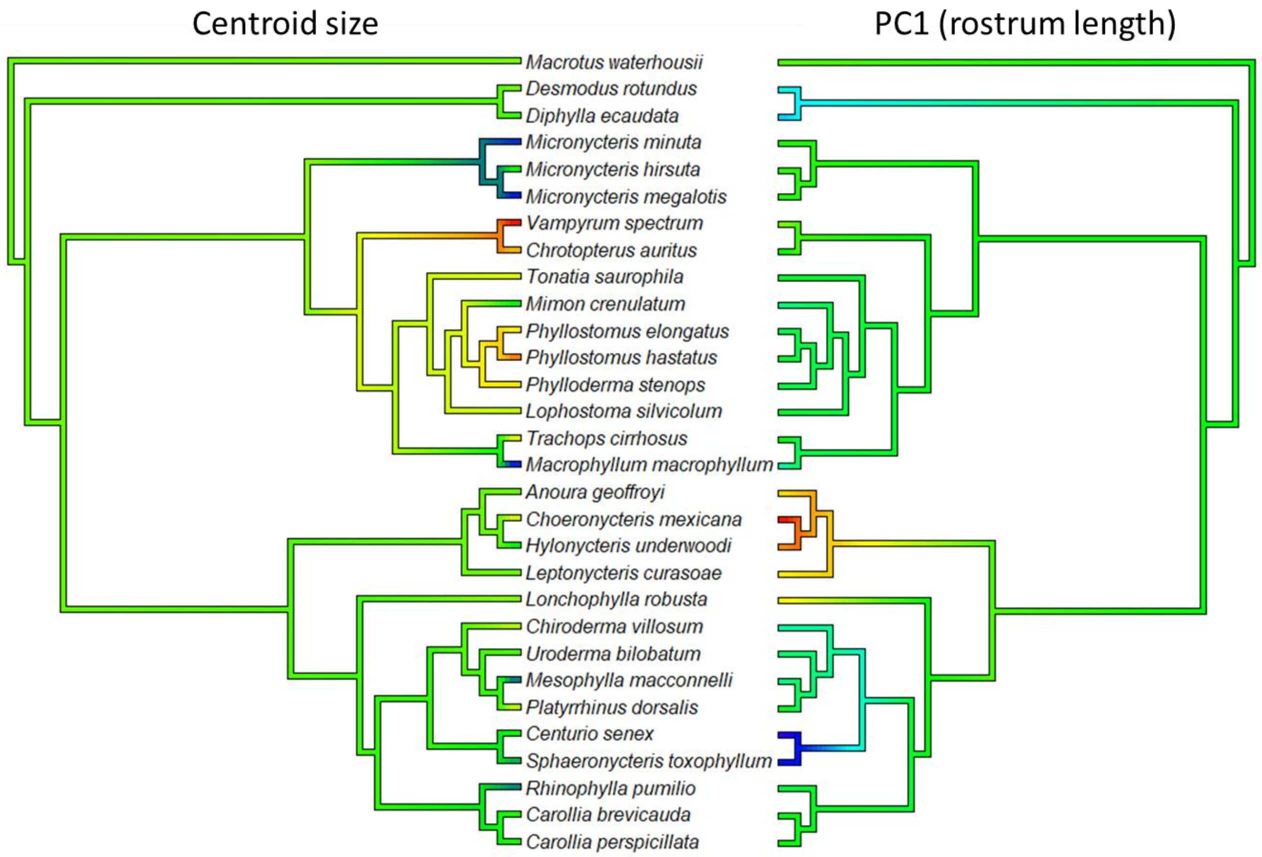

**Figure S15: Phylogeny of the Phyllostomidae, coloured on the left by cranial centroid size from smallest (blue) to largest (red), and on the right by PC1 representing relative rostrum length from shortest (blue) to longest (red).**

**Molossidae**

The Molossidae dataset comprises 13 species with a centroid size scaling range of 2.03. There was no significant allometry detected (Fig. S16A). The largest and smallest species are morphologically average within the shape space (Fig. S16B), but the first principal component does represent relative face length (Fig. S16C). There is no significant allometry after phylogenetic correction, nor is there a significant phylogenetic signal of centroid size. In fact, this is the only group tested that shows no significance in any tests carried out. Much of the craniodental variation within this family has been attributed to dietary variation, with species feeding on tougher insects tending to be larger, with more robust heads (Villalobos-Chaves & Santana, 2022). No evidence of allometry suggests overall isometry in cranial growth, which would suggest that, on average, the rate of insect toughness included in diets increases at a similar rate to bite force with size (i.e., an ABC of 1; see Fig 8 of main manuscript).

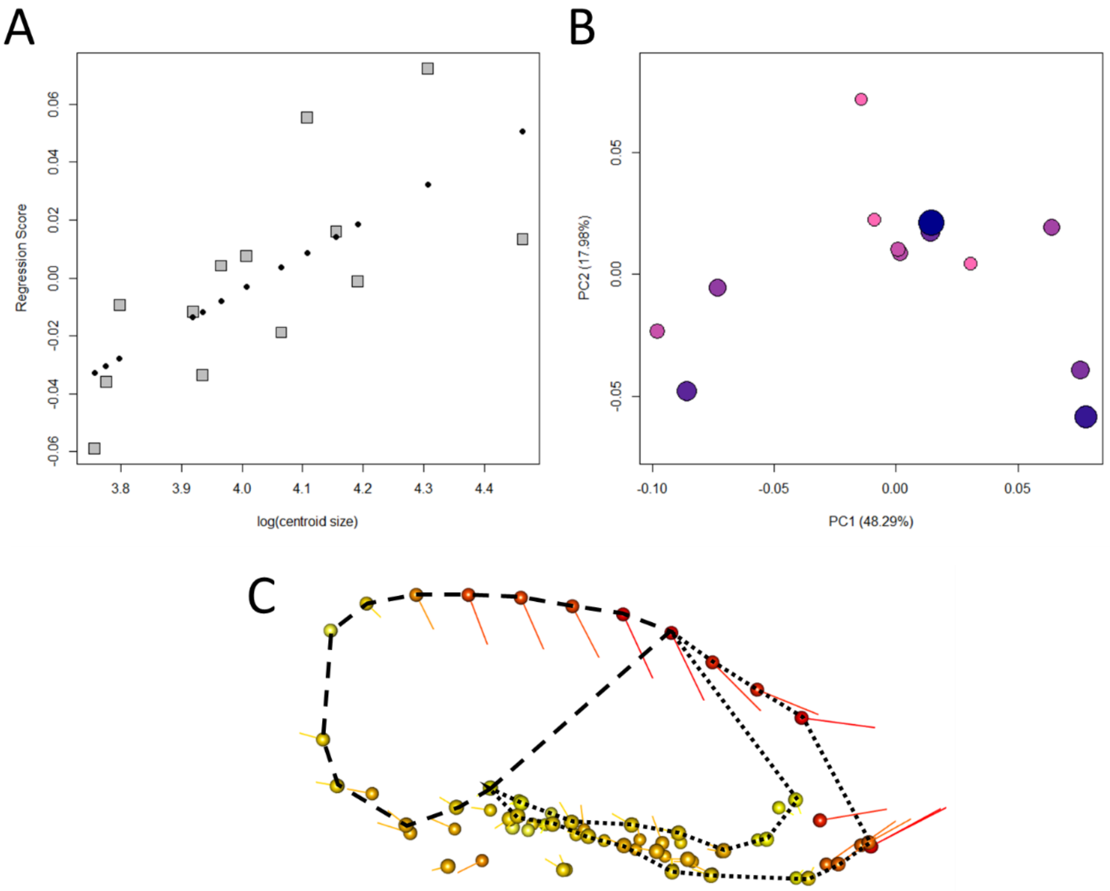

**Figure S16: (A) Shape regression score over log(centroid size) (black dots = PredLine). (B) plot of the first two principal components (orb size represents cranial centroid size), (C) shape variation defined by PC1 (orbs represent the minimum extreme, line tips are the maximum). Dashed lines = braincase, dotted lines = face.**

### Vespertilionidae

The Vespertilionidae comprises 29 species with a centroid size scaling index of 1.980. Allometry was significant ($R^2$ = 0.084, p=0.041) (Fig. S17A), with larger species predicted to have relatively smaller braincases and shorter, deeper facial skeleton, inconsistent with predictions of facial gracility (Fig. S17B). No principal components were correlated with size (Fig. S17C), despite the first PC being associated with face length (Fig. S17D). There is no significant allometry detected after phylogenetic correction, but there is a significant phylogenetic signal for centroid size (p=0.001).

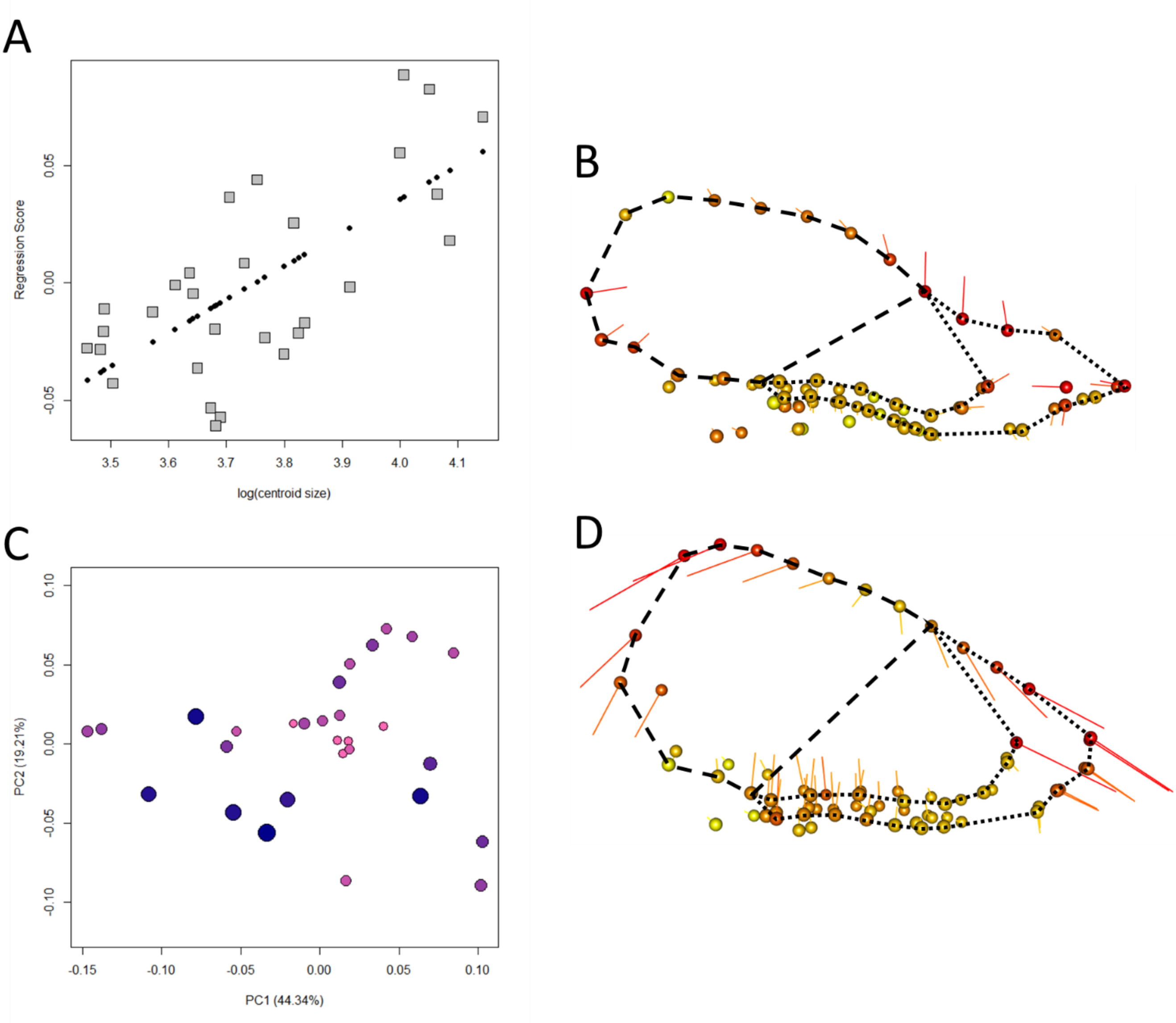

**Figure S17: (A) Shape regression score over log(centroid size) indicates allometric relationship (black dots = PredLine). (B) The predicted shape of the PredLine for the ordinary least squares allometry test, (C) plot of the first two principal components (orb size represents cranial centroid size), (D) shape variation defined by PC1 (orbs represent the minimum extreme, line tips are the maximum). Dashed lines = braincase, dotted lines = face.**

**Felidae**

The Carnivora dataset comes from Meloro & Tagmanini (2021). The Felidae data includes 32 species with a centroid size scaling index of 3.962. The landmarks consist of 2D ventral landmarks. Allometry is significant ($R^2$ = 0.367, p=0.001) (Fig. S18A), with larger species predicted to have relatively smaller basicranial, larger temporomandibular joints (TMJs) and extended premaxillae (Fig. S18B). The first principal component (Fig. S18C) is significantly correlated with centroid size (p<0.001) and is defined by basicranial size, size of the jaw joints, and premaxilla projection (Fig. S18D). There is no allometry following phylogenetic correction but there is a significant phylogenetic signal for centroid size (p=0.001). The Felidae clearly exhibit the gracilisation pattern, with similar distributions of cranial size and face length across the phylogeny (Fig. S19).

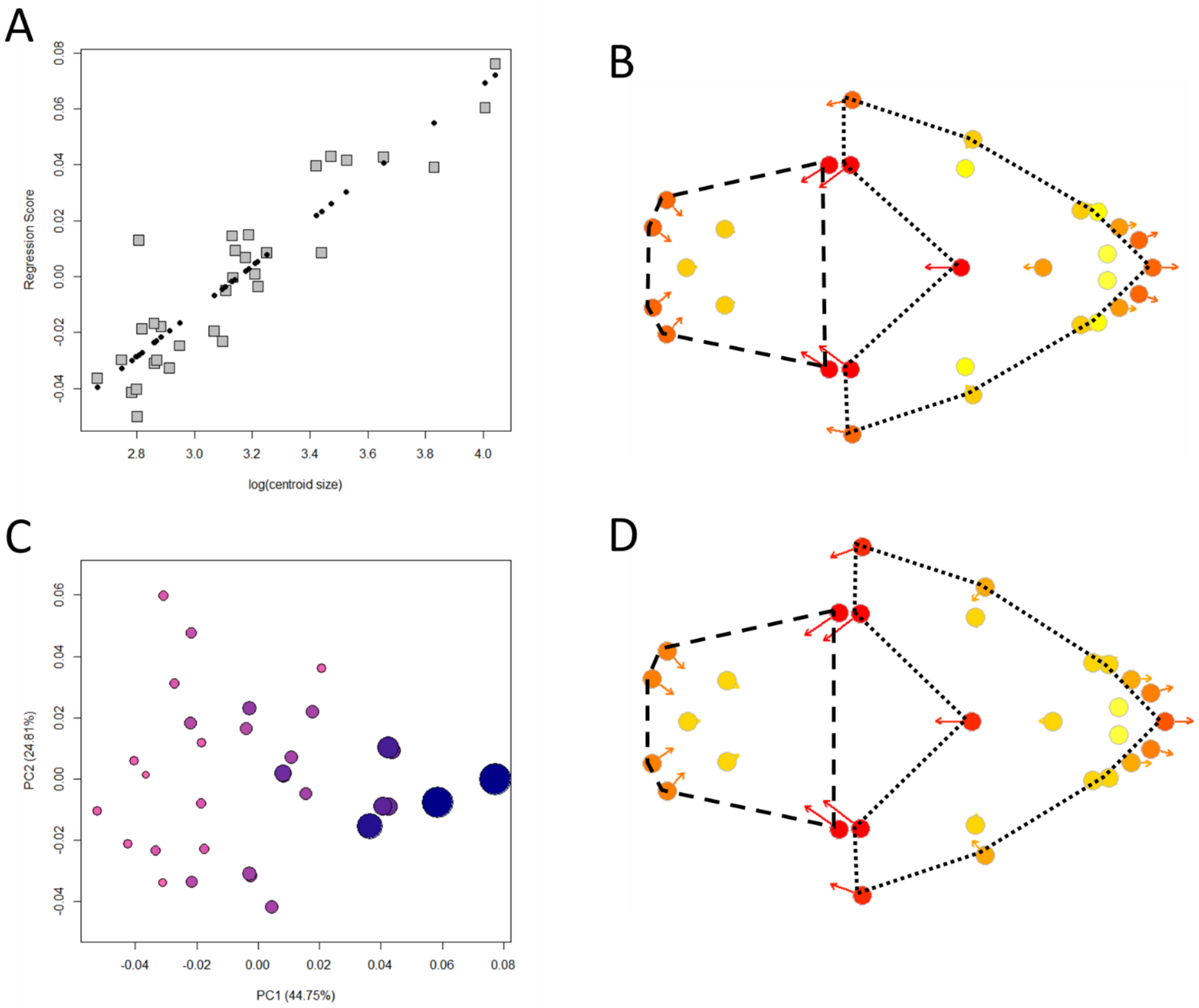

**Figure S18: (A) Shape regression score over log(centroid size) indicates allometric relationship (black dots = PredLine). (B) The predicted shape of the PredLine for the initial allometry test (orbs represent smaller sizes). (C) Plot of the first two principal components (orbs represent centroid sizes) (D) Shape variation defined by PC1 (orbs represent the minimum extreme). Dashed lines = braincase, dotted lines = face.**

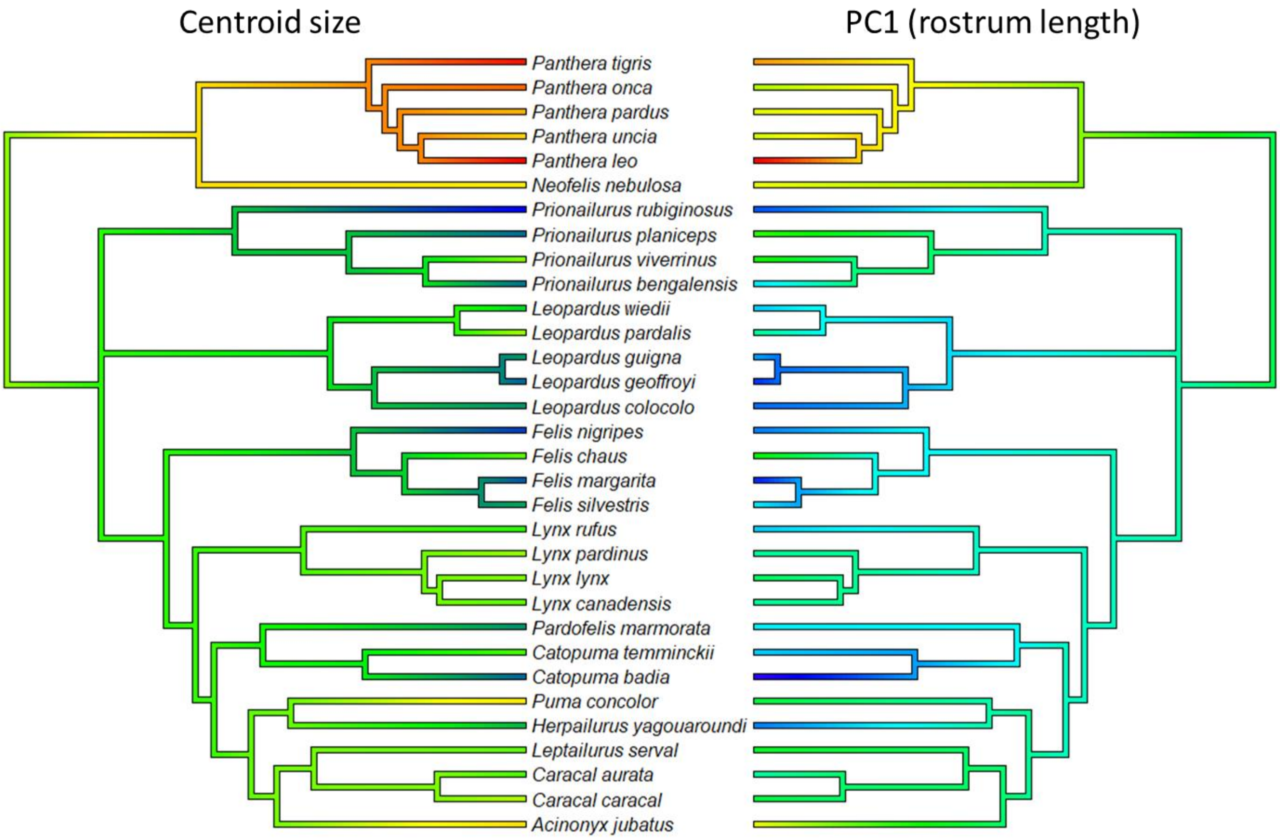

**Figure S19: Phylogeny of the Felidae, coloured on the left by cranial centroid size from smallest (blue) to largest (red), and on the right by PC1 representing relative rostrum length from shortest (blue) to longest (red).**

### Viverridae

The Viverridae data includes 15 species with a centroid size scaling index of 1.745. Allometry is significant ($R^2$ = 0.216, p=0.018) (Fig. S20A), represented by smaller basicrania and projected premaxillae in larger species (Fig. S20B). PC1 is weakly, yet significantly correlated with size (p=0.038) (Fig. S20C), representing the position of the dentition and temporomandibular joint size, but not facial gracility (Fig S20D). The correlation is largely attributable to the cluster of small species (*Genetta* spp.) representing the minimum range of PC1. PC2 is associated with relative face length but is not correlated with size. Allometry is no longer significant after phylogenetic correction, but there is a significant phylogenetic signal for centroid size (p=0.011).

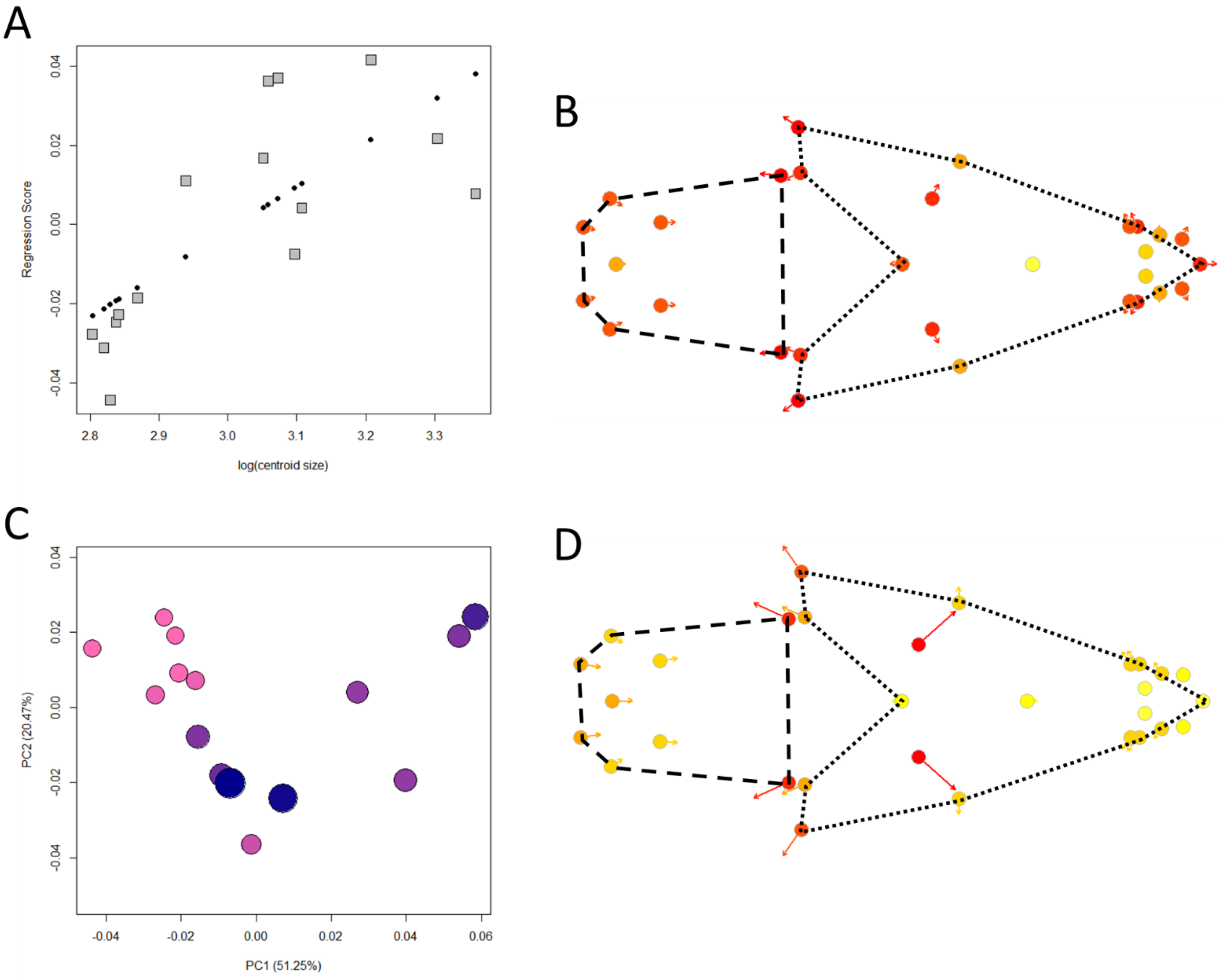

**Figure S20: (A) Shape regression score over log(centroid size) indicates allometric relationship (black dots = PredLine). (B) The predicted shape of the PredLine for the initial allometry test (orbs represent smaller sizes). (C) Plot of the first two principal components (orbs represent centroid sizes) (D) Shape variation defined by PC1 (orbs represent the minimum extreme). Dashed lines = braincase, dotted lines = face.**

**Herpestidae**

The Herpestidae data includes 17 species with a centroid size scaling index of 2.348. Allometry is not quite significant ($R^2$ = 0.141, p=0.052) (Fig. S21A), however, we are willing to concede the possibility of marginal statistical error here, given the significant allometry found in prior research (Cardini & Polly, 2013). The trend is for a decreased braincase size and a narrower cranium with increased size (Fig. S21B). PC1 represents cranial width but is not correlated with size (Fig. S21C). PC2 is correlated with size (p=0.002) and represents a smaller basicranium and a longer, narrower face with increased size (Fig S21D). Allometry is significant after phylogenetic correction ($R^2$=0.289, p=0.028). Larger species are expected to have narrower crania, but no shift in premaxillae projection (Fig. S21E) supporting the pattern of hyperallometric gracilisation. There is no significant phylogenetic signal of centroid size. While PC2 is correlated with size, the largest species, *Herpestes naso* and *Bdeogale nigripes*, are located at the mid-range of PC2 (Fig. S22).

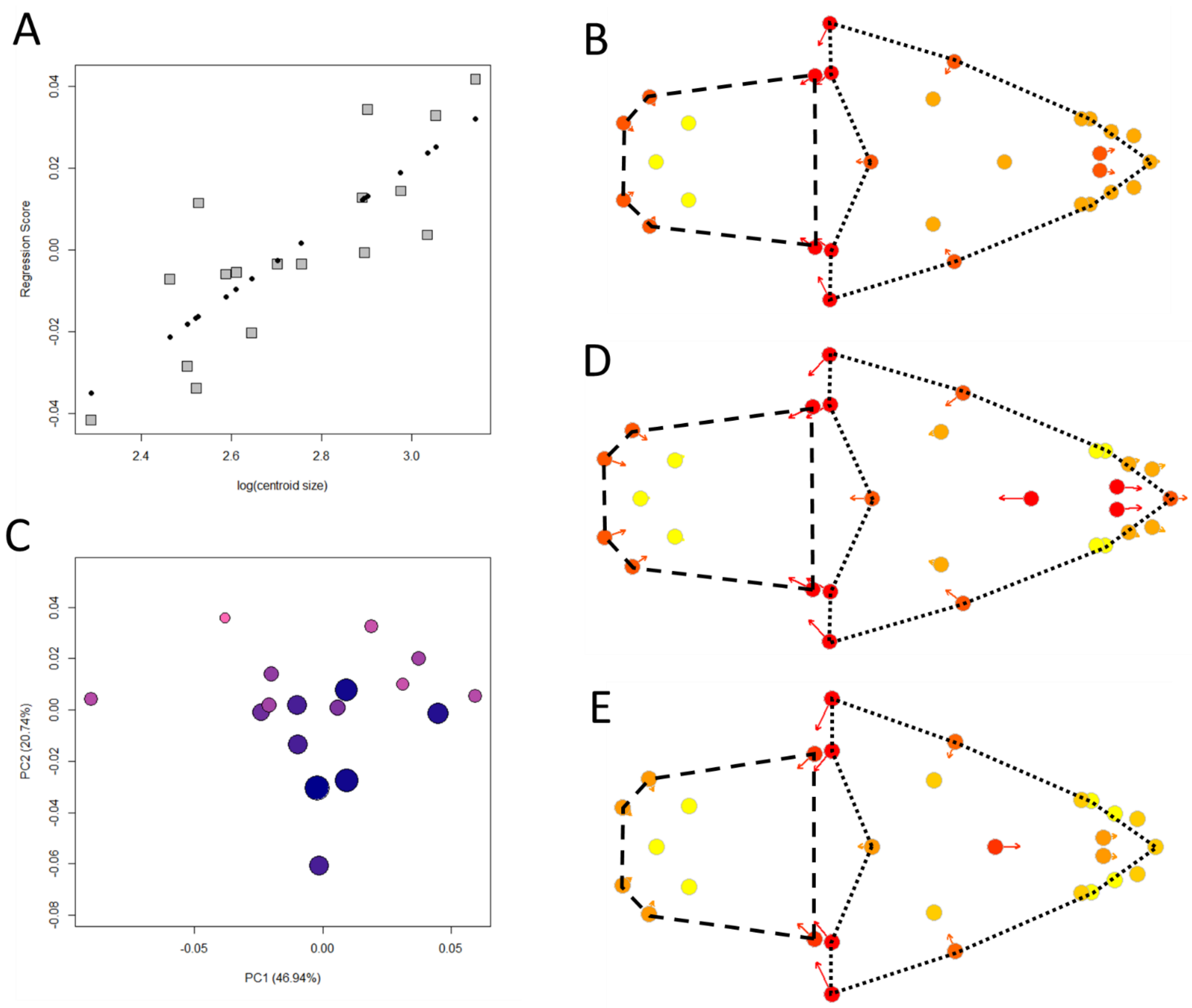

**Figure S21: (A) allometry plot of cranial shape score regressed on log(cranial centroid size), (B) allometric predictions of cranial shape for ordinary least squares, (C) Plot of first and second principal components, (D) shape differences across the second principal component, (E) allometric predictions of cranial shape for phylogenetic generalised least squares. Dashed lines = braincase, dotted lines = face.**

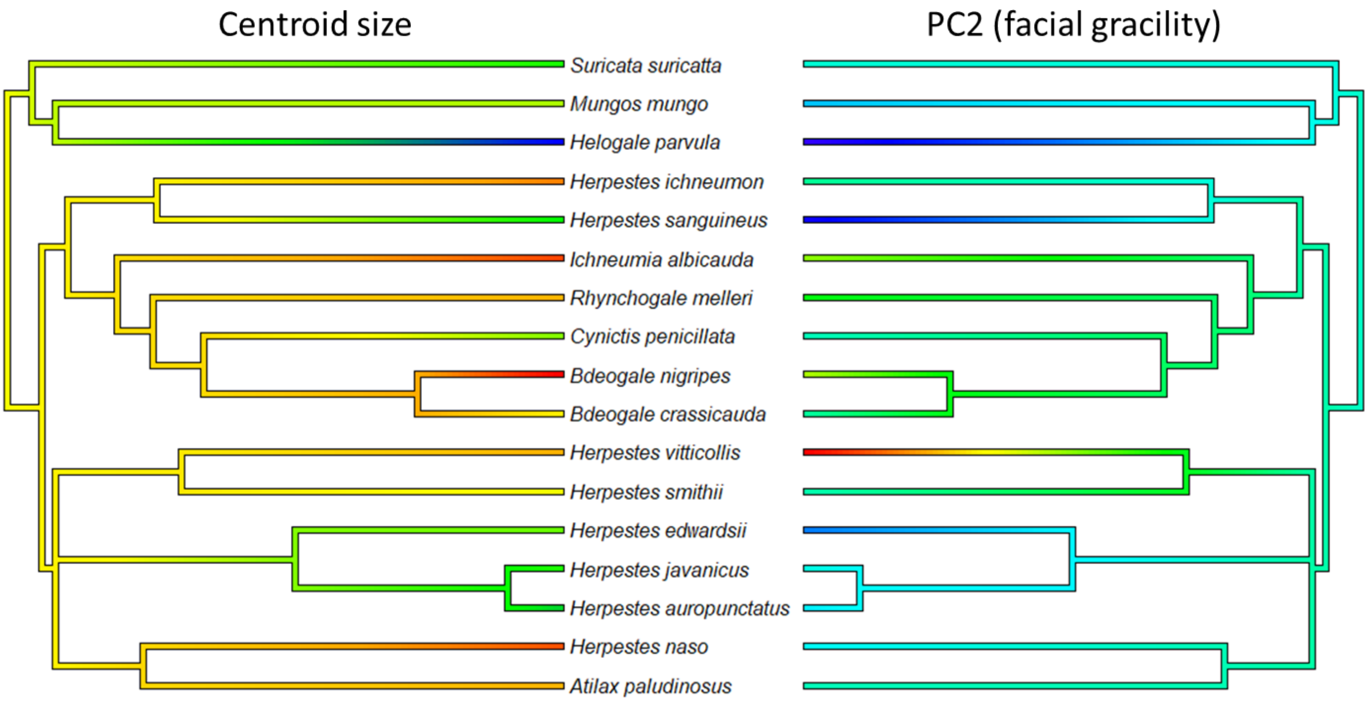

**Figure S22: Phylogeny of the Herpestidae, coloured on the left by cranial centroid size from smallest (blue) to largest (red), and on the right by PC2 representing relative facial gracility from robust (blue) to gracile (red).**

### Canidae

The Canidae data includes 30 species with a centroid size scaling index of 2.707. Allometry is significant ($R^2$ = 0.108, p=0.017) (Fig. S23A), but is largely confined to TMJ size, with only very slight premaxillae extension (Fig. S23B). This supports the relative isometry in facial size previously found across canids (Wayne, 1986; and see Fig. 3 of the main manuscript showing comparable facial length in the smallest species (*Vulpes zerda*) and the largest (*Canis lupus*), despite allometric differences in relative braincase size and orbit size). Neither PC1 nor PC2 are correlated with size. PC3 is correlated with size and is largely defined by the size of the temporomandibular joint, which explains our allometric predictions (Curth et al., 2017). The first principal component is defined by facial gracilisation, via relative cranial width and anteroposterior positioning of the zygomatic arch (Fig. S23D). The minimum PC1 extreme for canids is occupied by the three particularly robust crania of the South American bush dog (*Speothos venaticus*), the African wild dog (*Lycaon pictus*), and the dhole (*Cuon alpinus*); three of the four recognised large prey specialists (Van Valkenburgh & Koepfli, 1993). These species are medium-sized canids with their relatively shorter faces determined by a widening and anterior projection of the zygomatic arches (Van Valkenburgh 1991; Van Valkenburgh and Koepfli, 1993). There is no allometric signal following phylogenetic correction but there is a significant phylogenetic signal for centroid size (p=0.001). Figure S24 shows little similarity between cranial size and facial gracilisation across the entire clade, however, there is some indication of a correlation within the *Vulpes* and *Canis* clades (see main manuscript).

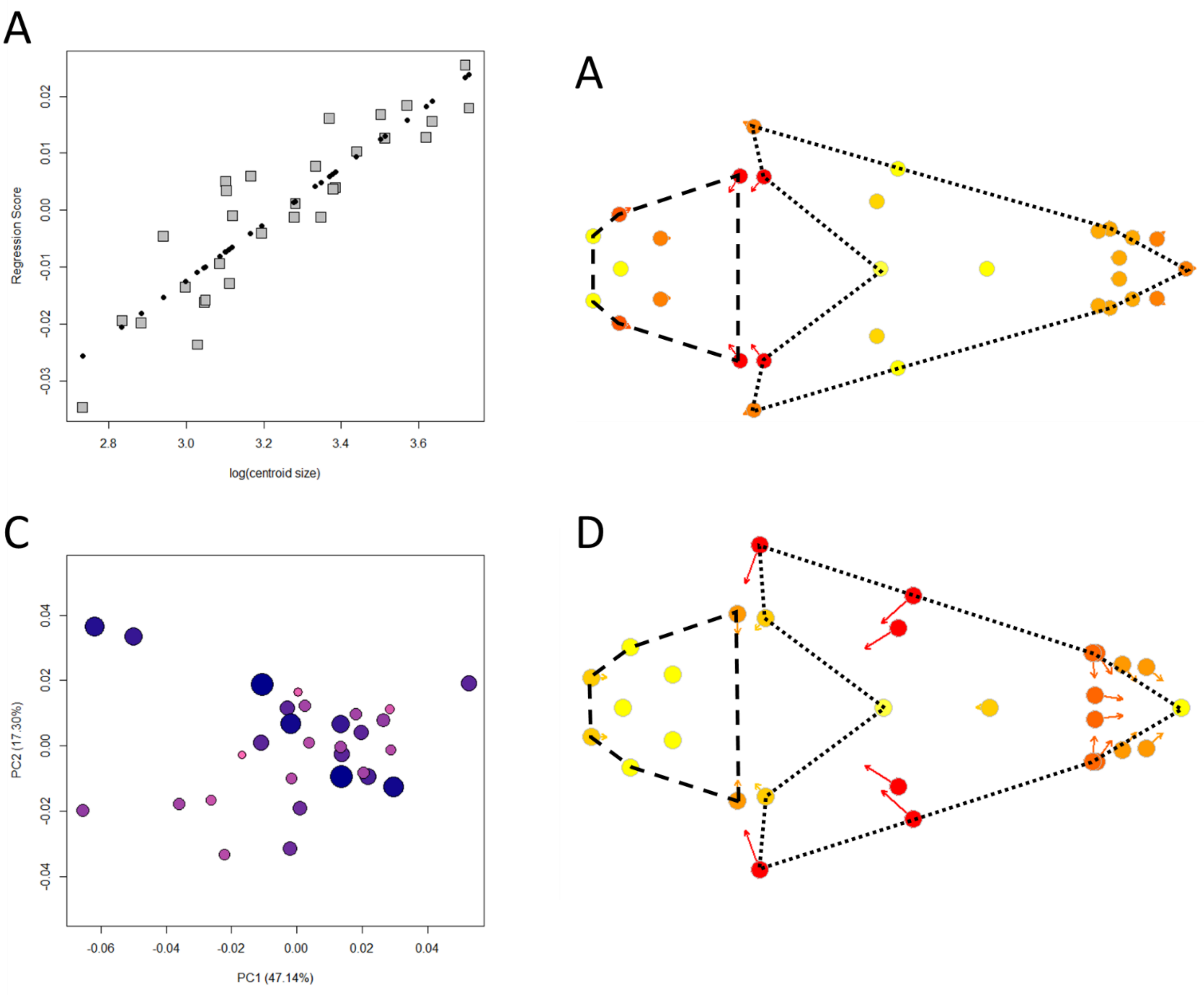

**Figure S23: (A) Shape regression score over log(centroid size) indicates allometric relationship (black dots = PredLine). (B) The predicted shape of the PredLine for the initial allometry test (orbs represent smaller sizes). (C) Plot of the first two principal components (orbs represent centroid sizes) (D) Shape**

**variation defined by PC1 (orbs represent the minimum extreme). Dashed lines = braincase, dotted lines = face.**

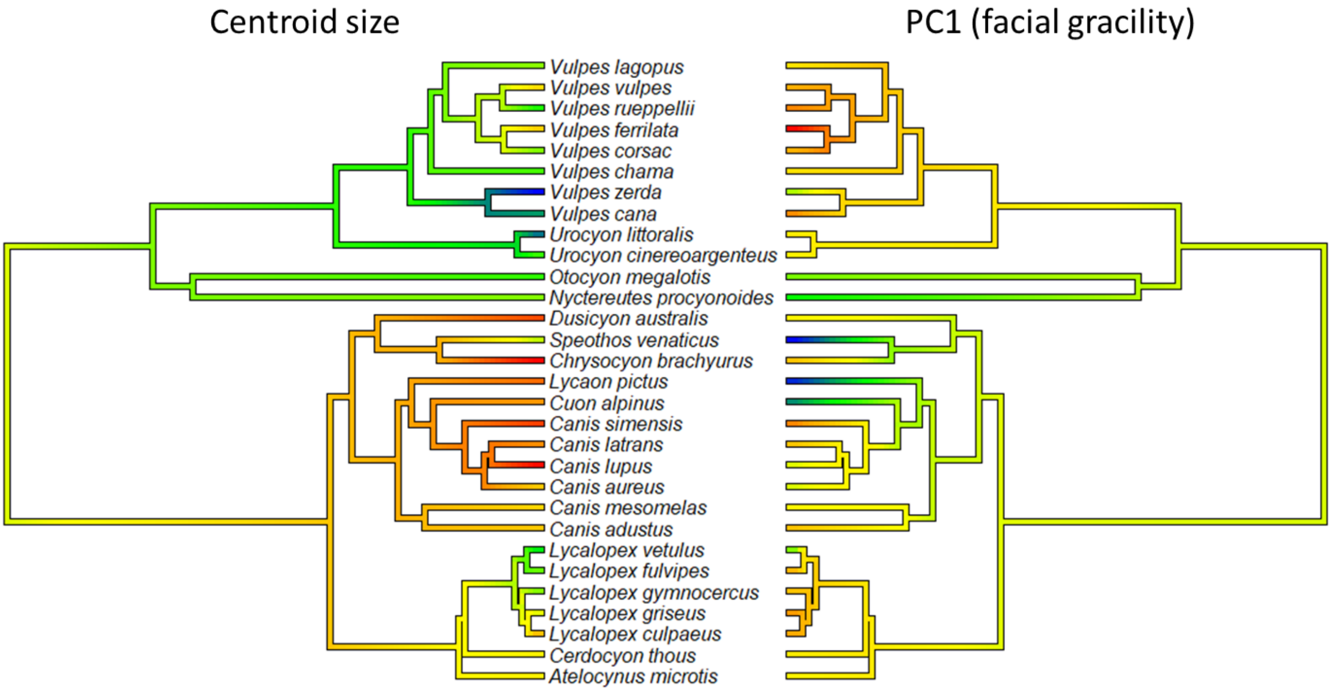

**Figure S24: Phylogeny of the Canidae, coloured on the left by cranial centroid size from smallest (blue) to largest (red), and on the right by PC1 representing relative facial gracility from robust (red) to gracile (blue).**

**Mustelidae**

The Mustelidae data includes 34 species with a centroid size scaling index of 4.387. Allometry is significant ($R^2$ = 0.249, p=0.001) (Fig. S25A), with larger species predicted to have relatively smaller basicrania, and slightly extended premaxillae (Fig. S25B). The first principal component (Fig. S25C) is significantly correlated with centroid size (p<0.001) and is defined by basicranial size and premaxillae projection (Fig. S25D), supporting the pattern. However, PC2 is also correlated with size (P=0.003) and is almost entirely explained by cranial width (Fig. S25E). The Mustelidae therefore appear to demonstrate two uncorrelated allometric trajectories, PC1 in support of hyperallometric gracilisation, PC2 being in opposition. Interestingly, the species representing extremes of both trajectories identify opposing extremes of rostrum elongation: the long-faced hog badger (*Arctonyx collaris*) that feeds primarily on soft forest fruits and invertebrates and the short-faced sea otter (*Enhydra lutris*) which often feeds on hard shellfish, both of which are similar in size and were presented by Radinsky (1981) as a showcase of mustelid cranial diversity. There is no allometric signal following phylogenetic correction but there is a significant phylogenetic signal for centroid size (p=0.001).

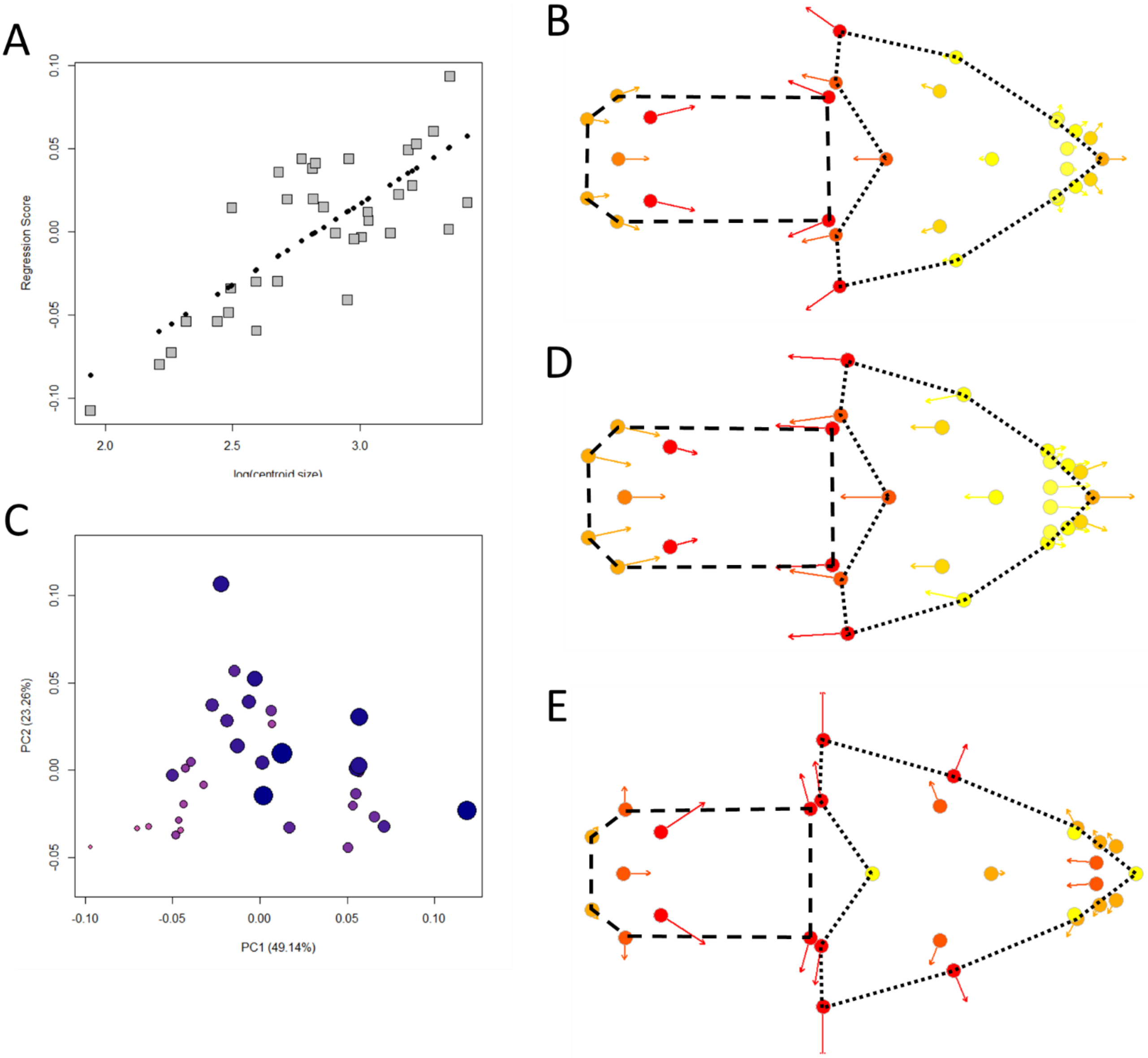

**Figure S25: (A) Shape regression score over log(centroid size) indicates allometric relationship (black dots = PredLine). (B) The predicted shape of the PredLine for the initial allometry test (orbs represent smaller sizes). (C) Plot of the first two principal components (orbs represent centroid sizes), (D) Shape variation defined by PC1 (orbs represent the minimum extreme) (E) Shape variation defined by PC2 (orbs represent the minimum extreme). Dashed lines = braincase, dotted lines = face.**

**Phocidae**

The Phocidae data includes 15 species with a centroid size scaling range of 2.062. Allometry is significant ($R^2$ = 0.209, p=0.003) (Fig. S26A), represented by smaller a basicranium and longer zygomatic arches in larger species (Fig. S26B). This is likely to result in more robust facial proportions. PC1 is significantly correlated with size (p=0.002) (Fig. S26C), representing basicranium size and zygomatic arch elongation (Fig S26D). PC2 describes facial gracilisation via narrowing of the zygomatic arches and projection of the premaxillae, but is not correlated with size. Allometry is significant after phylogenetic correction ($R^2$=0.366, p=0.023). Larger species for this family differ from all others, with what appears to be isometric growth of the facial skeleton in both length and width, alongside smaller braincase size (Fig. S26E). This does not agree with predictions of hyperallometric gracilisation. There is also a marginal phylogenetic signal of centroid size (p=0.037).

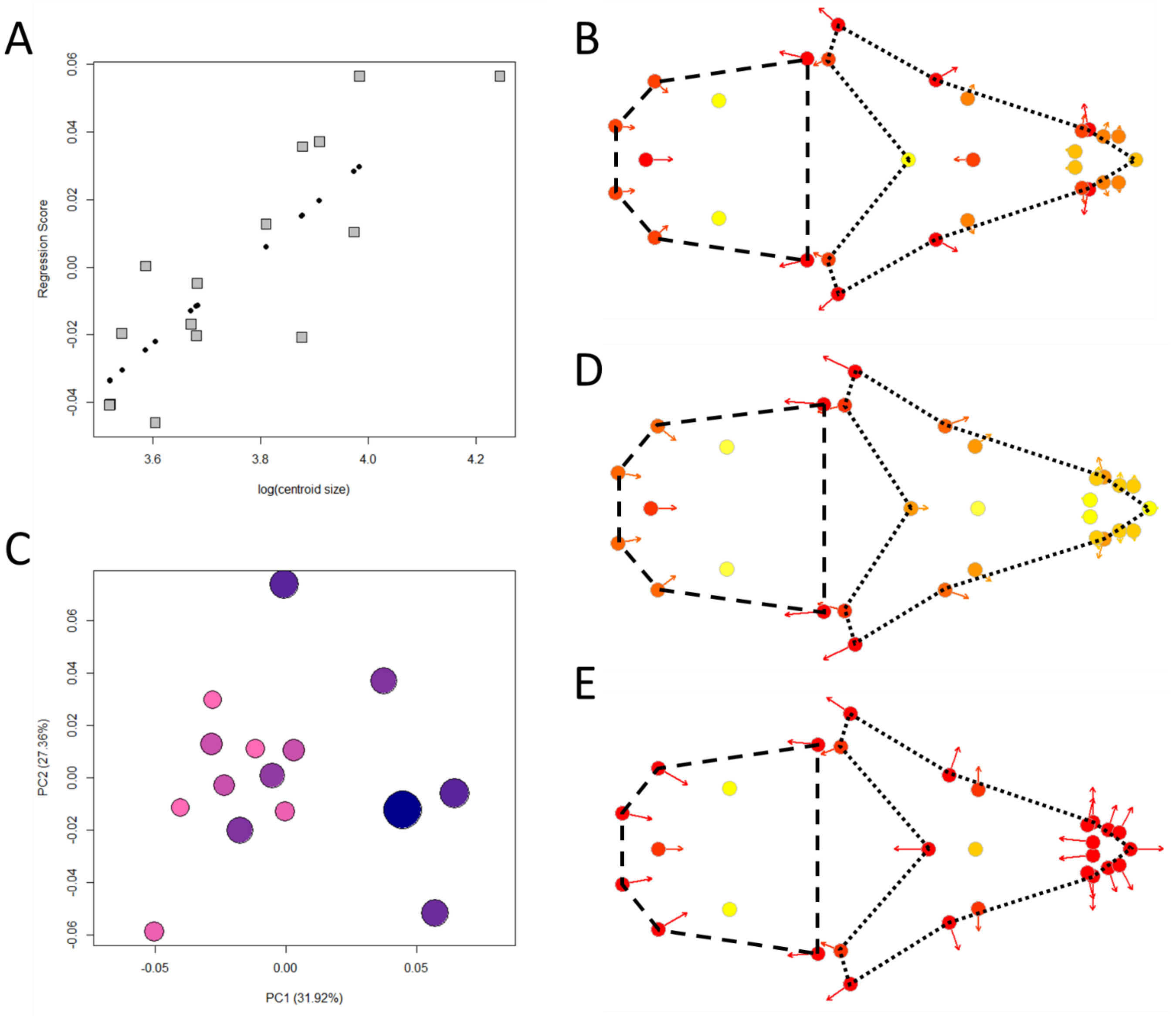

**Figure S26: (A) allometry plot of cranial shape score regressed on log(cranial centroid size), (B) allometric predictions of cranial shape for ordinary least squares, (C) Plot of first and second principal components, (D) shape differences across the first principal component, (E) allometric predictions of cranial shape for phylogenetic generalised least squares. Dashed lines = braincase, dotted lines = face.**

### Otariidae

The Otariidae data includes 12 species with a centroid size scaling index of 1.607. Allometry is significant ($R^2$ = 0.343, p=0.001) (Fig. S27A), represented largely by the position of the rear palate, with some minor predictions of reduced basicranial size (Fig. S27B). PC1 is significantly correlated with size (p<0.001) (Fig. S27C), almost entirely representing the position of the rear palate (Fig S27D). PC2 is not correlated with size. Allometry is not significant after phylogenetic correction, nor is there a significant phylogenetic signal of centroid size.

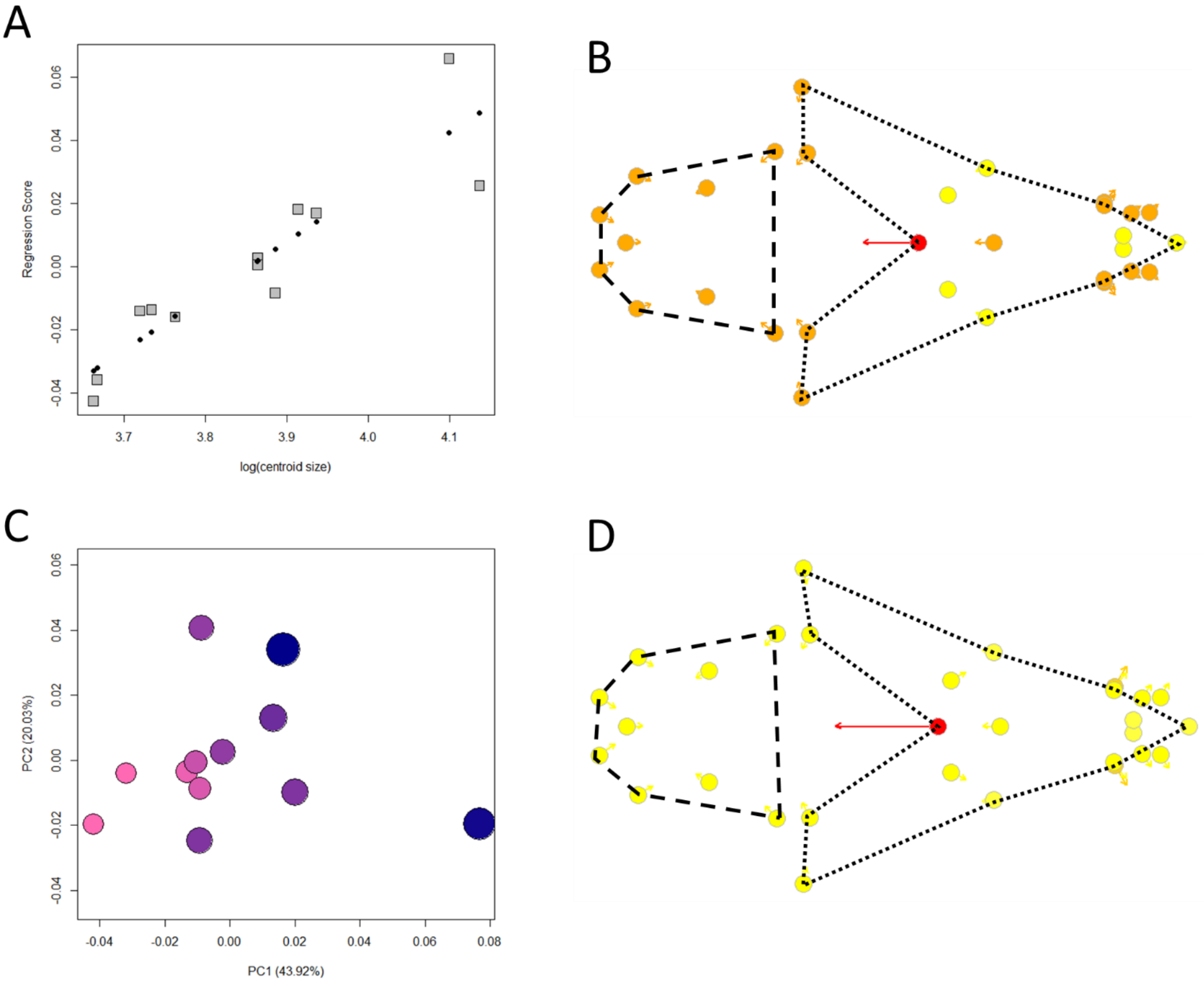

**Figure S27: (A) Shape regression score over log(centroid size) indicates allometric relationship (black dots = PredLine). (B) The predicted shape of the PredLine for the initial allometry test (orbs represent smaller sizes). (C) Plot of the first two principal components (orbs represent centroid sizes) (D) Shape variation defined by PC1 (orbs represent the minimum extreme). Dashed lines = braincase, dotted lines = face.**

**Bovidae**

This data comes from Bibi & Tyler (2022). We tested allometry on 88 species. The centroid size scaling index was 6.664. Allometry is significant ($R^2$ = 0.188, p=0.001) (Fig. S28A), with larger species predicted to have relatively smaller orbits and braincases. There is a slight tendency for larger species to exhibit elongated premaxillae (Fig. S28B). The first principal component (Fig. S28C) is significantly correlated with centroid size (p<0.001) and is defined by orbit size, braincase size, and a stronger association with premaxillae projection (Fig. S28D). PC3 is also correlated with size, and largely represents the positioning of the antlers. Allometry remains significant after phylogenetic correction ($R^2$=0.066, p=0.001). Larger species are expected to exhibit smaller braincases, smaller orbits, and elongated premaxillae (Fig. S28E) supporting the predicted pattern of hyperallometric gracilisation on all fronts, with a significant phylogenetic signal of centroid size (p=0.001). A comparison of cranial size and face length across phylogenies is shown in Fig. S29. While absolute values might differ, the trends for both features are similar.

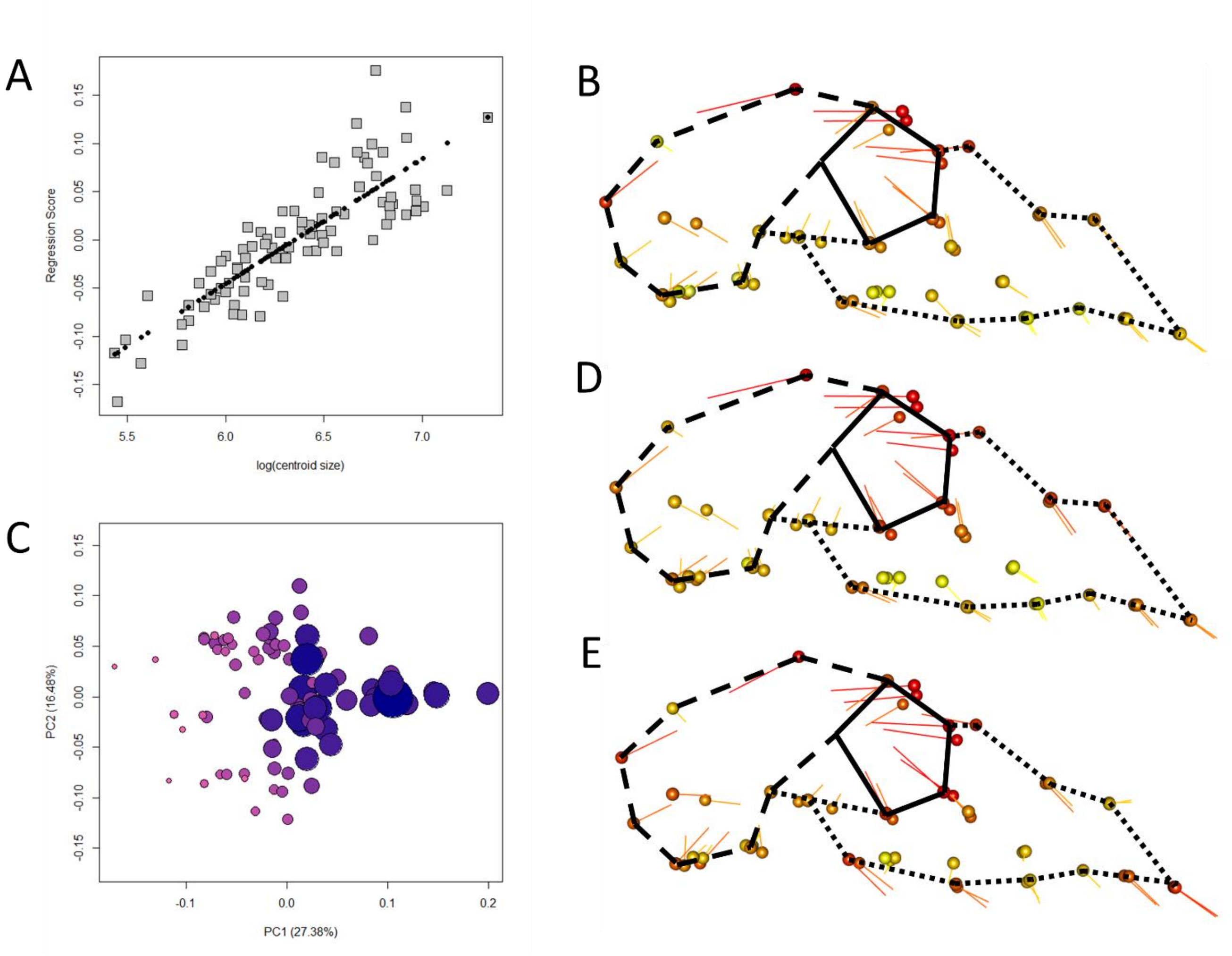

**Figure S28: Bovidae plots (A) allometry plot of cranial shape score regressed on log(cranial centroid size), (B) allometric predictions of cranial shape for ordinary least squares, (C) Plot of first and second principal components, (D) shape differences across the first principal component, (E) allometric predictions of cranial shape for phylogenetic generalised least squares. Dashed lines = braincase, dotted lines = face.**

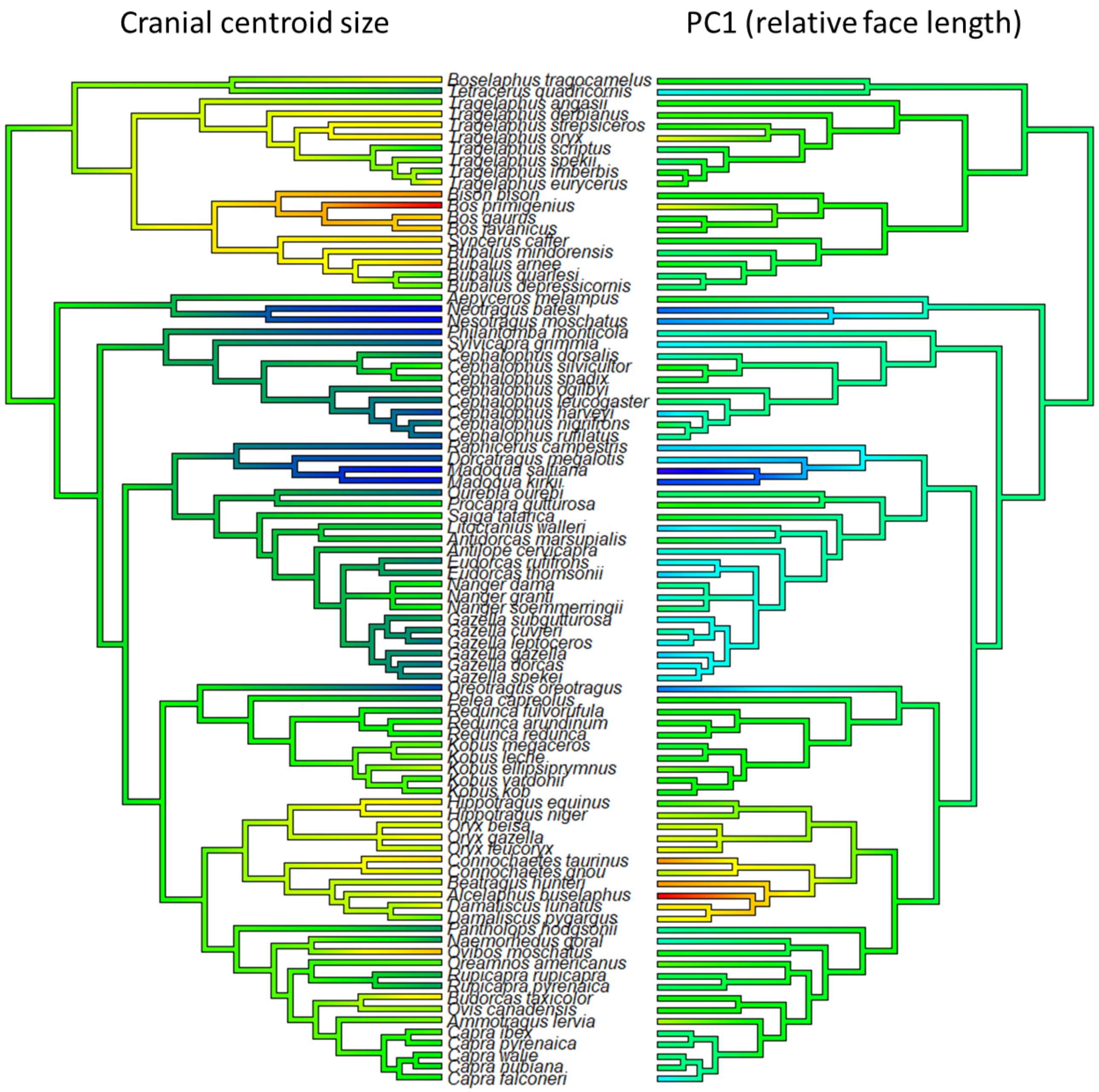

**Figure S29: Phylogeny of the Bovidae, coloured on the left by cranial centroid size from smallest (blue) to largest (red), and on the right by PC1 representing relative rostrum length from shortest (blue) to longest (red).**

**Delphinidae**

The data for the Cetacea comes from Coombs et al. (2020) and includes extinct taxa. The Delphinidae data includes 38 species with a centroid size scaling index of 3.824. There is a significant allometric signal ($R^2$=0.084, p=0.017) (Fig. S30A). The predicted shape changes suggest larger species have a smaller braincase, but no shift in rostrum length (Fig. S30B). PC1 is not correlated with size (p=0.715) but is largely explained by the length of the rostrum (Fig. S30D). But PC2 is correlated with size (p<0.001) and represents relative braincase size (Fig S30E). Braincase size and rostrum length are therefore uncorrelated across this group, likely associated with the relatively more robust crania of the largest species, *Orcinus orca*, and members of the mostly short-faced subfamily, Globicephalinae (*Steno bredanensis* and company), that span much of the size range (Fig. S31). While *O. orca* is known to take much larger prey, such as seals, the Globicephalinae are specialised suction feeders of smaller, soft-bodied prey. They therefore present an interesting case where a similar, stout-faced morphology occurs alongside entirely opposing biting mechanics. Allometry is not significant after phylogenetic correction, but there is a significant phylogenetic signal of cranial centroid size (p=0.001).

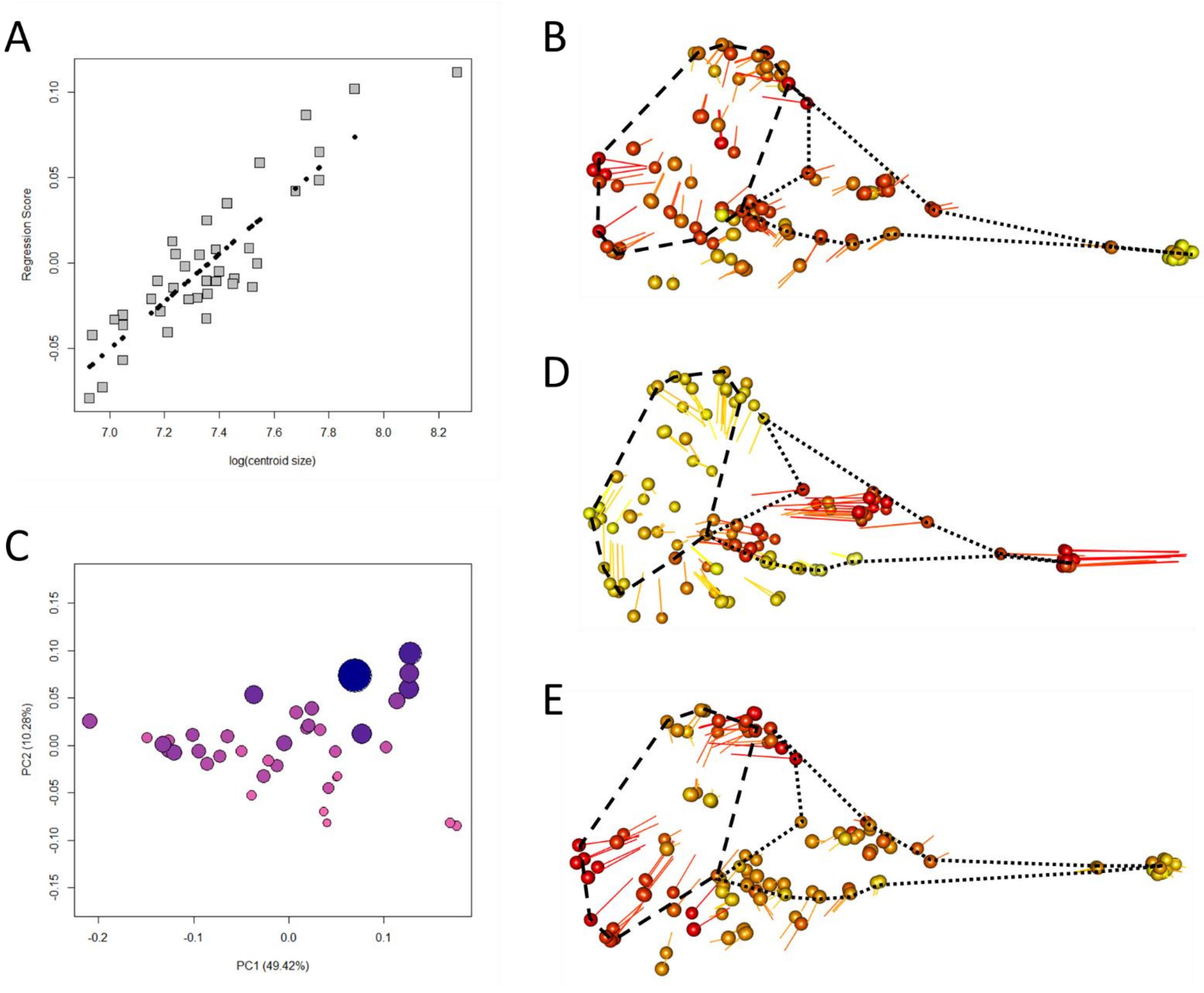

**Figure S30: (A) allometry plot of cranial shape score regressed on log(cranial centroid size), (B) allometric predictions of cranial shape for ordinary least squares, (C) Plot of first and second principal components, (D) shape differences across the first principal component, (E) shape differences across the second principal component. Dashed lines = braincase, dotted lines = face.**

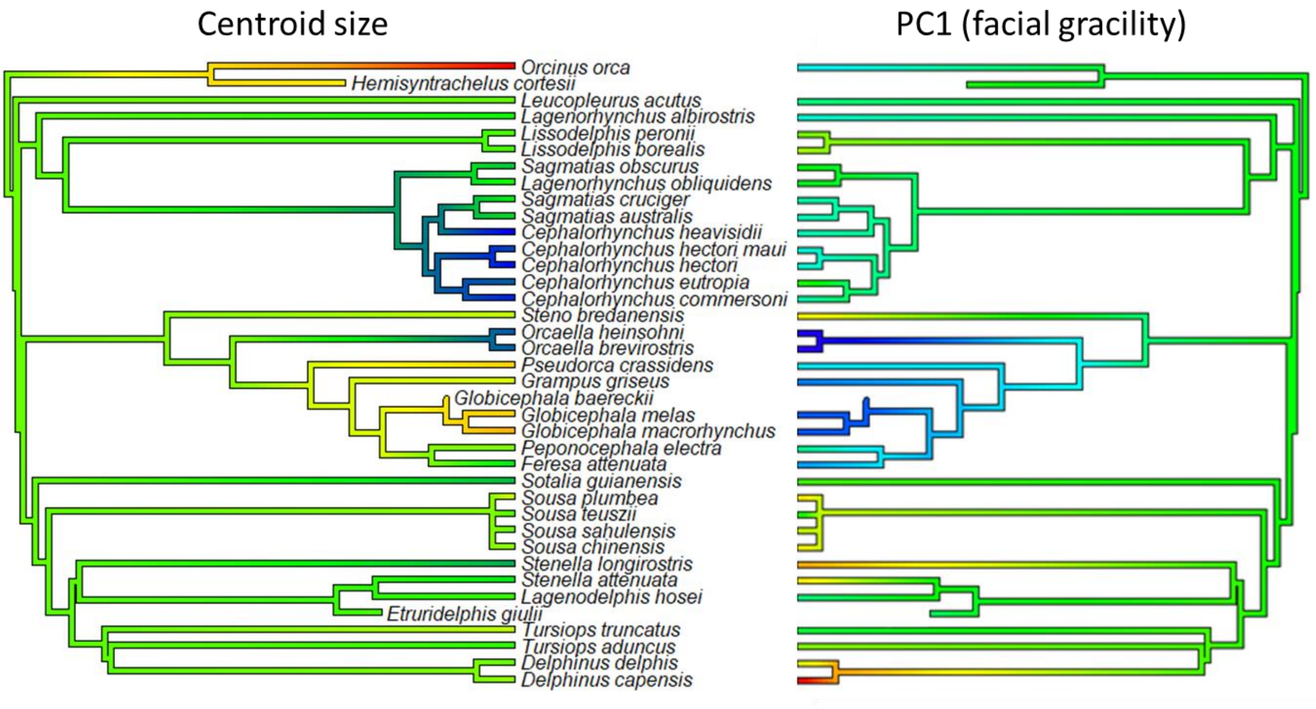

**Figure S31: Phylogeny of the Delphinidae, coloured on the left by cranial centroid size from smallest (blue) to largest (red), and on the right by PC1 representing relative rostrum length from shortest (blue) to longest (red).**

### Phocoenidae

The Phocoenidae data includes 11 species with a size scaling index of 2.162. There is a significant allometric signal ($R^2$ = 0.418, p=0.002) (Fig. S32A). The predicted shape changes suggest larger species have a smaller braincase and longer rostrum (Fig. S32B). PC1 is correlated with size (p<0.001) and includes braincase and rostrum length (Fig. S32C). PC2 and PC3 are not correlated with size. Allometry remains significant after phylogenetic correction ($R^2$=0.172, p=0.009). Larger species are predicted to have a smaller braincase and longer rostrum (Fig. S32E), supporting hyperallometric gracilisation. There is no significant phylogenetic signal for centroid size. Fig. S33 shows high congruence between centroid size and facial gracility across the phylogeny. Of relevance to this review, snout morphology for the Phocoenidae and Delphinidae (together Delphinoidea) have been shown to correlate strongly with diet across latitudinal gradients (McCurry et al., 2023).

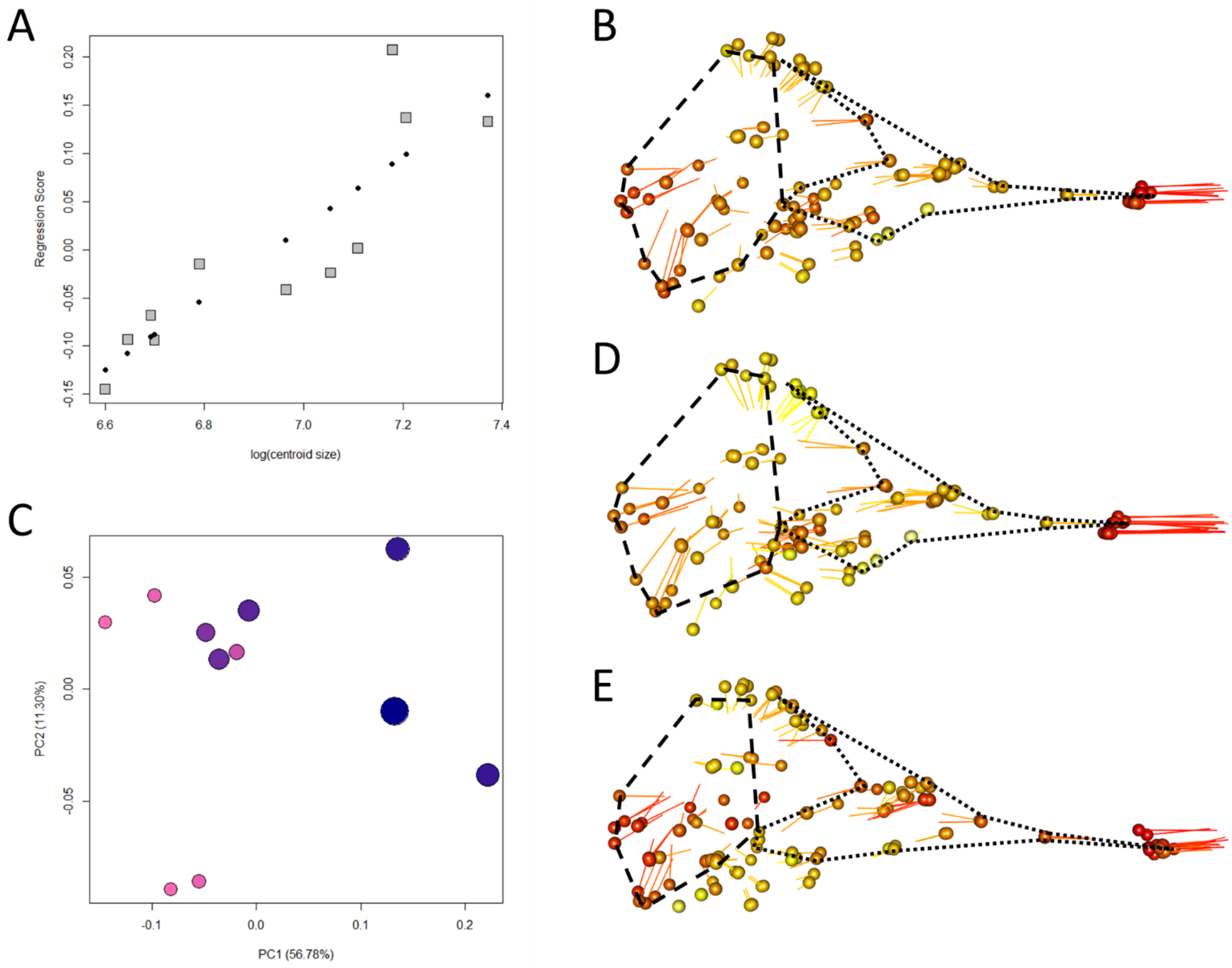

**Figure S32: (A) allometry plot of cranial shape score regressed on log(cranial centroid size), (B) allometric predictions of cranial shape for ordinary least squares, (C) plot of first and second principal components, (D) shape differences across the first principal component, (E) allometric predictions of cranial shape for phylogenetic generalised least squares. Dashed lines = braincase, dotted lines = face.**

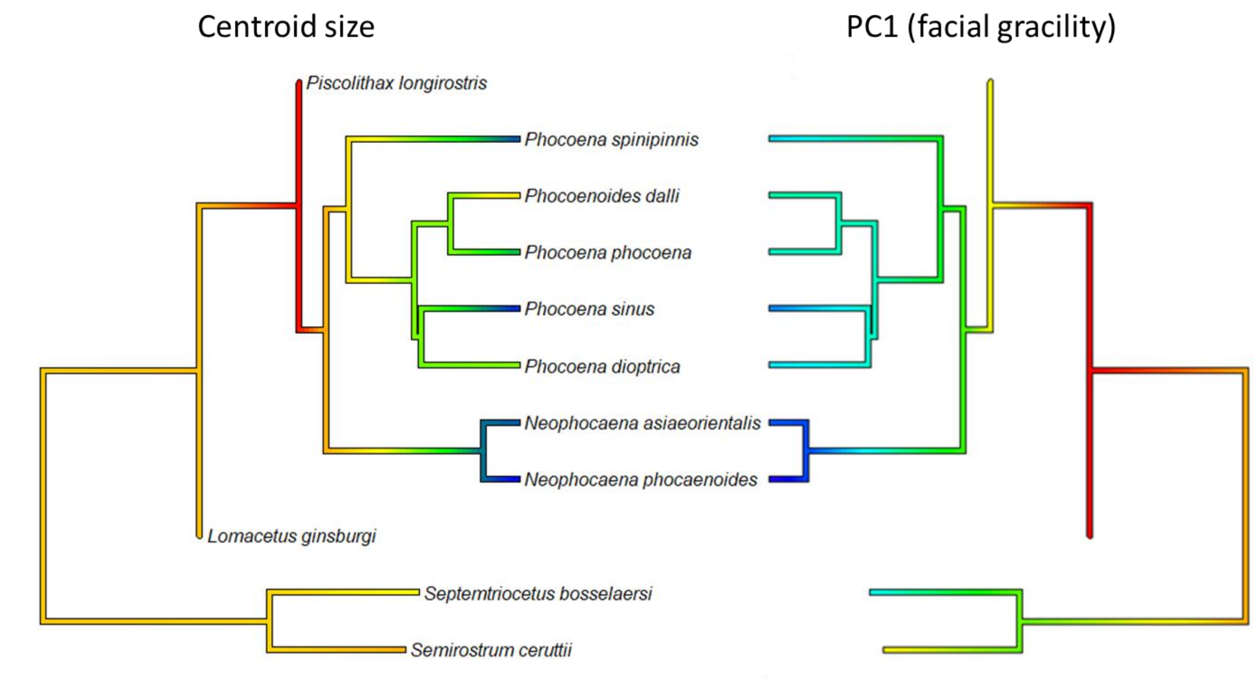

**Figure S33: Phylogeny of the Phocoenidae, coloured on the left by cranial centroid size from smallest (blue) to largest (red), and on the right by PC1 representing relative rostrum length from shortest (blue) to longest (red).**

### Ziphiidae

The Ziphiidae data includes 22 species with a centroid size scaling range of 4.230. There is significant allometry ($R^2$ = 0.129, p=0.014) (Fig. S34A). The predicted shape changes suggest larger species have a smaller braincase, with little change in facial gracility (Fig. S34B). PC1 is not correlated with size (Fig. S34C) but is largely explained by the size of the braincase and length of the rostrum (Fig. S34D). However, PC2 is correlated with size (p<0.001) and represents relative cranial width (Fig S34E). Allometry remains significant after phylogenetic correction ($R^2$=0.144, p=0.012). Larger species are predicted to have smaller braincases and projected premaxillae (Fig. S34F), supporting the pattern after phylogenetic adjustment. There is no phylogenetic signal of centroid size. A comparison between centroid size and facial gracility across the phylogeny shows that, while the smaller species appear to follow the expected pattern, the larger species span the entire range of rostrum length (Fig. S35).

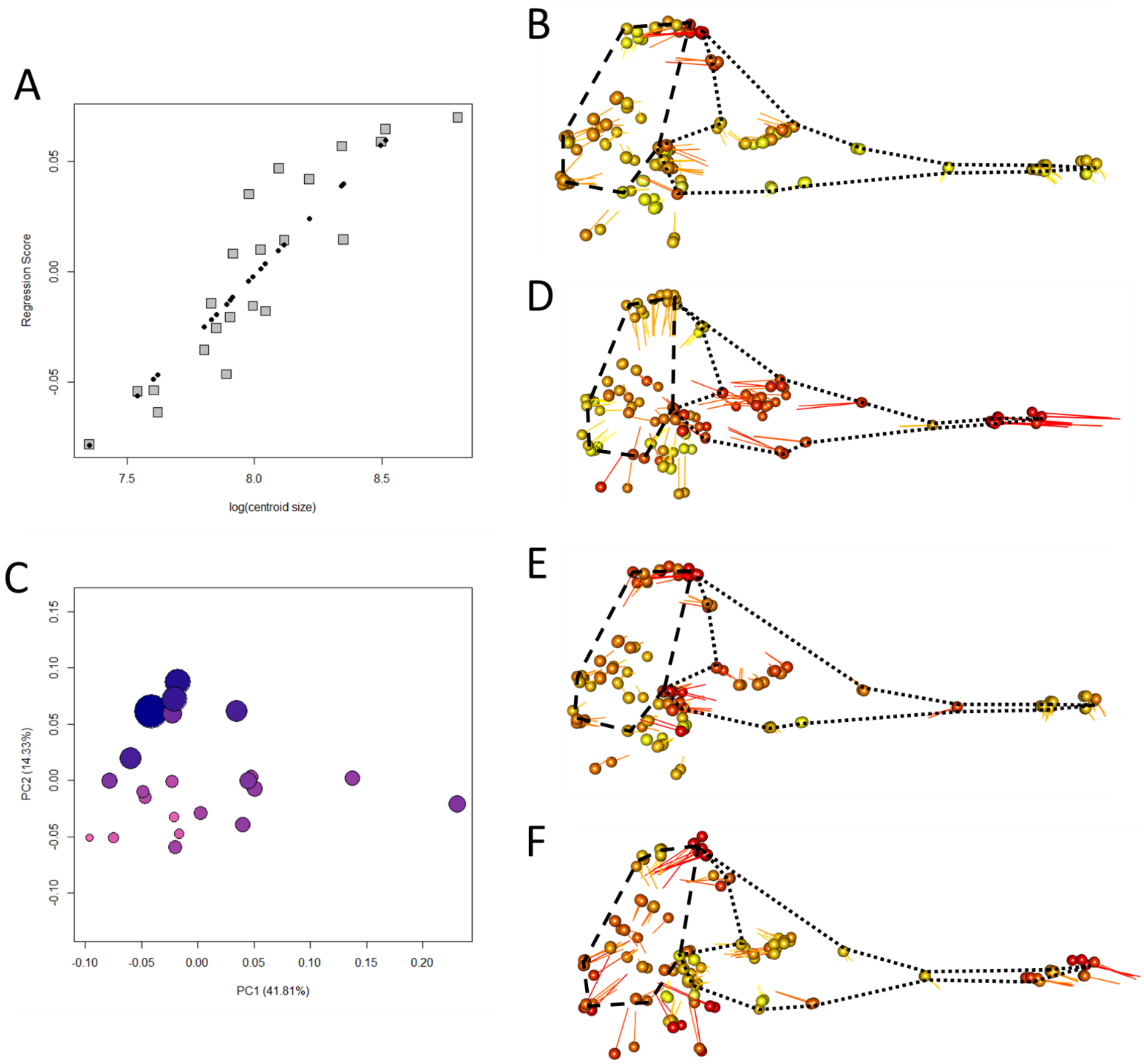

**Figure S34: (A) allometry plot of cranial shape score regressed on log(cranial centroid size), (B) allometric predictions of cranial shape for ordinary least squares, (C) plot of first and second principal components, (D) shape differences across the first principal component, (E) shape differences across the second principal component, (F) allometric predictions of cranial shape for phylogenetic generalised least squares. Dashed lines = braincase, dotted lines = face.**

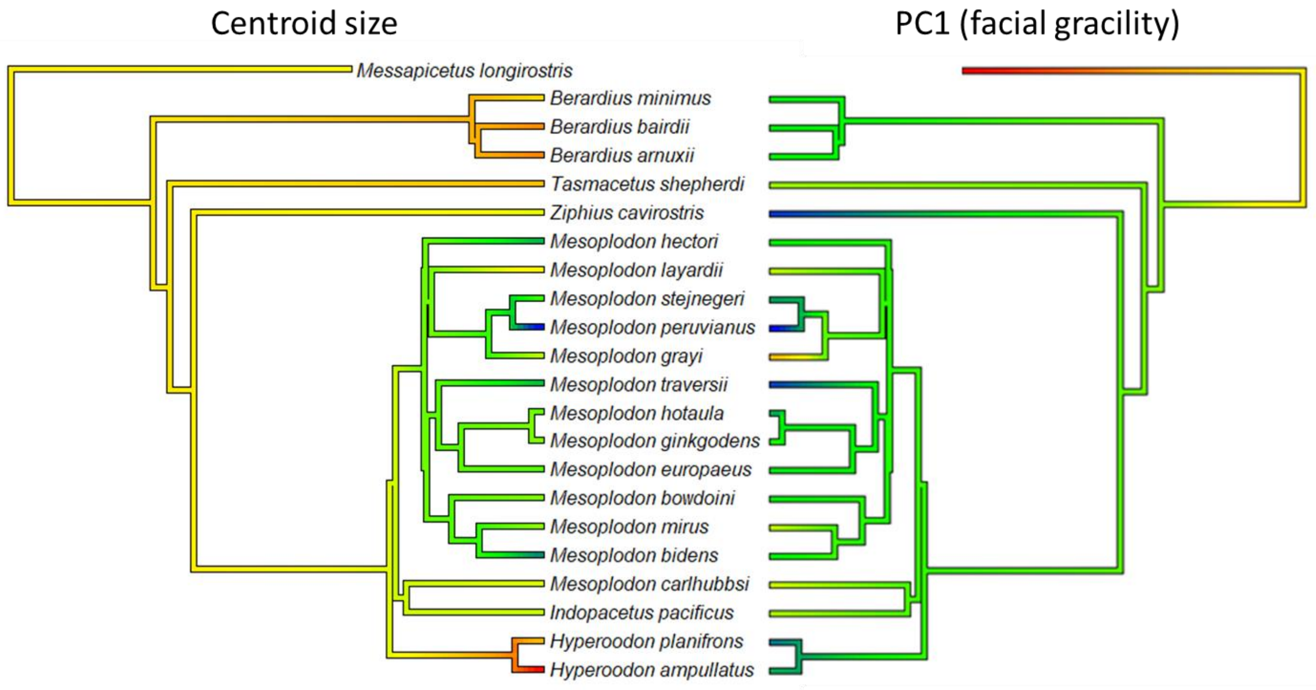

**Figure S35: Phylogeny of the Ziphiidae, coloured on the left by cranial centroid size from smallest (blue) to largest (red), and on the right by PC1 representing relative rostrum length from shortest (blue) to longest (red).**

### Balaenopteridae

The Balaenopteridae data includes 11 species with a centroid size scaling range of 4.169. There is a significant allometric signal ($R^2$ = 0.199, p=0.028) (Fig. S36A). The predicted shape changes suggest larger species have a smaller braincase (Fig. S36B). PC1 is not correlated with size and includes cranial depth and rostrum length (Fig. S36C,D). PC2 is, however, correlated with size (p=0.012) and represents relative braincase size and skull flatness (Fig S36E). Allometry is not significant after phylogenetic correction, and there is no significant phylogenetic signal of centroid size.

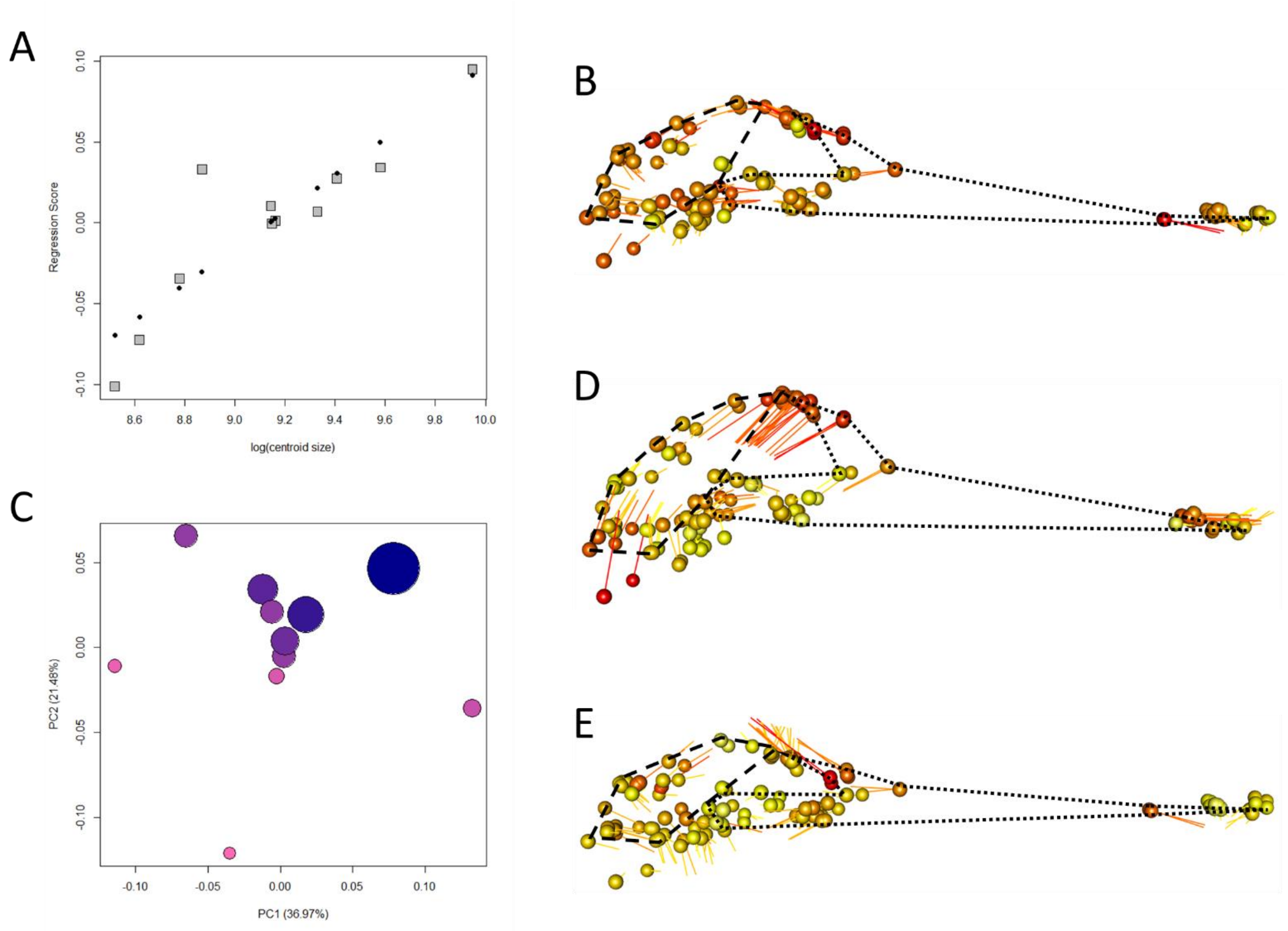

**Figure S36: (A) allometry plot of cranial shape score regressed on log(cranial centroid size), (B) allometric predictions of cranial shape for ordinary least squares, (C) plot of first and second principal components, (D) shape differences across the first principal component, (E) shape differences across the second principal component. Dashed lines = braincase, dotted lines = face.**

### Further Canidae tests

The phylogeny of Canidae (Fig. S37A) can be broadly separated into three monophyletic groups: One dominated by foxes (*Vulpes* spp.), one dominated by South American foxes (*Lycalopex* spp.), and one containing wolves, dogs, and coyotes (*Canis* spp.). Evolutionary allometry is evident in the *Vulpes* clade alone ($R^2$=0.220, p=0.044), and the predicted shape change with increasing size suggests that larger species within the *Vulpes* clade have a more gracile cranium because it is narrower and has projected premaxillae (Fig. S37B). There is no allometry found in the *Canis* clade. The PCA (Fig. S37C) indicates that PC1 defines relative cranial width (Fig. S37D) and separates the large prey specialists from the small prey specialists. Since the large prey specialists occupy both size extremes, this explains the lack of facial elongation predicted with increased size, as the species with the most gracile crania occupy the mid-range of cranial sizes. The bush dog (*Speothos venaticus*) within the *Canis* clade is larger than most species of the *Vulpes* clade (Fig. S37A) and yet has the most robust cranium in the entire Canidae family, in defining PC2 (Fig. S37C) which represents a further anterior shift of the masseter muscles. The bush dog is the smallest member of the *Canis* clade, with the largest being wolf (*Canis lupus*), which has a more stout cranium than the small prey specialists, but more gracile than the smaller large prey specialists. Therefore, while the smaller *Vulpes* species might appear at a glance to be at odds with predictions, in being "longer faced" and smaller than the more robust species, hyperallometric gracilisation can be found in both the *Vulpes* branch and the *Canis* clade when observed in isolation, and accounting for diet mode. When the three smaller large prey specialists of the *Canis* clade are combined with the *Vulpes* clade, allometry is again significant ($R^2$ = 0.175, p=0.044), but in this case facial stoutness is predicted with increased size. This suggests that if larger species switch to more demanding diets, predictions might be reversed from hyperallometric gracilisation.

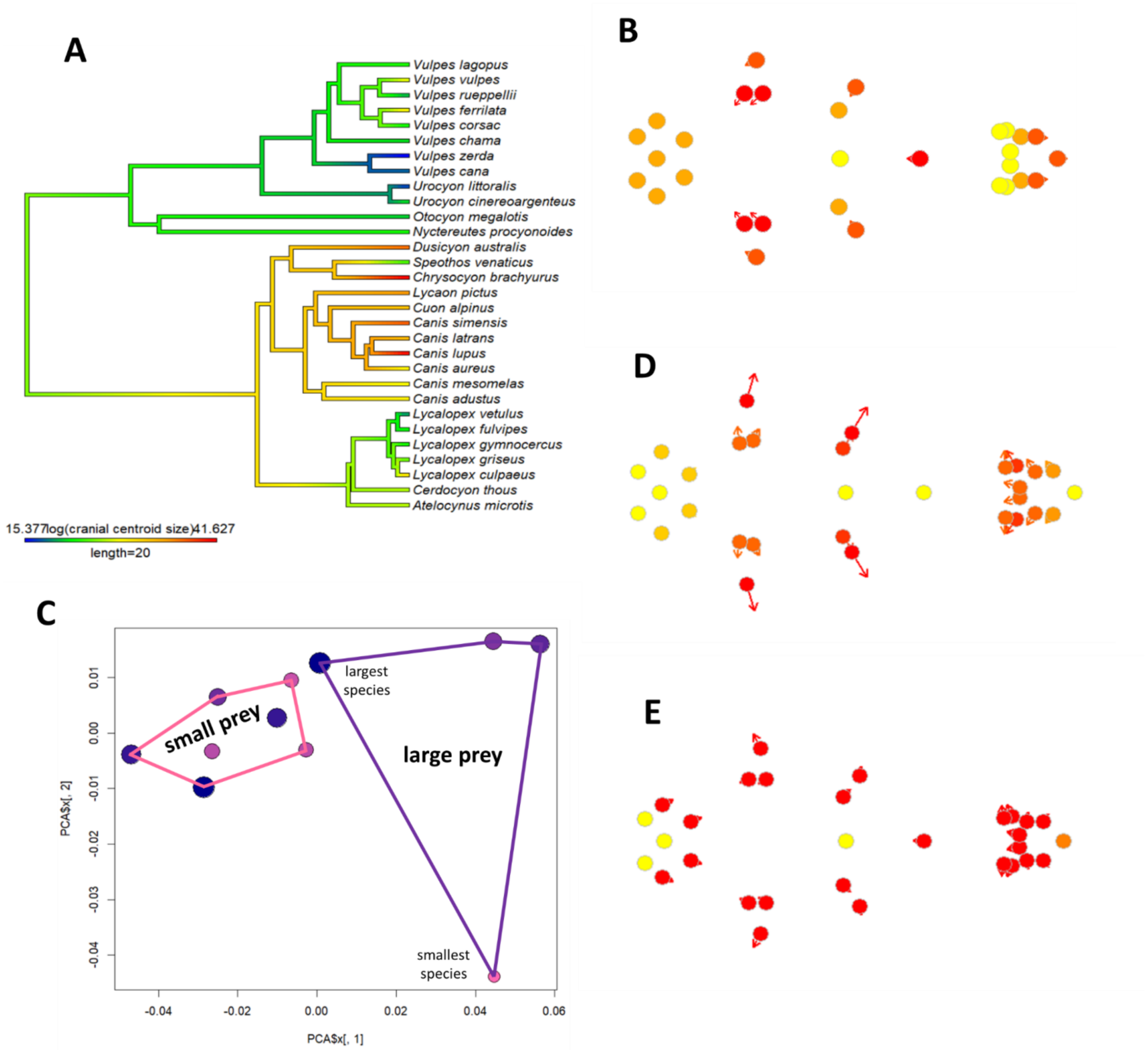

**Figure S37: (A) The phylogeny of the Canidae coloured by cranial centroid size, (B) allometric prediction for the *Vulpes* clade with points representing small crania and arrow tips representing large crania (C) PCA of the *Canis* clade with points representing PC1 minimum and arrow tips representing PC1 maximum, (D) shape differences that define PC1 in the *Canis* clade, (E) allometric predictions for the hypothetical clade of *Vulpes* spp. plus the three smaller large prey specialists that occupy the maximum of PC1 in the *Canis* clade.**
