## Appendix B for "Facing the facts: Adaptive trade-offs along body size ranges determine mammalian craniofacial scaling"

We assessed cranial evolutionary allometry across 22 mammalian families (each with n>10 species), including representatives of the artiodactyls (ruminants and whales), carnivores, bats, rodents, rabbits, and marsupials. We did not include domesticated, brachycephalic morphologies, as these often include pathological degrees of facial foreshortening and malocclusion and there are multiple potential pleiotropic and developmental causes for these occurrences unique to the processes of domestication, resulting in morphologies unlikely to be prevalent or persistent under natural selection (see review by Geiger et al 2021).

We considered the Family level as an arbitrary but consistent taxonomic delineation for testing. The family level tends to contain sufficient species diversity for a reasonable sample across diverse mammalian taxa from published research. Focusing on the family level is arbitrary and might not be biologically meaningful in terms of morphological and ecological ranges. However, choosing an arbitrary taxonomic level removes the potential for biased sampling due to prior assessment of what kind of lineage might be suitable for a study. With exception to the marsupial carnivores (Dasyuridae), which we landmarked ourselves following a landmark protocol of fixed landmarks derived from Viacava et al. (2020), all datasets analysed are from previously published data (Kraatz & Sherratt, 2016; Mitchell et al., 2018; Arbour et al., 2019; Marcy et al., 2020; Bibi & Tyler, 2022; Coombs et al., 2020; Meloro & Tamagnini, 2021).

Phylogenetic trees (time-calibrated) were sourced from the publications whose landmarks we used, or generated using TimeTree (Kumar et al., 2022). Landmarks for each family were extracted from larger datasets based on the matching names of the respective phylogenies. Specimens were then reordered within the data array to match the order of the phylogeny. All data is available on Github (https://github.com/DRexMitchell/Mitchell-etal-facial-scaling).

We employed a widely-used Procrustes-based geometric morphometrics approach, which separates each structure into its shape and isometric size (“centroid size”), to examine allometry (Zelditch et al., 2004; Klingenberg 2016). All analyses were carried out in R version 4.2.1 (R Core Team, 2013) using the *geomorph* package version 4.0.5 (Adams & Otárola-Castillo, 2013; Adams et al., 2021). We first performed a Procrustes superimposition on the data using the ‘gpagen’ function to remove variation attributable to differences in size, rotation, and translation (Rohlf & Slice, 1990). For datasets with more than one specimen per species, cranium shape was characterised by species-averaged values of landmark coordinates aligned with generalised Procrustes analysis. Cranium centroid size was calculated as the square root of the sum of squared distances of each landmark to the centroid (the average coordinates of the landmark configuration; Zelditch, 2004). This value was used as a proxy for body size (e.g., Hood, 2000; Cardini & Polly, 2013; Marcy et al., 2020).

Regarding sexual dimorphism (SD), the Bovidae dataset addressed SD by focussing on males. The Carnivora data found SD to have a negligible effect, using a subsample of 50 species with sufficient sexed individuals. The Cetacea dataset could not consider SD due to a lack of information for these specimens. The Macropodidae are known to show little SD in shape, with most differences relating to size. This is similar for rodents, leporids, and marsupial carnivores. The broad shape variation among bats means that SD was not considered in the initial publication.

**Table S2: Sample details. "n" = sample size, "dim" = dimensions of landmarks, "lmks" = number of landmarks.**

| Family | n | dim | lmks |
|---|---|---|---|
| Dasyuridae | 16 | 3D | 50 |
| Macropodidae | 12 | 3D | 32 |
| Leporidae | 20 | 3D | 52 |
| Muridae | 37 | 3D | 325 |
| Pteropodidae | 24 | 3D | 69 |
| Emballonuridae | 18 | 3D | 69 |
| Rhinolophidae | 28 | 3D | 69 |
| Phyllostomidae | 30 | 3D | 69 |
| Molossidae | 13 | 3D | 69 |
| Vespertilionidae | 29 | 3D | 69 |
| Felidae | 32 | 2D | 30 |
| Viverridae | 15 | 2D | 30 |
| Herpestidae | 17 | 2D | 30 |
| Canidae | 30 | 2D | 30 |
| Mustelidae | 34 | 2D | 30 |
| Phocidae | 15 | 2D | 30 |
| Otariidae | 12 | 2D | 30 |
| Bovidae | 88 | 3D | 53 |
| Delphinidae | 38 | 3D | 123 |
| Phocoenidae | 11 | 3D | 123 |
| Ziphiidae | 22 | 3D | 123 |
| Balaenopteridae | 11 | 3D | 123 |

For each family, evolutionary allometry was assessed using the model, shape ~ log(centroid size) with an ordinary least-squares (OLS) multivariate regression (e.g., Monteiro 1999) via the 'procD.lm' function. This was plotted to depict the regression. Alongside this, we used the 'shape.predictor' function to predict the shape changes along the allometric range (Drake & Klingenberg, 2008; Adams & Nistri, 2010; Hennekam et al., 2020) and visualised the predicted size-related differences between the smallest and largest species using the *LandVR* R package version 0.5.2 (Guillerme & Weisbecker, 2019).

To examine the relationships between evolutionary allometry and facial gracility, we ran a principal component analysis (PCA) of shape on the Procrustes coordinates. PCA is an ordination method that reduces the dimensionality of the landmark coordinate data, an important step in geometric morphometrics to identify the principal axes of variation (Dryden and Mardia, 1998). If allometry and facial gracility are correlated, shape variation contributed by each should be partitioned into the same principal component (PC). To identify which PCs were associated with cranial size, we tested the correlations between each of the first three PCs, representing the majority (usually >65%) of total shape variation, and the natural logarithm of cranial centroid size using a simple linear model with the 'lm' function. Since facial length is often a highly influential region in the analysis of cranial shape, and elongation is often expressed by PC1 for most shape data and taxa (e.g., Mitchell et al., 2018; van der Geer et al., 2018; Arbour et al., 2019; Weisbecker et al., 2020; Bibi & Tyler, 2022;

Meloro & Tamagnini, 2022), we accepted PC3 as a reasonable cut-off for assessing relationships between facial elongation and size. We visualised shape changes along the PCs that described allometry and/or facial gracility using *LandVR*.

We then tested for evolutionary allometry while accounting for phylogenetic non-independence, using a phylogenetic generalised least squares regression for high-dimensional data (PGLS; Adams 2014) with the 'procD.pgls' function. This tested the same model as the OLS but also adjusted for phylogenetic relatedness between species. Where PGLS analyses were significant, we then again visualised the predicted size-related differences between the smallest and largest sizes using *LandVR*. Note that PGLS has some caveats; for example, it assumes Brownian Motion as the only evolutionary model so that accurate estimates of evolutionary models in comparative analyses are unlikely, particularly when the number of species is modest (Tamagnini et al., 2017). We acknowledge that across 44 independent tests, there is also a possibility of inflating Type 1 errors. In addition, PGLS cannot consider uncertainties in the phylogeny. However, PGLS is still the most widely – and successfully - used method for incorporating phylogeny in GMM studies, so that we follow these common protocols here.

Finally, we assessed the phylogenetic signal in centroid size using the K statistic (Blomberg et al. 2003) in order to see if there is a phylogenetic pattern to species cranial size across each family, which might affect phylogenetic adjustments to data.

Importantly, shape predictions for each test were generated via estimates of shape scores along the regression line. However, a regression line for the sample can be computed whether the regression is significant (i.e. likely biologically meaningful) or not. Therefore, we only accepted evidence of CREA when predicted facial gracility in larger species was accompanied by significant support (at $p \leq 0.05$) for allometry from the model of shape~log(cranial size). As outlined in the main manuscript, gracility is here defined relatively broadly to include one or more of the following: projection of the rostrum (usually involving elongation of the maxillae/premaxillae), retraction of the zygomatic arches, or a more gracile (narrower/flatter) cranium in larger species.

For some families with particularly relevant combinations of shape and size, we visually presented phylogenetic distributions of cranial centroid size and/or facial gracilisation via relevant PCs using the 'contMap' function in the *phytools* package version 1.0-3 (Revell, 2012). In the main manuscript, we used this method to highlight the allometric patterns in the Canidae (Figure 6).

Results of all of these analyses for each family are presented in the Appendix C. However, to be concise, we only present visualisations for shape predictions of significant evolutionary allometry or facial gracility.

For the main manuscript, to observe any associations between centroid sizes across Families and a tendency for facial elongation, we also plotted centroid size ranges across all families tested. However, since the Carnivora landmarks were 2D, we converted these to an approximation of a 3D centroid size by cubing the root of the 2D centroid size, i.e.:

$$CS_{3D} = \sqrt{CS_{2D}}^{3}$$

Cranial meshes used in figures were obtained from Morphosource (https://www.morphosource.org/) under the following codes: *Felis margari*ta (000110512),

*Panthera tigris* (000123442), *Vulpes zerda* (000371195), *Vulpes vulpes* (000116031), *Canis simensis* (000115948), *Canis lupus* (000117236), *Speothos venaticus* (000116025), *Mustela nivalis* (000420959), *Enhydra lutris* (000121164), and *Arctonyx collaris* was obtained from Digimorph (http://www.digimorph.org/specimens/Arctonyx_collaris).
